# Targeted DNA transposition using a dCas9-transposase fusion protein

**DOI:** 10.1101/571653

**Authors:** Shivam Bhatt, Ronald Chalmers

## Abstract

Homology directed genome engineering is limited by transgene size. Although DNA transposons are more efficient with large transgenes, random integrations are potentially mutagenic. Catalytically inactive Cas9 is attractive candidate for targeting a transposase fusion-protein because of its high specificity and affinity for its binding site. Here we demonstrate efficient Cas9 targeting of a mariner transposon. Targeted integrations were tightly constrained at two adjacent TA dinucleotides about 20 bp to one side of the gRNA binding site. Biochemical analysis of the nucleoprotein complexes demonstrated that the transposase and Cas9 moieties of the fusion protein can bind their respective substrates independently. In the presence of the Cas9 target DNA, kinetic analysis revealed a delay between first and second strand cleavage at the transposon end. This step involves a significant conformational change that may be hindered by the properties of the interdomainal linker. Otherwise, the transposase behaved normally and was proficient for integration in vitro and in vivo.

## INTRODUCTION

Biotechnology and medicine are increasingly reliant on our ability to engineer the mammalian genome. Lentiviruses are useful because gene-delivery across the cell membrane is efficient and integration of the viral genome provides long-term transgene expression. However, they are mutagenic because they integrate at random sites using a mechanism similar to the cut-and-paste transposons. An alternative strategy for ex vivo applications is to establish genomic safe-havens for site-specific recombinases such as the phage C31 integrase or the Cre recombinase (Branda and Dymecki, 2004; Song and Palmiter, 2018). This approach lacks flexibility because it is limited to transgene integration at a predetermined site.

Scar-less engineering of a target site must rely on the cells homologous recombination machinery. Although desired modifications can be made very precisely, the events are rare and difficult to recover. One way to boost the rate of recombination by orders of magnitude is to make a DNA double strand break at the target site (Urnov et al., 2005). This can be achieved using zinc finger nucleases and transcription activator-like effector (TALE) nucleases, which can themselves be engineered to target a user-defined site (Chandrasegaran and Smith, 1999; Zhang et al., 2011). Two significant problems are the extended lead-times and off-target cleavages. TALE and zinc finger nucleases have now been largely superseded by the Cas9 nuclease. It can be programed to target a 20 bp recognition sequence, simply by providing a matching guide RNA (gRNA). Off-target cleavage is relatively low and the long recognition sequence provides ample specificity. However, homologous recombination remains a limiting factor because the efficiency falls off as the transgene size increases (Li et al., 2014).

Transposon vectors are widely used for gene delivery applications (Grabundzija et al., 2010; Lamberg et al., 2002; Liang et al., 2009; Mates et al., 2009; Way et al., 1984). Although their efficiency falls off as the cargo size is increased, they are less sensitive to this parameter than host-mediated homologous recombination (Balciunas et al., 2006; Claeys Bouuaert et al., 2013; Ding et al., 2005; Izsvak et al., 2000; Lohe and Hartl, 1996). However, like the Lentiviruses, transposon vectors are mutagenic because the integration sites are essentially random.

To combat random integration various groups have used site-specific DNA binding proteins to target specific loci. Examples include Mos1, Sleeping Beauty, piggyBac and ISY100, which were variously fused to zinc finger proteins, TALEs and Gal4 (Feng et al., 2010; Ivics et al., 2007; Luo et al., 2017; Maragathavally et al., 2006; Owens et al., 2013; Yant et al., 2007). Although successful, targeting strategies have all me with significant problem: namely, that the transposases are still capable of random integration. Since, DNA binding proteins, such as zinc fingers and TALEs, spend most of their time searching for their binding sites, targeted integrations are recovered against a background of random events. To tackle this problem we are currently developing a switchable transposase, which allows control over the timing of integration. In parallel, we were considering the advantages of potential targeting moieties. Cas9 is an obvious and attractive candidate because extensive base pairing with the target provides a dwell time of several hours (Ma et al., 2016). In contrast, DNA binding proteins remain bound to their specific sites only for a matter of seconds or minutes.

Previously, there was an unsuccessful attempt to use a catalytically inactive Cas9 (dCas9) to target piggyBac insertions to a hypoxanthine phosphoribosyl transferase (HPRT) gene in human cells (Luo et al., 2017). Surprisingly, instead of targeting the HPRT locus, the dCas9-piggyBac chimera protected it from insertions. We therefore set out to test whether Cas9 is a general inhibitor of transposition or whether the effect is peculiar to piggyBac. The choice of the transposase moiety is limited by several factors. Although Tn5 is very active in vitro, and is used extensively in bacteria, it transposes poorly in mammalian cells (Blundell-Hunter et al., 2018). Sleeping Beauty is efficient in vivo but lacks an in vitro system (Mates et al., 2009). While piggyBac is probably the most efficient for gene delivery in vivo, it has a poor in vitro system (Liang et al., 2009; Mitra et al., 2008), which precludes the development of advanced applications such as direct delivery of transpososomes.

Our model system is based on Hsmar1, a reconstituted mariner-family transposon (Miskey et al., 2007; Robertson and Zumpano, 1997). Although it is not as active as piggyBac or Sleeping Beauty in cell culture transfection experiments, the reaction is 100% efficient in vitro (Claeys Bouuaert and Chalmers, 2017). Furthermore, if the mariner transpososome is assembled in vitro, it can be delivered cells directly (Trubitsyna et al., 2017). Hsmar1 also has much lower non-specific nuclease activity than other mariner elements such as Mos1, Mboumar1 and Himar1 (Claeys Bouuaert and Chalmers, 2010; Dawson and Finnegan, 2003; Lampe et al., 1996; Lipkow et al., 2004; Munoz-Lopez et al., 2008; Trubitsyna et al., 2015).

## MATERIALS AND METHODS

DNA oligonucleotides and most dry chemicals were from Sigma Aldrich. Enzymes were from New England Biolabs and DNA purification kits were from Qiagen. The nucleotide sequences of all plasmids used in this work are given in Supplemental Table 1. The β-galactosidase assays were as described by Miller (Miller, 1972). LB agar indicator plates contained 40 µg/ml X-gal, when present,.

For in vivo assays, dCas9 was expressed from derivatives of plasmid pdCas9 (Bikard et al., 2013). When present, CRISPR spacers were cloned into the BsaI site as oligoduplexes (Supplemental Table 2). The Hsmar1 transposase gene was added to the 5’- or 3’-end of the dCas9 gene by PCR using pdCas9 and pRC880 as templates: transposase-dCas9, pRC2302; dCas9-transposases, pRC2303. An oligoduplex encoding spacer-7 was cloned into the CRISPR locus of these plasmids to produce pRC2304 and pRC2305, respectively. The native wild type transposase was expressed from pRC2306, which is identical to pdCas9 except that the dCas9 gene is replaced by the transposase gene.

The transcription-strength series of expression vectors for Figure 2C, D was created using the constitutive promoters described by Tellier and Chalmers (Tellier and Chalmers, 2018). In the plasmid with the strongest promoter, P4 (pRC2309), the native S. pyogenes Cas9 promoter in pdCas9 was replaced by Tellier-P2. In plasmid P3 (pRC2308) the S. pyogenes promoter was replaced by Tellier-P1. In plasmid P2 (pRC2307) the S. pyogenes promoter was deleted by inverse PCR and residual transcription was from a cryptic promoter(s). The plasmid with the lowest level of transcription, P1 (pRC2311), was created by replacing the multi-copy origin-of-replication in plasmid P3 (above) with a single-copy origin-of-replication from a bacterial artificial chromosome vector.

Hsmar1 transposase was expressed from pRC880 and purified as a maltose binding protein (MBP) fusion by affinity chromatography as described (Claeys Bouuaert and Chalmers, 2010; Claeys Bouuaert et al., 2011). The expression vector for the dCas9-transposase fusion protein was pRC2303 described above.

The dCas9-transposase fusion-protein was expressed from the constitutive promoter in pRC2303. The plasmid was transformed into E. coli NiCo21 (New England Biolabs). A single colony was picked into a starter culture of LB-Lennox supplemented with 30 µg/mL chloramphenicol and grown at 37°C for 16 hours with shaking at 250 rpm in a baffled Erlenmeyer flask. The culture was diluted 1:100 into fresh media and growth was continued until the OD600 reached 0.5. The temperature was then reduced to 18°C and the incubation was continued for 16 hours. Cell (500 ml) were harvested by centrifugation at 4ºC and resuspended in 40 ml binding buffer (20 mM Tris, pH 8.0, 5 mM imidazole, 0.5 M sodium chloride, 2 mM DTT). All subsequent steps were carried out at 4ºC. Cells were lysed by five passes through a French pressure cell at 16,000 psi and the extract was clarified by centrifugation at 40,000 x g for 30 minutes. The clarified extract was applied to a 1 mL HisTrap HP column (GE Life Sciences) on an AKTA FPLC (Amersham Pharmacia). The column was equilibrated with binding buffer and loosely bound proteins were eluted with 40 column volumes of wash buffer (20 mM Tris at pH 8.0, 60 mM imidazole, 0.5 M sodium chloride, 2 mM DTT). The column was then developed with a 20 column-volume linear-gradient up to 1 M imidazole. Peak fractions were pooled an analyzed by SDS polyacrylamide gel electrophoresis (Supplemental Figure 1). When required for in vitro reactions, the sgRNA and Cas9-transposase were assembled into a ribonucleoprotein complex by incubating for 20 minutes at room temperature in dCT binding buffer (20 mM HEPES pH 7.5, 250 mM potassium chloride, 2 mM MgCl2, 0.01 % Triton X-100, 0.1 mg/mL BSA, 10% glycerol) (Anders et al., 2015). This was done every time it was required immediately before it was used.

The EMSA binding buffers for the native transposase and the dCas9-transposase were, respectively, [20 mM HEPES at pH 7.5, 100 mM NaCl, 2 mM DTT, 10 % glycerol, 5 % DMSO, 5 mM CaCl2, 250 µg/ml BSA] and [20 mM HEPES pH 7.5, 250 mM potassium chloride, 2 mM MgCl2, 0.01 % Triton X-100, 0.1 mg/mL BSA, 10% glycerol]. Reactions (20 µl) contained 10 nM of a Cy-5 labeled oligoduplex encoding either the transposon end or the target of the sgRNA (Supplemental Table 3). After addition of the dCas9-sgRNA-transposase complex, the reactions were incubated for 20 minutes at 37ºC. Binding reactions with the native transposase were incubated for 60 minutes at room temperature to be consistent with our previous work. Binding reactions were loaded onto a TBE buffered 7% polyacrylamide gel and electrophoresed at 120 V. Gels were recorded on a GE Typhoon imager.

In vitro transposition reactions were performed as described (Claeys Bouuaert and Chalmers, 2010). The reaction buffer contained 25 mM Tris-HCl at pH 8.0, 5% glycerol, 100 mM sodium chloride, 2 mM DTT, 2.5 mM MgCl2 and 6.5 nM of supercoiled pRC650 as substrate. Transposition reactions (50 µl) were initiated by the addition of transposase and incubated for 4 h at 37ºC. Reactions were made 0.4% in SDS and 19 mM in EDTA and heat inactivated at 75ºC for 30 minutes. Samples (35 µl) were electrophoresed on a TBE buffered 1.1% agarose gel at 60 V for 16 – 24 h. Gels were stained with ethidium bromide and photographed. In vivo transposition reactions were performed as described (Liu and Chalmers, 2014). The reporter strain [E. coli RC5096, F-fhuA2 Δ(lacZ)r1 glnV44 e14-(McrA-) trp-31 his-1 rpsL104 xyl-7 mtl-2 metB1 Δ(mcrC-mrr)114::IS10 argE::Hsmar1-lacZ’-kanR] was derived from ER1793 (New England Biolabs).

Targeted transposition experiments were based on a well known ‘plasmid-hop’ assay eg. (Munoz-Lopez et al., 2008). The transposon donor plasmid was pRC704, which encodes an R6K origin of replication and a mini-Hsmar1 transposon with a kanamycin resistance gene. The small target plasmid was pRC2312, which is essentially a dimer of pBluescript II SK+ with one of the copies of lacZα removed. The large target was pRC2313, which is identical to the small target except that a 4.5 kb fragment of non-specific DNA from the E. coli chromosomal argE-H locus was inserted. Reactions (15 µl) were performed in our standard transposition buffer (25 mM Tris-HCl at pH 8.0, 5% glycerol, 100 mM sodium chloride, 2 mM DTT, 2.5 mM MgCl2) with 9 nM of target plasmid and 13.5 nM pRC1105 to provide a background of non-specific DNA. Reactions were incubated for 20 minutes at 37ºC, initiated by the addition of the transposon donor plasmid (9 nM) and incubated for a further 24 hours. Reactions were brought to 50 µl with NEB Cutsmart buffer, supplemented with 5 units of NheI and 5 units of lambda exonuclease to degrade the non-specific plasmid. Digestion was stopped by adding 15 µl of stop solution (0.5 mg/mL proteinase K, 0.1% SDS, 50 mM EDTA) and incubating at 60ºC for 2 h. DNA was purified using a spin column from the Qiagen PCR Clean-up kit, which was eluted with 30 µl of EB (10 mM Tris-HCl, pH 8.5). 3-10 µl of the reaction was transformed into chemically competent E. coli NEB5α and plated on LB agar plates supplemented with 50 µg/ml kanamycin, 100 µg/ml ampicillin, 40 µg/ml X-gal and 0.1% lactose. Targeting efficiency (%) was calculated by dividing the number of white colonies by the total number of colonies and multiplying by 100. Target plasmids from randomly selected white colonies were purified and the location of the transposon insertion in lacZα was determined by Sanger sequencing using the M13 reverse primer.

## RESULTS

### Selection of gRNAs and construction of transposase-dCas9 fusions

Since gRNAs do not all bind their targets equally well (Liu et al., 2016) we screened ten candidates for their ability to inhibit transcription of the target lacZ gene. Oligoduplex spacer sequences, encoding candidate gRNAs, were ligated into the CRISPR locus of plasmid pdCas9 (Figure 1A). Subsequent transcription and processing in vivo produces a mature gRNA, which forms a ribonucleoprotein complex with dCas9. The relative locations and orientations of the selected gRNA target-sequences are shown in Figure 1B. *E. coli* BL21 cells were transformed with each dCas9-gRNA expression plasmids separately and spread on X-gal indictor plates to test the ability of the gRNAs to inhibit transcription of *lacZ*. Only spacers 4 and 7 produced completely white colonies (not shown). Because spacer-7 is within the coding sequence of the gene, it was selected for all further experiments. To create a fusion protein capable of targeted transposition, the Hsmar1 transposase gene was added to the 5’- or the 3’-end of the dCas9 gene in pdCas9 (Figure 1A). Two versions of the fusion protein expression vectors were created, one with and one without spacer-7.

**Figure 1.**
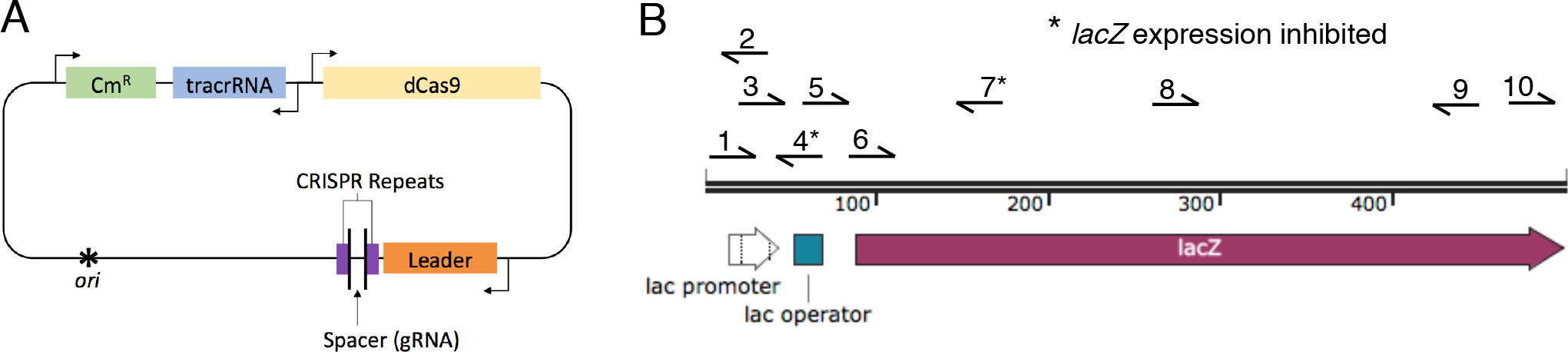
Plasmid pdCas9 and the chromosomal *lacZ* target. **A)** Plasmid pdCas9 (Bikard et al., 2013) encodes a catalytically inactive Cas9 variant (dCas9) in which the nuclease activity is abolished by the D10A and H840A mutations. Expression is driven by the strong native Cas9 promoter from Streptococcus pyogenes. The spacer is the user-defined sequence that base pairs with the target site. **B)** The target site for dCas9 binding was the *E. coli lacZ* gene. Ten candidate spacers were selected to target the promoter and early open reading frame of *lacZ.* Of these 10 candidates, only spacers 4 and 7 inhibited transcription completely, as judged by a white colony on X-gal indicator plates.

### The fusion-proteins are active in vivo

To test whether the fusion proteins were still capable of gRNA-mediated transcriptional repression, the four plasmids were transformed into E. coli BL21 and the lacZ activity was measured using the Miller assay (Figure 2A). There was little silencing with dCas9 or either of the fusion proteins in the absence of the gRNA. In the presence of the gRNA the N- and C-terminal dCas9 fusion proteins both repressed lacZ expression effectively. We then used a bacterial transposition assay to test whether the transposase domain retained activity (Figure 2B). We also tested a plasmid in which the dCas9 gene in pdCas9 was substituted with the transposase gene. This provided a positive control in which transposase expression was driven by the same promoter as the fusion proteins. The rate of transposition was highest for the transposase-dCas9 plasmid, which has the transposase fused to the N-terminus of Cas9. This result should be interpreted with caution as Hsmar1 transposition is subject to over-production inhibition (OPI) whereby an increase in the transposase concentration beyond a low level decreases the rate of transposition (Claeys Bouuaert et al., 2013; Tellier and Chalmers, 2018). Since the promoter on pdCas9 is relatively strong, the system will be dominated by OPI. In the presence of the gRNA the rate of transposition for the N- and C-terminal fusion proteins decreased significantly (Figure 2B). If this result is interpreted in terms of the OPI model, it indicates that the gRNA increased the transpositional-potential of the system, which decreased the rate of transposition on account of OPI. Since the dCas9-transposase protein, which has the transposase on the C-terminus of dCas9, had the best transpositional-potential and the best transcriptional repression in the Miller assay, it was selected for all further experiments (Figure 2A, B).

**Figure 2.**
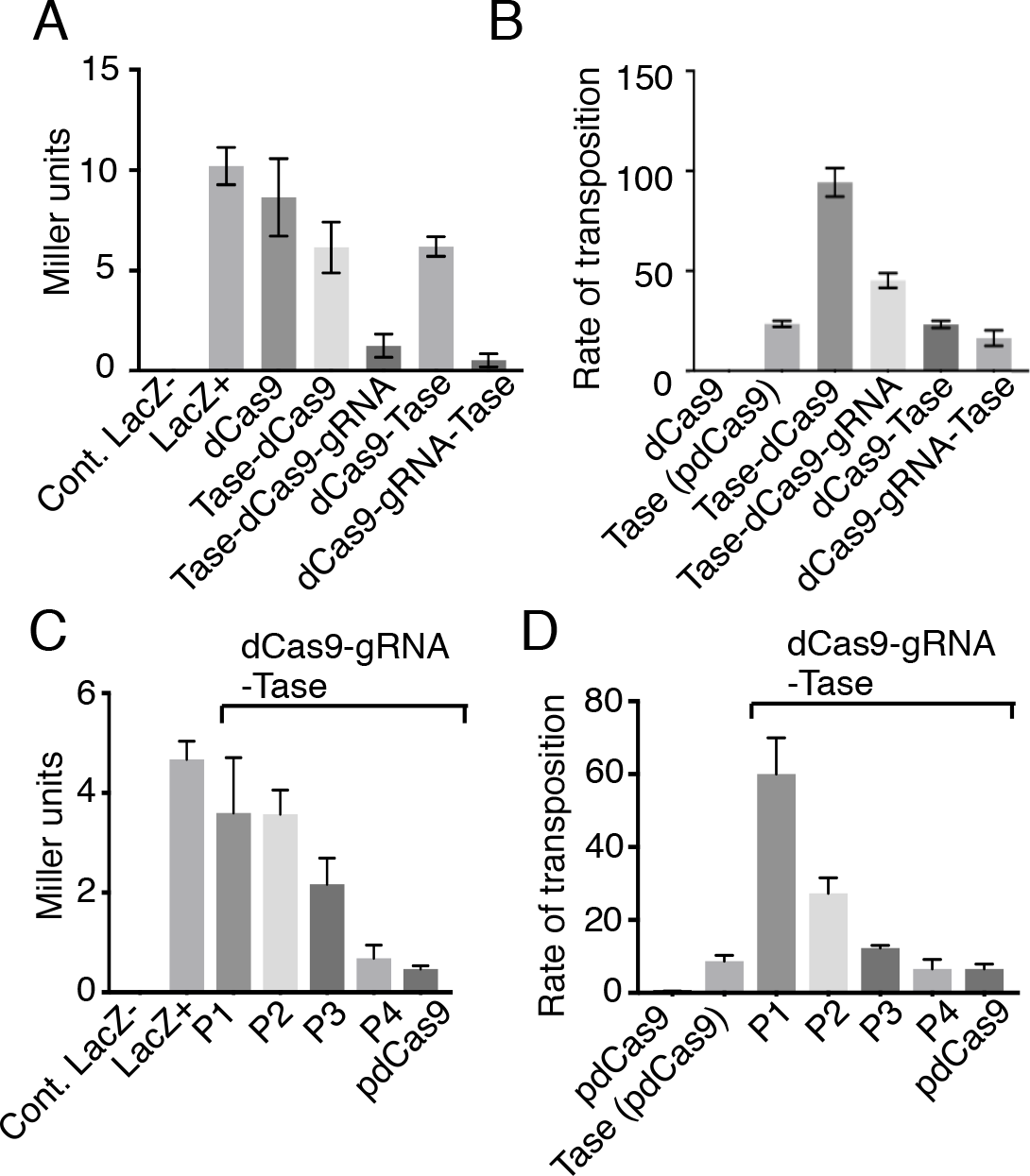
Fusion-protein activity in vivo. The Hsmar1 transposase gene was added to the 5’- or the 3’-end of the dCas9 gene in pdCas9, which is illustrated in Figure 1A. **A**, The plasmids were transformed into E. coli BL21 and lacZ activity was measured using the Miller assay (Miller, 1972). The lacZ-control was provided by E. coli NEB5α, while LacZ+ is the activity with the untransformed BL21 host. Proteins were expressed from plasmids: dCas9, pdCas9 = pRC2301; transposase-dCas9, pRC2302; transposase-dCas9-gRNA, pRC2304; dCas9-transposase, pRC2303; dCas9-gRNA-transposase, pRC2305. **B**, The relative rates of transposition supported by the N- and C-terminal fusions was assayed in E. coli as described previously (Liu and Chalmers, 2014). The wild type transposase was expressed from pRC2306, which is identical to pdCas9 except that the dCas9 gene is replaced by the transposase gene. Other protein expression vectors were as in part B. **C**, The relationship between lacZ repression and the fusion-protein expression level was explored. The highest level of expression was with the native Cas9 promoter form pdCas9. Promoters P4 to P1 were progressively weaker (Tellier and Chalmers, 2018). The expression vectors were: P1, pRC2311; P2, pRC2307; P3, pRC2308; P4, pRC2309. **D**, The relationship between the expression level and the rate of transposition was explored. Owing to auto-inhibition (or OPI) the highest rate of transposition was with the lowest expression level, as expected (Claeys Bouuaert et al., 2013). Protein expression vectors were as in previous parts.

To explore the relationships between the expression level of the fusion protein and its activities, the native Cas9 promoter in pdCas9 was replaced with promoters P4 to P1, which are of progressively lower strength (Tellier and Chalmers, 2018). As expected, transcriptional repression by the fusion protein was highest with the native Cas9 promoter and decreased progressively with promoter strength (Figure 2C). In contrast, the rate of transposition had an inverse relationship to the strength of the promoter, consistent with the expected effects of OPI mentioned above (Claeys Bouuaert et al., 2013).

### dCas9-transposase binds transposon ends and the gRNA target

The assembly pathway for the Hsmar1 transpososome and its behavior in an electrophoretic mobility shift assay (EMSA) is illustrated in Figure 3A, B. Binding of the first transposon end to the transposase dimer yields single-end complex 2 (SEC2). Recruitment of a second end yields the paired-ends complex (PEC). This would normally be followed by the first cleavage step of the reaction, except that the binding buffer lacks the catalytic magnesium ion. During electrophoresis, the PEC decays into single-end complex 1 (SEC1) (Claeys Bouuaert et al., 2013). When the purified dCas9-transposase is titrated into the reaction the disappearance of the free transposon ends is accompanied by the appearance of two clear shifted bands plus a third band close to the bottom of the well (Figure 3C). This is broadly similar to the behavior of the wild type transposase except that the complexes are further up the gel owing to the much larger size of the fusion protein. The band that just enters the gel above SEC2 might represent the PEC, which is detected in some experiments with the wild type transposase (Claeys Bouuaert et al., 2013).

**Figure 3.**
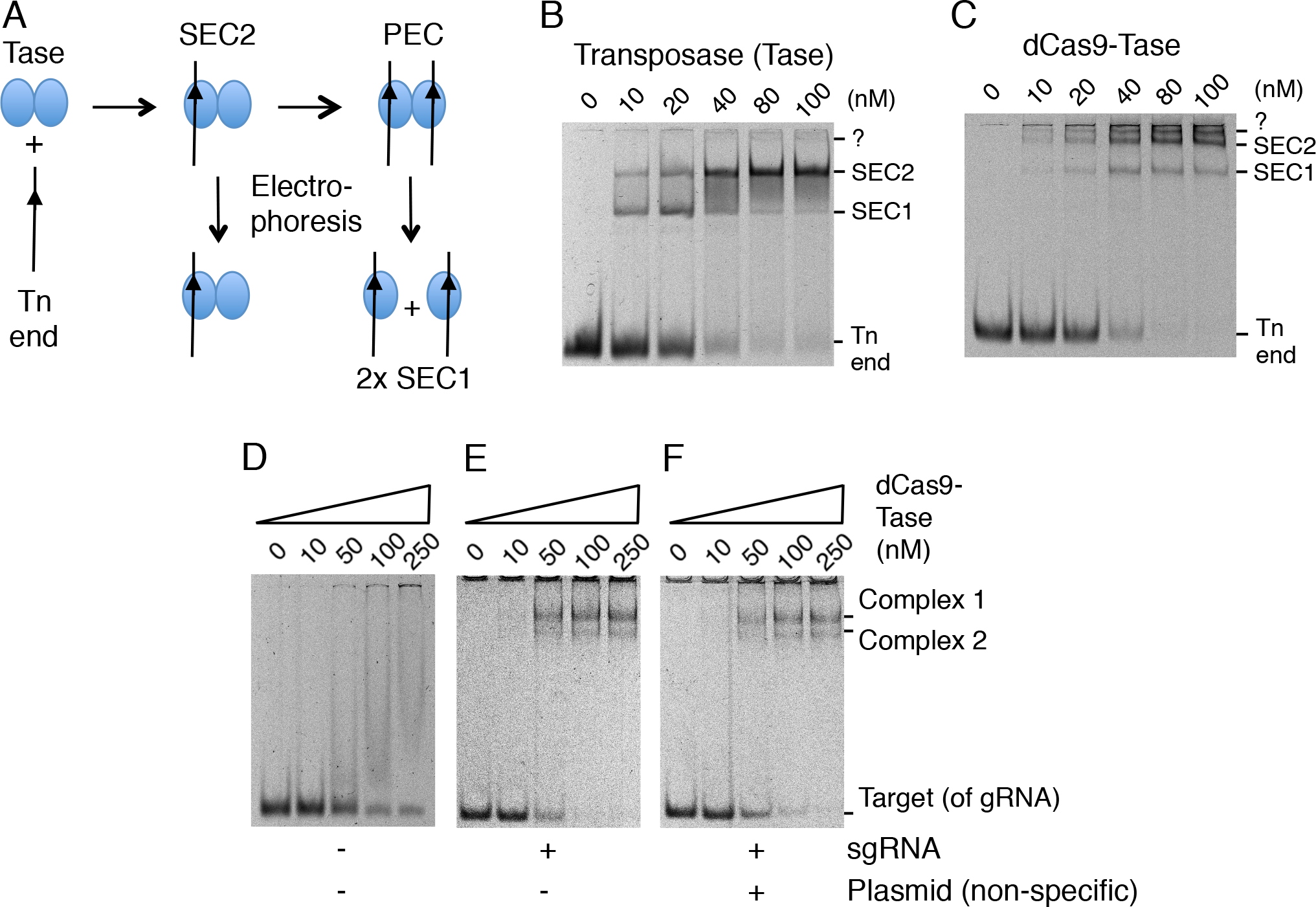
EMSA analysis of the dCas9-transposase fusion protein. **A**, Illustration of protein-DNA complexes formed during the transposition reaction and decomposition of the paired-end complex (PEC) during electrophoresis. **B**, EMSA of purified Hsmar1 transposase on a TBE buffered 7% polyacrylamide gel. Reactions contained 10 nM of a 70 bp Cy5-labelled oligoduplex encoding a transposon-end substrate. Bands were visualized on a GE Typhoon Fluorimager. **C**, EMSA as in part B but with the indicated amounts of the dCas9-transposase fusion protein without sgRNA. The band marked ?, just below the wells, might represent the PEC, which is detected with the wild type transposase if the PEC is artificially stabilized (Claeys Bouuaert et al., 2013). **D**, **E** and **F**, the EMSAs were as in part B but with the purified dCas9-transposase fusion protein with its sgRNA. Reactions contained 10 nM of a 70 bp Cy5-labelled oligoduplex complementary to sgRNA-7. When present, the sgRNA was preassembled in a 1:1.5 molar excess to the protein prior to the EMSA. The non-specific competitor plasmid was 1 *µ*g of pRC2301 per binding reaction.

To explore the binding ability of the dCas9 moiety of the fusion protein, an EMSAs was performed with an oligoduplex complimentary to the spacer-7 sequence (Figure 3D, E, F). Since this experiment was performed in vitro we used a single-guide RNA (sgRNA) in which the tracrRNA and the gRNA were joined in a single strand. In the absence of the sgRNA, titration of the fusion protein produced a smear with no clear bands (Figure 3D). Presumably, this indicates weak interactions between the protein and the DNA, which dissociate during electrophoresis. When the fusion protein was provided with the sgRNA, two clear complexes were formed (Figure 3E). The stoichiometry of the complexes is unknown but they may correspond to a dimer of the fusion protein bound by either one or two target oligoduplexes. When we added unlabeled competitor DNA to the reaction it had not effect, which indicates that the oligoduplex binding to the fusion protein is specific (Figure 3F).

### In vitro transposition with the dCas9-transposase fusion-protein

Since we had demonstrated DNA binding by the respective dCas9 and transposase moieties of the fusion protein we next wanted to examine the intermediates and products of the in vitro transposition reaction. When the wild type transposase is incubated with a supercoiled substrate, excision of the transposon leaves behind the plasmid backbone, which is an end product of the reaction and a convenient measure of the efficiency (Figure 4). After excision, target sites for integration are acquired by a random collision and tracking mechanism (Claeys Bouuaert and Chalmers, 2010). Titration of the reaction with the wild type transposase or dCas9-transposase yielded a very similar range of products (Figure 4). Minor differences were probably owing to the differences in the quality of the purified proteins. However, adding an oligoduplex encoding the target of the sgRNA reduced the amount of protein required for the maximum production of backbone. It also changed the amount and relative proportions of the reaction products. Most noticeable is the accumulation of the relaxed substrate. This is an intermediate of the reaction generated by the first nick at one of the transposon ends (Claeys Bouuaert and Chalmers, 2010; Claeys Bouuaert et al., 2014). The transition between the first nick and the second, which completes cleavage at the transposon end, involves a significant conformational change (Claeys Bouuaert and Chalmers, 2017; Dornan et al., 2015). Since nicking is absolutely dependent on PEC formation (Claeys Bouuaert et al., 2011), the accumulation of the relaxed intermediate shows that the dCas9 moiety of the fusion protein can interact with its oligoduplex target at the same time as the transposase moiety is engage in a transposition reaction. Clearly, this is a prerequisite for successful targeting.

**Figure 4.**
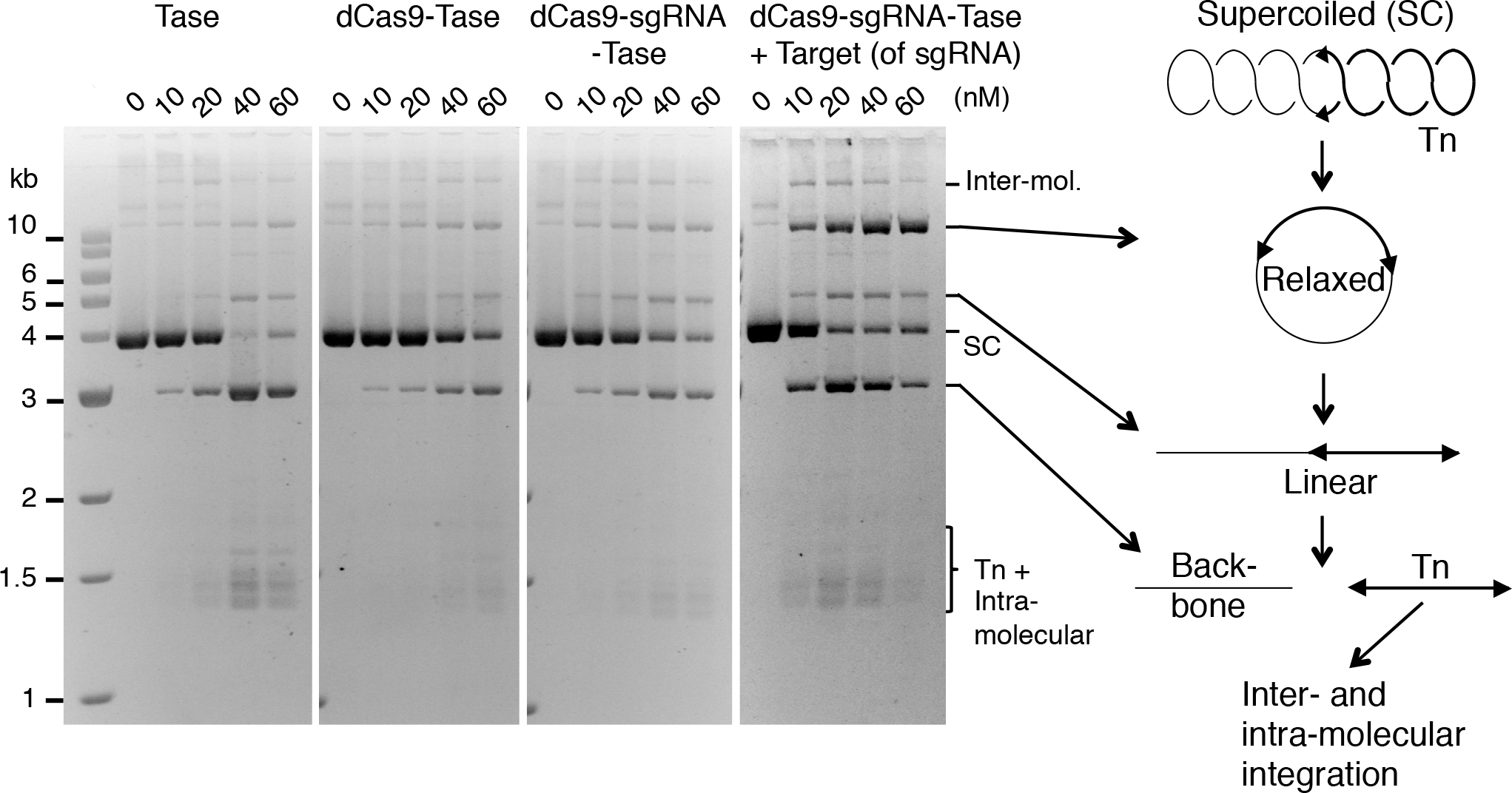
*In vitro* transposition assays with the dCas9-transposase. Transposition reactions were 50 *µ*l with 6.5 nM of supercoiled plasmid substrate (pRC650). The reactions were initiated by adding transposase and incubated for 4 h at 37ºC. Reactions were deproteinated with proteinase K and SDS and 20 *µ*l was electrophoresed on a TBE buffered 1.1% agarose gel at 60 V overnight. Photographs of ethidium bromide stained gels are shown. When present, the sgRNA was preassembled in a 1:1.5 molar excess to the protein prior to the reaction. Where indicated, reactions contained 10 nM of a 70 bp oligoduplex complementary to the sgRNA.

### Targeted transposition reactions

To test whether the dCas9-transposase protein could target transposon insertions to a specific site in vitro we used a modification of the mini-transposon hop assay (Munoz-Lopez et al., 2008). In this assay, transposase catalyses the movement of a transposon encoding a kanamycin resistance gene onto a target plasmid, which in this case encodes the lacZα fragment and an ampicillin resistance gene (Figure 5A). The target was derived from a dimer of pBluescript in which one of the two lacZα fragments had been removed. Since the plasmid has two ampicillin genes and two origins, it has no essential regions and can tolerate insertions anywhere. This allows unbiased recovery of integration events. We refer to the modified pBluescript dimer as the small target. We also created a larger target by adding 4.5 kb of DNA from the E. coli arginine biosynthesis operon. In addition to the transposon donor and target plasmids, transposition reactions also contained a decoy plasmid which provides a background level of non-specific DNA. The reactions were assembled and then initiated by the addition of the transposon donor plasmid.

**Figure 5.**
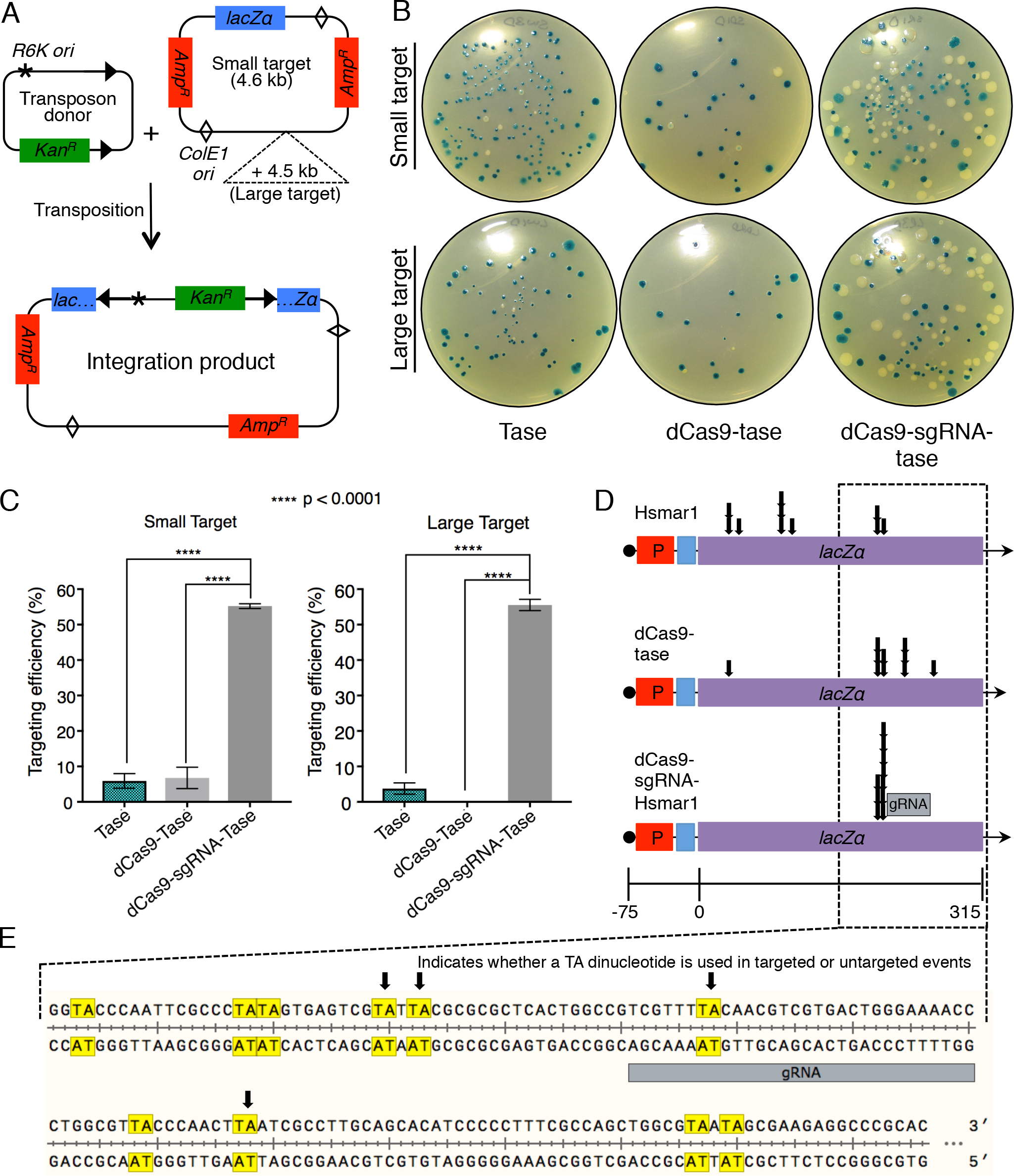
Transposon targeting reactions. **A**, Illustration of the transposon donor and target plasmids. The donor plasmid (pRC704) encodes and Hsmar1 transposon with an R6K origin of recognition and a kanamycin resistance gene. The target plasmid (pRC2312) is essentially a dimer of pBluescript with one copy of the lacZα gene removed. Since the plasmid has two origins and to ampicillin markers it can tolerate transposon insertions anywhere. The large target has an extra 4.5 kb of DNA non-specific DNA. Reactions also contained a background of non-specific DNA. **B**, Reactions were performed with the indicated transposase proteins, transformed into E. coli and plated on LB plus ampicillin, kanamycin and X-gal. Transposon integrations into the lacZα gene yield white colonies. Integrations elsewhere yield blue colonies. **C**, Targeting efficiency. Error-bars are standard error of the mean where n = 6 biological replicates. Ordinary one-way ANOVA analysis: Small target; transposase vs. dCas9-transposase, P = 0.95; transposase vs. dCas9-sgRNA-transposase, P = <0.0001; dCas9-transposase vs. dCas9-sgRNA-transposase, P = <0.0001. Large target: transposase vs. dCas9-transposase, P = 0.14; transposase vs. dCas9-sgRNA-transposase, P = <0.0001; dCas9-transposase vs. dCas9-sgRNA-transposase, P = <0.0001. **D**, Ten white colonies were picked from the three sets of plates and the location of the transposon insertion determined by Sanger sequencing. **E**, An expanded view of the indicated area of the lacZα gene.

Transposition reactions were performed with the wild type transposase and the dCas9-transposase fusion-protein either without or with sgRNA-7, which is complementary to the lacZα region of the target plasmids. The integration products were transformed into E. coli NEB5α and plated on LB agar plates supplemented with ampicillin, kanamycin and X-gal (Figure 5B, C). Double transformants containing the donor and target plasmids are not recovered because the R6K origin of replication requires the Pir protein. Transposition reactions with the wild type transposase or the dCas9-transposase fusion yielded very few white colonies (Figure 5B). This reflects the fact that the lacZα fragment represents only about 1/650^th^ of the DNA sequences available for integration. In contrast, more than half of the colonies are white when the dCas9-transposase protein is provided with the sgRNA. The targeting efficiency is plotted in Figure 5C where it is defined as the percentage of all colonies that were white.

In the case of the blue colonies, the transformants must have contained plasmids with transposon insertions outside of the *lacZα* region. We therefore focused on the white colonies and used DNA sequencing to determine the transposon integration sites (Figure 5D). In pBluescript the *lacZα* open reading frame (ORF) is engineered to include a polylinker and it is therefore longer than the corresponding region from the wild type lacZ gene. The ORF also codes for additional amino acids at the 3’-end that are not necessary for alpha complementation (Nishiyama et al., 2015). All in all, the target region in pBluescript in which integration events will yield a white phenotype is 315 bp long (Figure 5D). Within this region there are 18 TA dinucleotides, which are required for the integration of mariner transposons. Only eight of these sites are represented amongst the 20 un-targeted integrations mapped for the wild type transposase and the dCas9-transposase fusion without the sgRNA (Figure 5D). This is not unusual as mild integration bias is well documented for the mariner transposons, where preferred sites are known as hot spots (Claeys Bouuaert and Chalmers, 2013; Conte et al., 2019; Crenes et al., 2009).

We next examined the lacZα integration sites targeted by the dCas9-sgRNA-transposase. We found that all had occurred at two TA dinucleotides located 18 to 22 bp to one side of the sgRNA binding site (Figure 5D, E). On the other side of the binding site there are TA dinucleotides between 8 and 18 bp distant that were not used. The location of the integration sites immediately to one side of the binding site probably reflects constraints imposed by the relative orientations of the dCas9 and the transposase domains and the length of the linker region. It will be interesting to test the site-preference of the transposase-dCas9 fusion, in which the positions of the domains are swapped with respect to the N- and C-termini of the protein.

## DISCUSSION

The long dwell time of dCas9 at its target site makes it a powerful tool for gene activation and repression (Gilbert et al., 2014). It is also an attractive candidate for transposon targeting. However, one study reported that although piggyBac integrations at a specific target site were enriched by zinc finger and TALE protein-fusions, dCas9 appeared to protect the locus (Luo et al., 2017). In the present work we have demonstrated that a mariner transposon can be effectively targeted by dCas9 and that protection from integration is probably therefore peculiar to piggyBac. We found that the targeted mariner insertions were at two adjacent TA dinucleotides about 20 bp 5’ to the sgRNA binding site (Figure 4E). Other TA dinucleotides located between 31 and 34 bp 5’, and between 8 and 18 bp 3’, of the sgRNA binding sites were not used. This indicates that the integration site is tightly constrained.

The properties of the inter-domainal linker and the intrinsic target-site preference of the transposase may both contribute to the constrained selection of integration sites. Most, if not all, DNA transposon have preferred target hot spots. For example, there is one extremely hot spot for Mos1 integration that attracts almost all events in the Tn9 chloramphenicol acetyl transferase gene (Crenes et al., 2009). Hsmar1 is perhaps less biased but some sites are certainly less preferred than others (Claeys Bouuaert and Chalmers, 2013). To link the dCas9 and Hsmar1 transposase domains we used a sequence of 187 amino acids encoding the E. coli thioredoxin protein. Although this will be less prone to proteolysis than a flexible linker of the same length, the compaction due to elements of secondary and tertiary structure will prevent the transposase domain reaching distant sites. The linker may also constrain the orientation of the transposase domain with respect to dCas9 and its binding site. However, since the two targeted TA dinucleotides are 4 bp apart they will be on almost opposite faces of the DNA helix. This indicates that the angular distribution is not tightly constrained. Nevertheless, it will be interesting to engineer additional TA dinucleotides into the sequence to explore the constraint further. It will also be interesting to use flexible linkers of different lengths.

In targeting experiments with other transposase-fusion proteins the selection of integration sites was not tightly constrained as observed here. Fusing Mos1 or piggyBac transposases to Gal4 using a 22 amino acid linker enriched for integration sites about 900 bp away from the Gal4 upstream activating sequence (UAS). In the case of Mos1, 98% of the integrations were at a TA dinucleotide 954 bp from the binding site (Maragathavally et al., 2006). Such frequent use of a site, almost 10 DNA-persistence-lengths away from the Gal4-UAS, can not be explained by the constraints of the binding moiety. Instead, a combination of factors may be at work. For example, the short dwell time of DNA binding proteins, which range from seconds to minutes, will increase the local concentration of the transpososome, which may then select a nearby hot spot. This scenario may also explain the partial enrichment of AAV Rep-Sleeping Beauty integration at a site 700 bp from the Rep recognition sequence (Ammar et al., 2012). In another report, fusing the piggyBac transposase to zinc finger and TALE proteins with linkers as short as 15 amino acids enriched for integrations between 24 and 5000 bp from the binding site in the hypoxanthine phosphoribosyl transferase (HPRT) gene (Luo et al., 2017; Owens et al., 2013). Within this region, integrations were biased towards the HPRT transcriptional start site. This may be owing to the chromatin configuration or topological changes in the DNA. For example, mariner transposition is sensitive to supercoiling in the transposon donor and the target (Claeys Bouuaert and Chalmers, 2013; Claeys Bouuaert et al., 2011).

In addition to the targeting experiments, we also compared the reaction intermediates with the transposase and the dCas9-transposase proteins (Figure 2, 3). In the EMSA, the nucleoprotein complexes were very similar (Figure 2). Likewise, in reactions with a transposon on a plasmid substrate, the kinetics of the reaction intermediates were similar (Figure 3). However, there was a slight delay between first and second strand cleavage at the transposon end when the target of the sgRNA was included in the reaction. Overall, it is clear that the Hsmar1 transposase behaves normally when fused to dCas9 and that dCas9 is therefore a promising system for targeted integration. The single biggest problem in targeted-transposition protocols is that successful targeting in a particular cell is often accompanied by random integrations. It should be possible to minimize this problem delivering the transpososome to the cells after in vitro assembly (Trubitsyna et al., 2017). However, to overcome random integration events, it will be necessary to control the timing of integration to allow an opportunity for the targeting moiety to find its binding site. This work is currently underway.

---

**Supplemental Figure S1.**
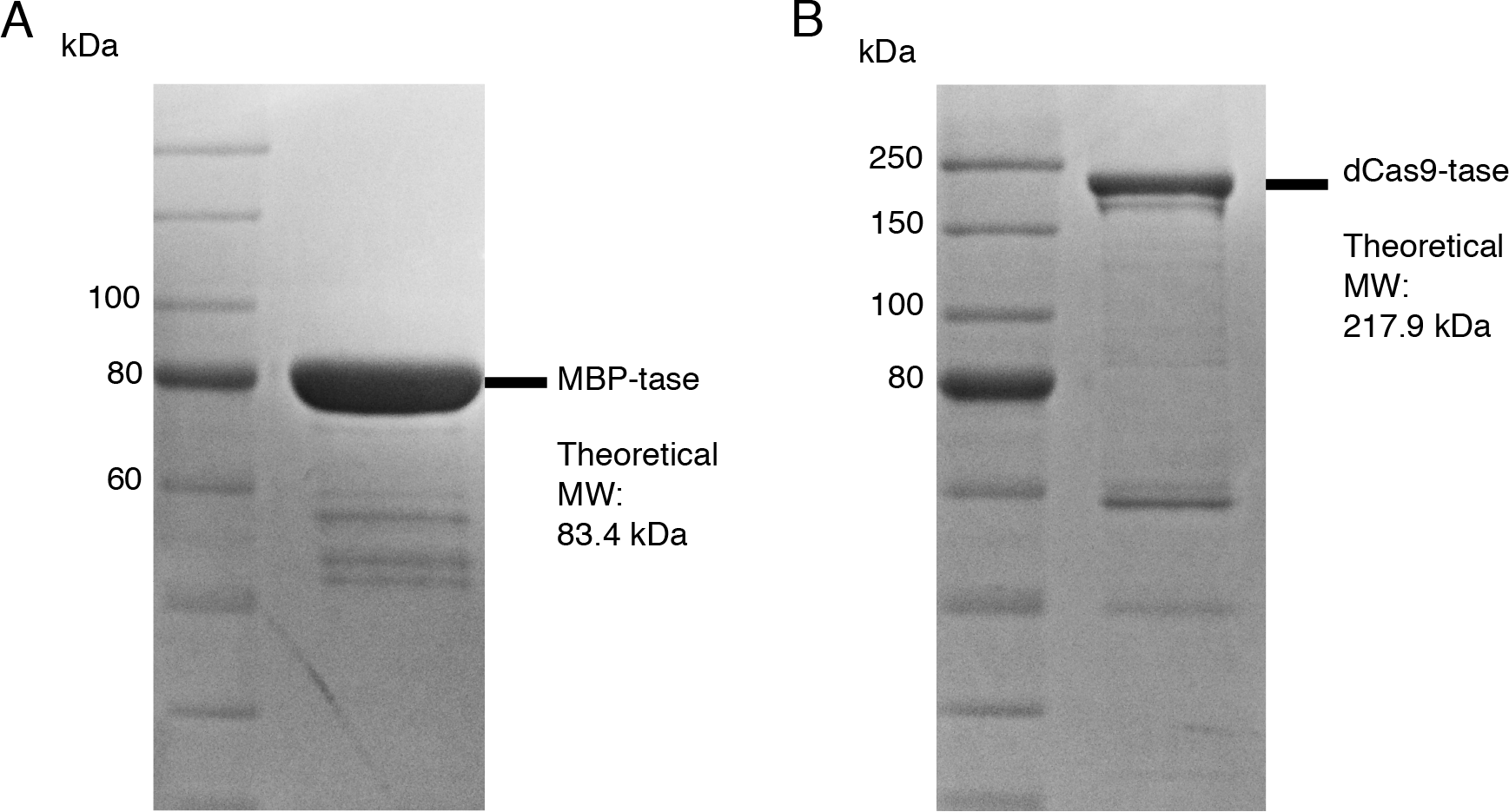
Purification of Hsmar1 transposase and dCas9 transposase.

**Supplemental Table 1.**
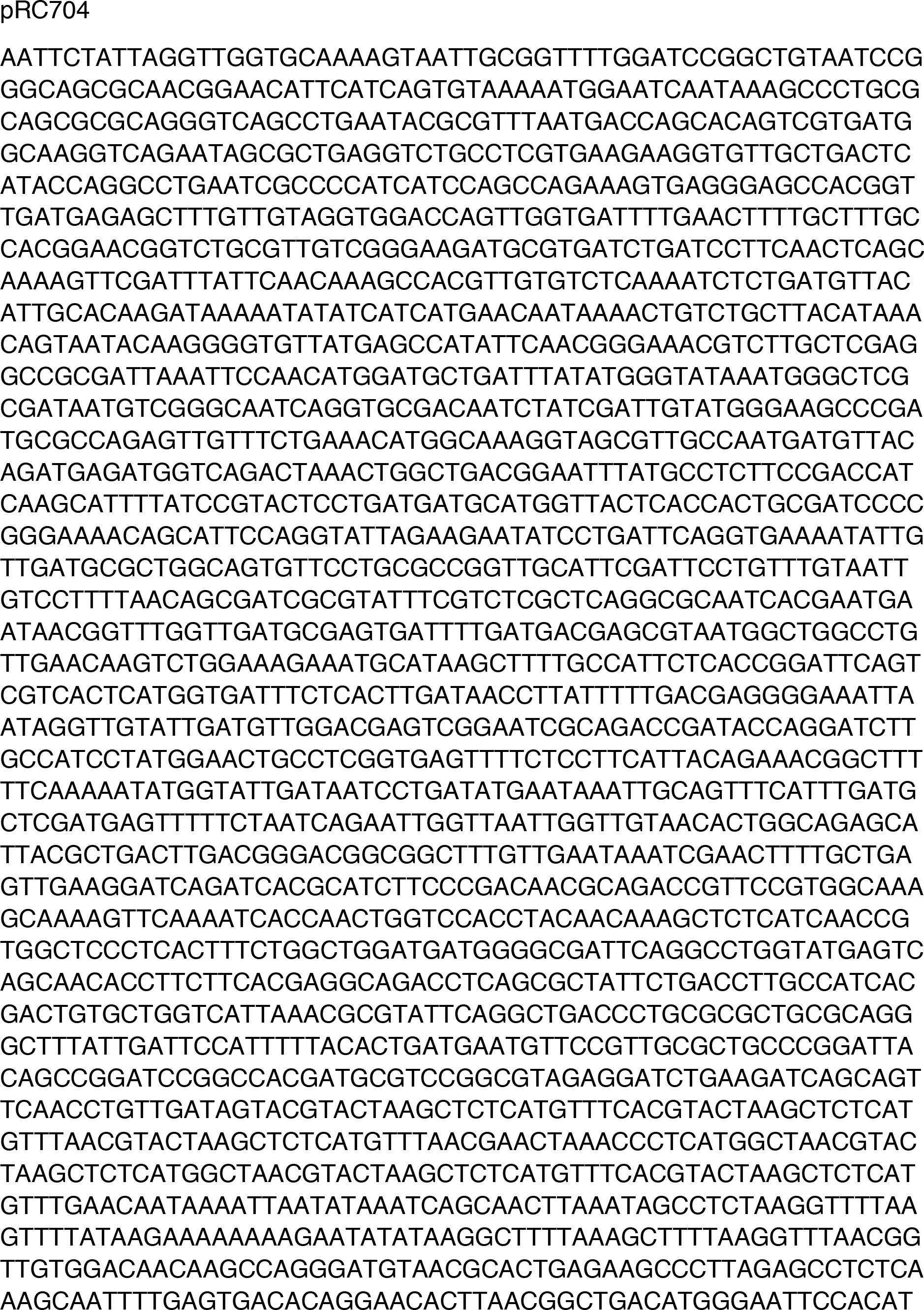
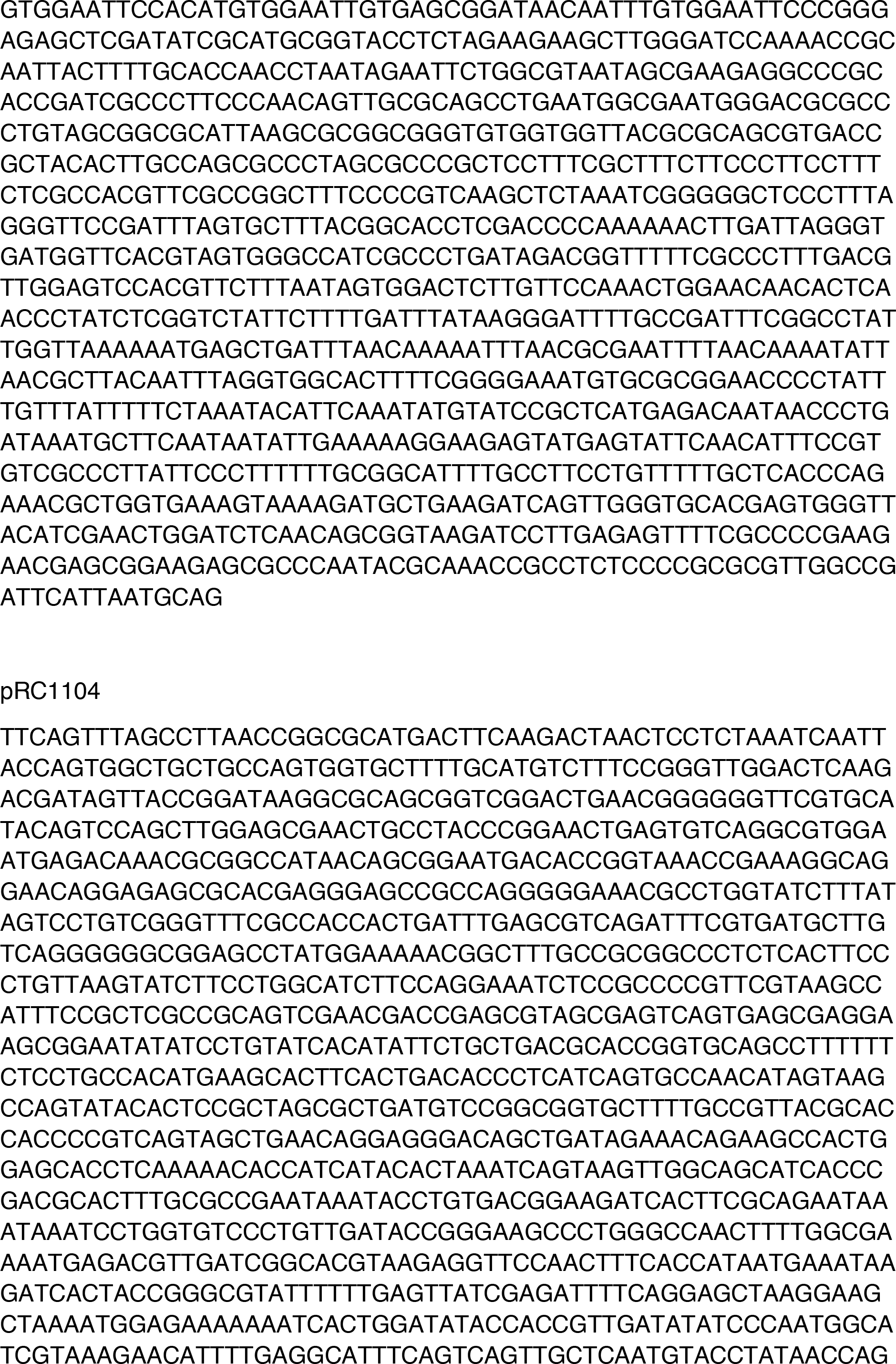
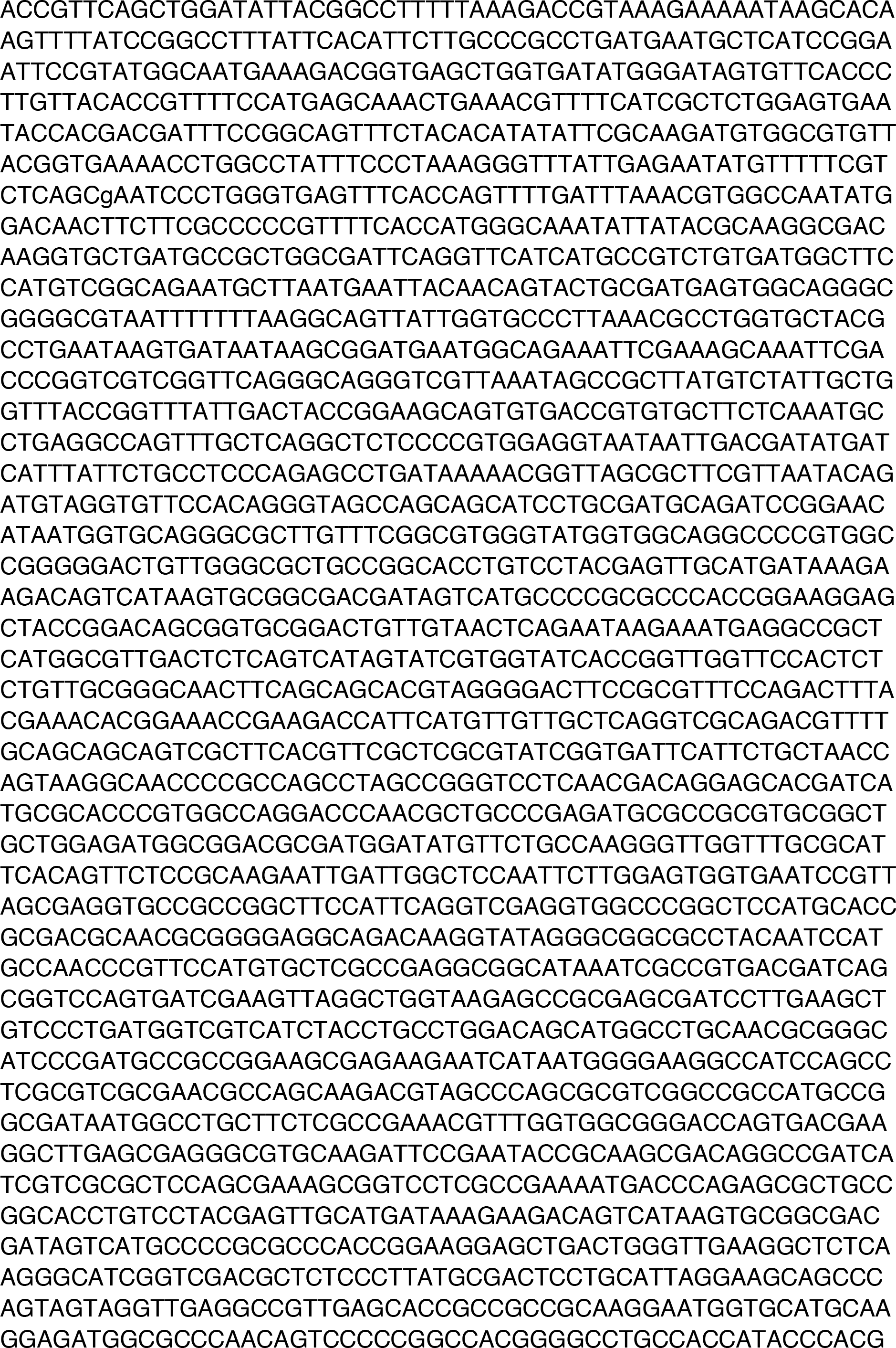
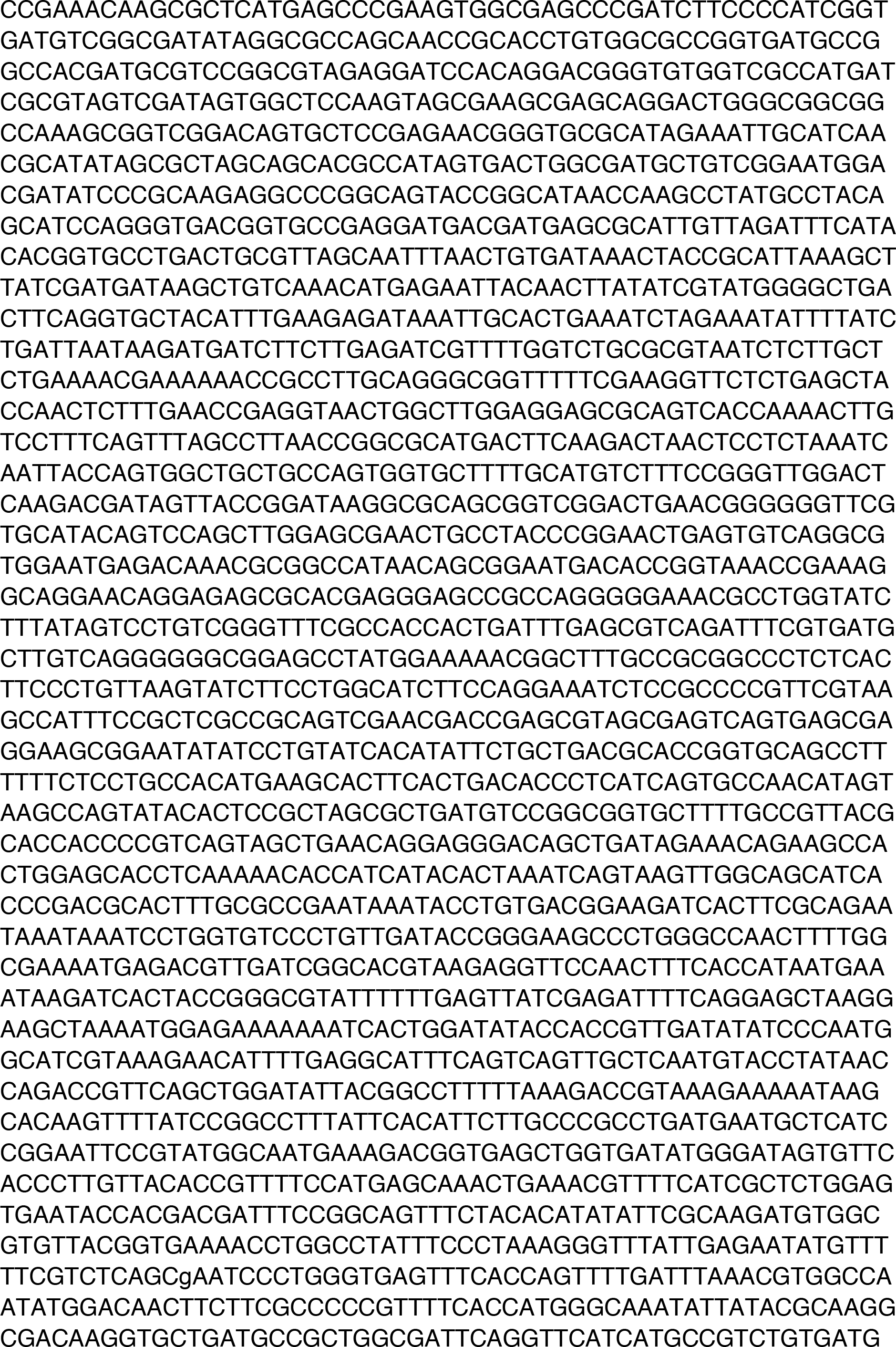
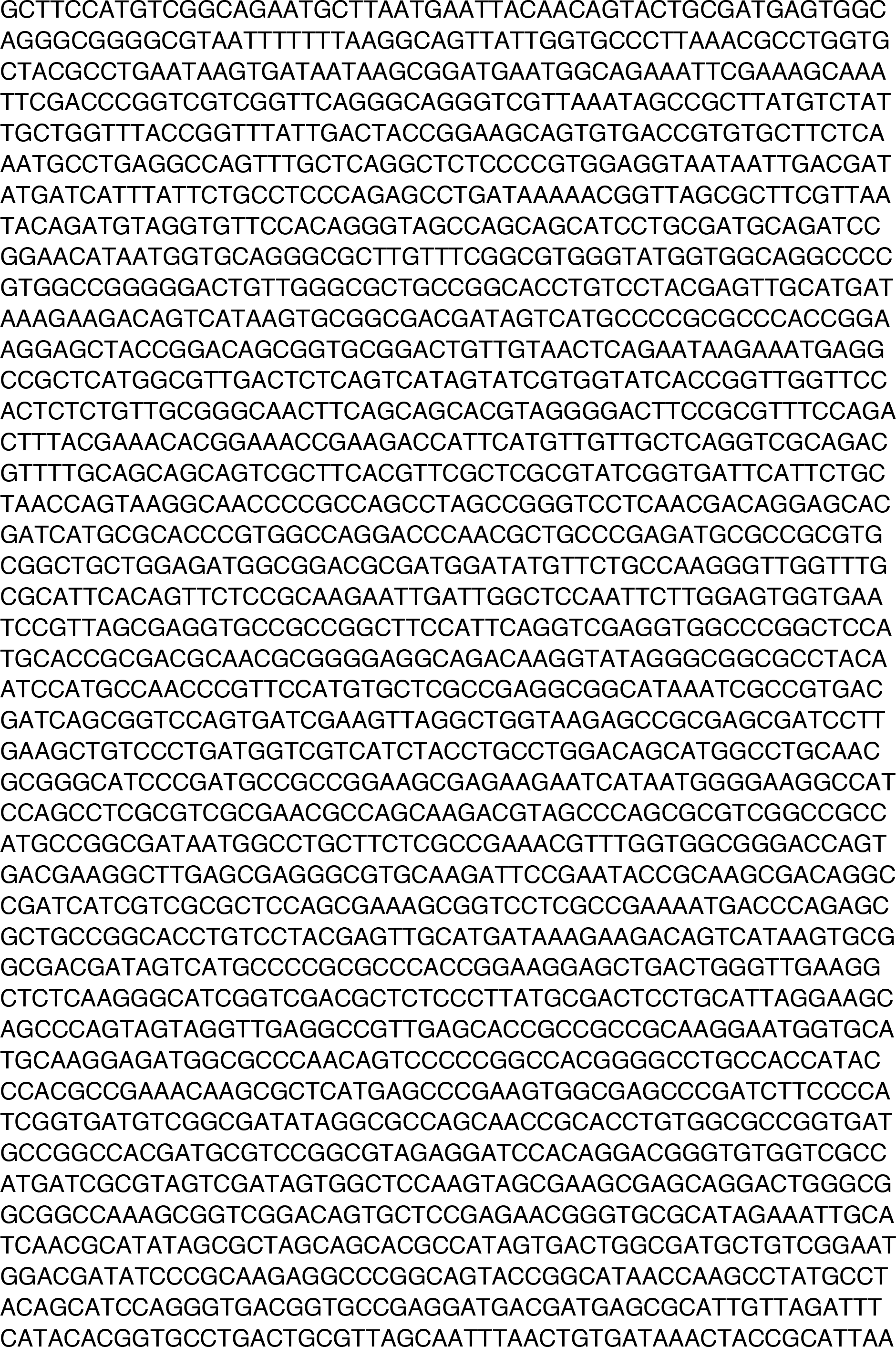
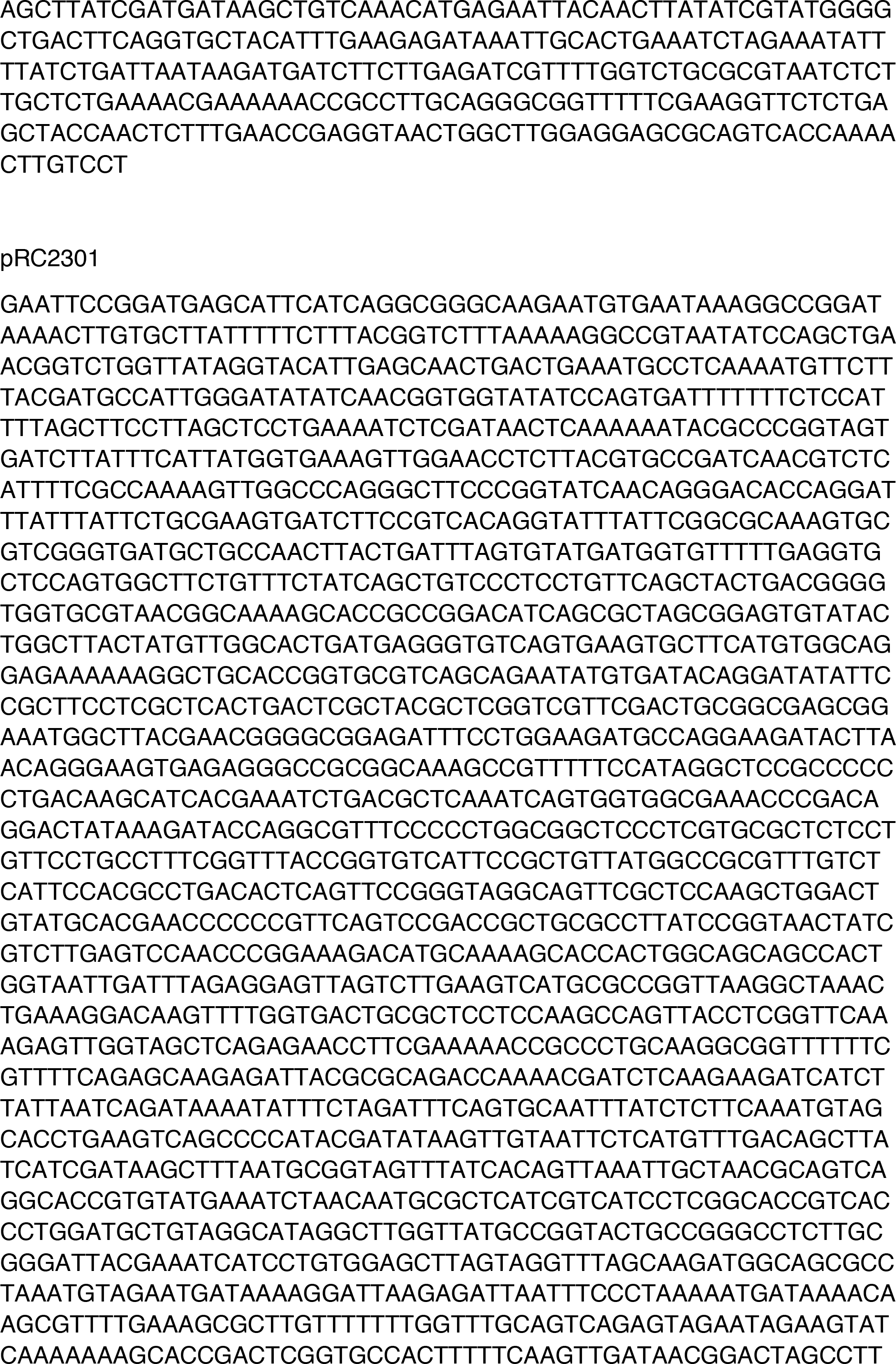
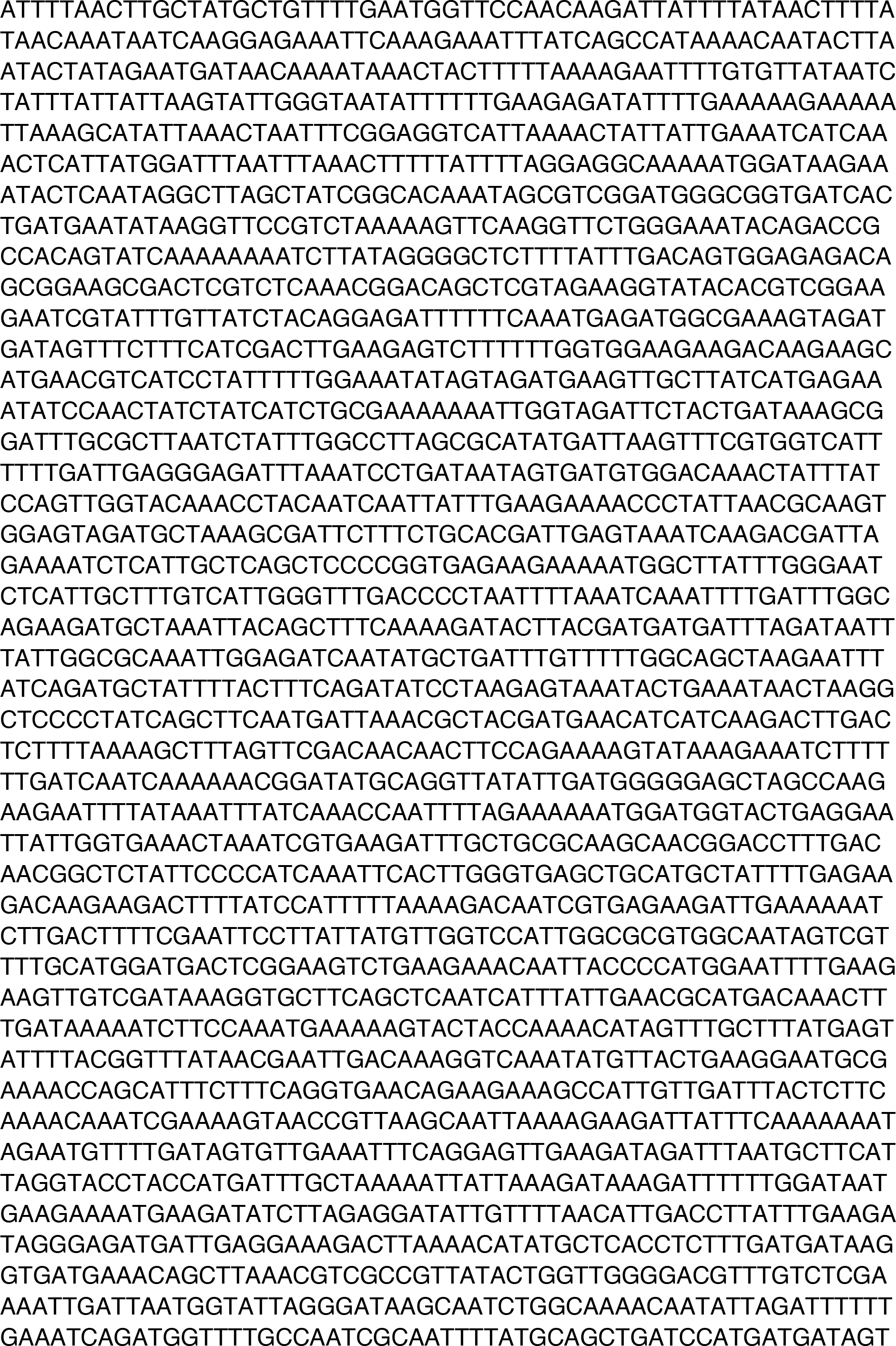
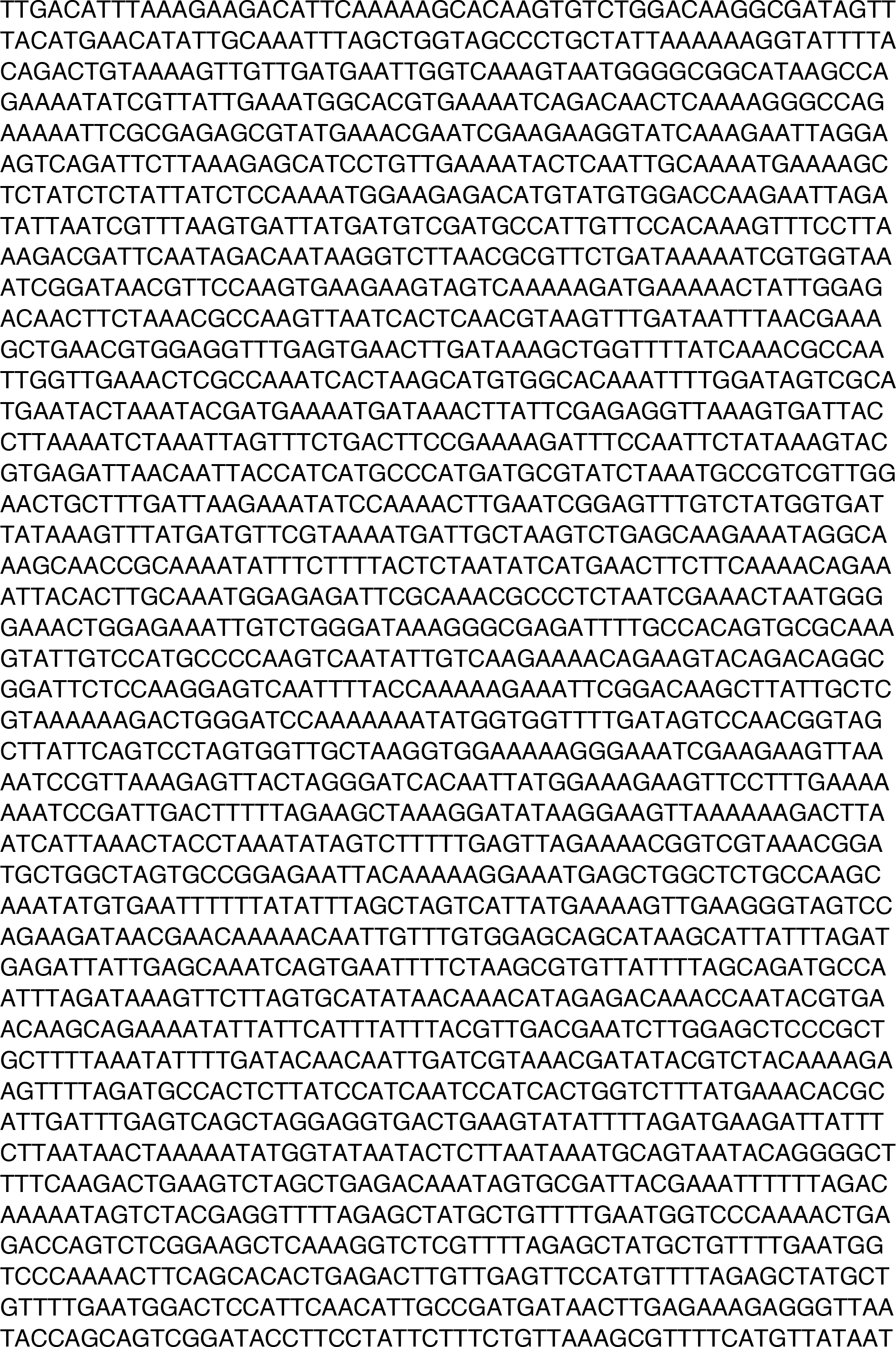
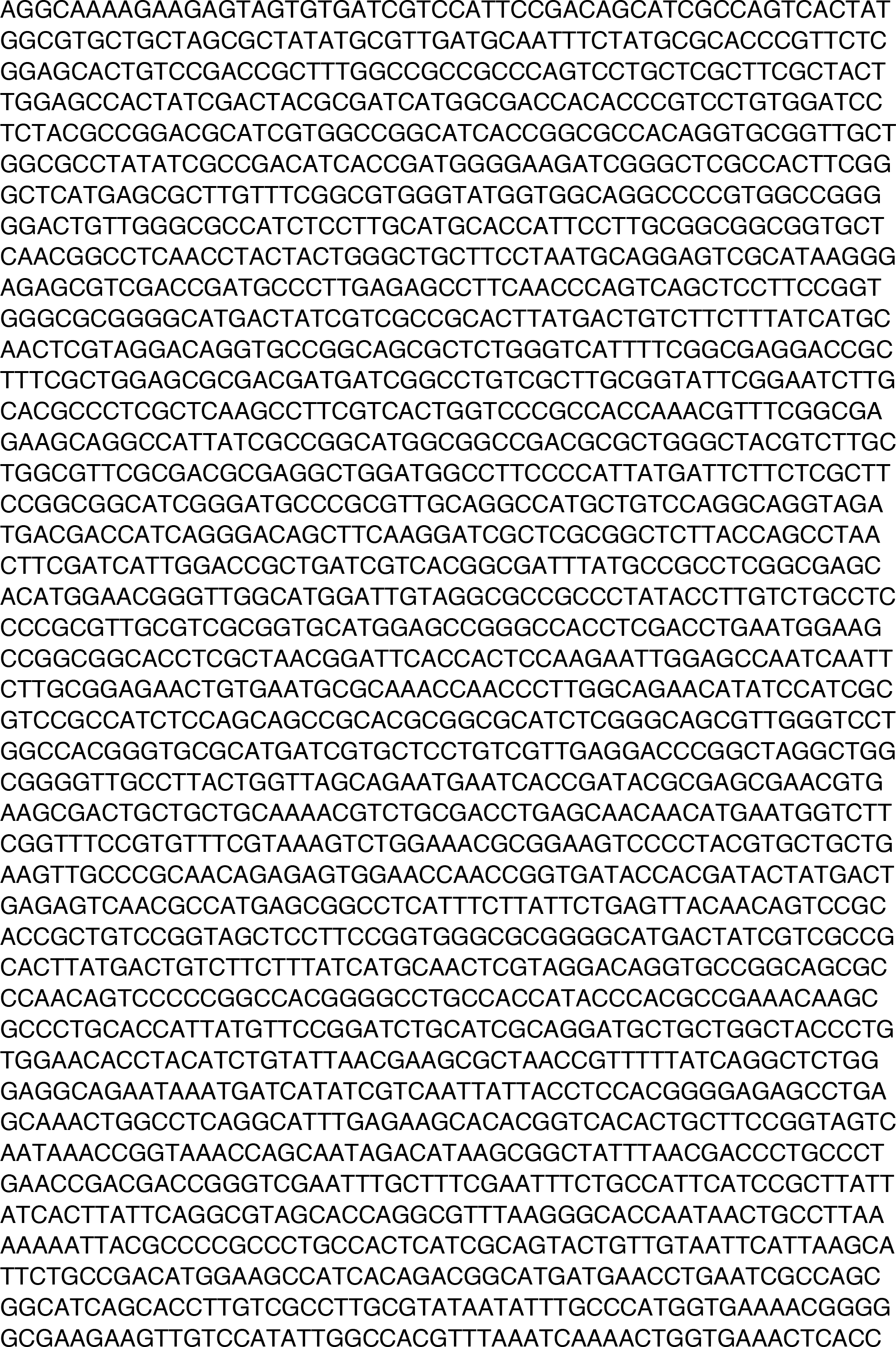
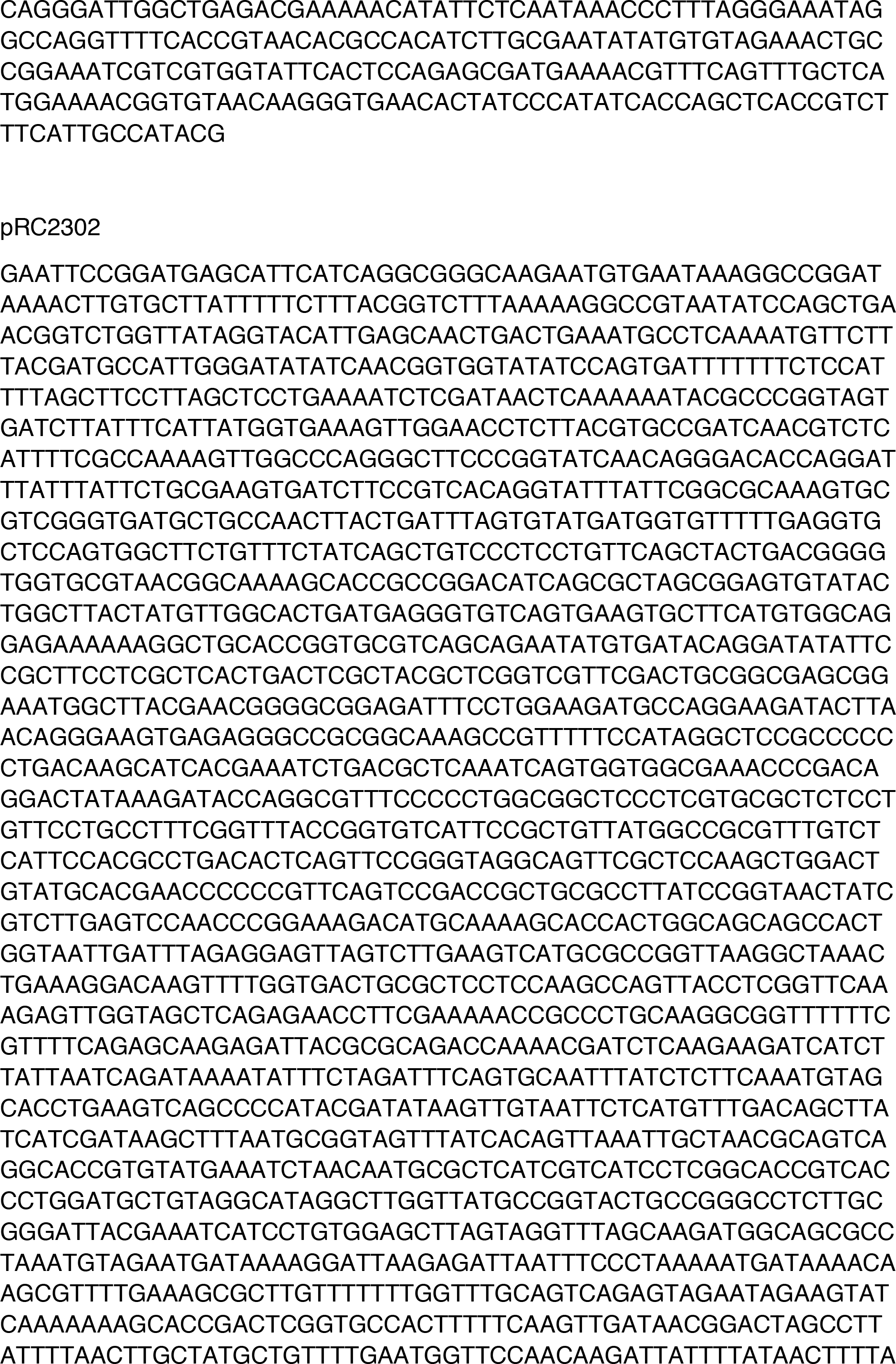
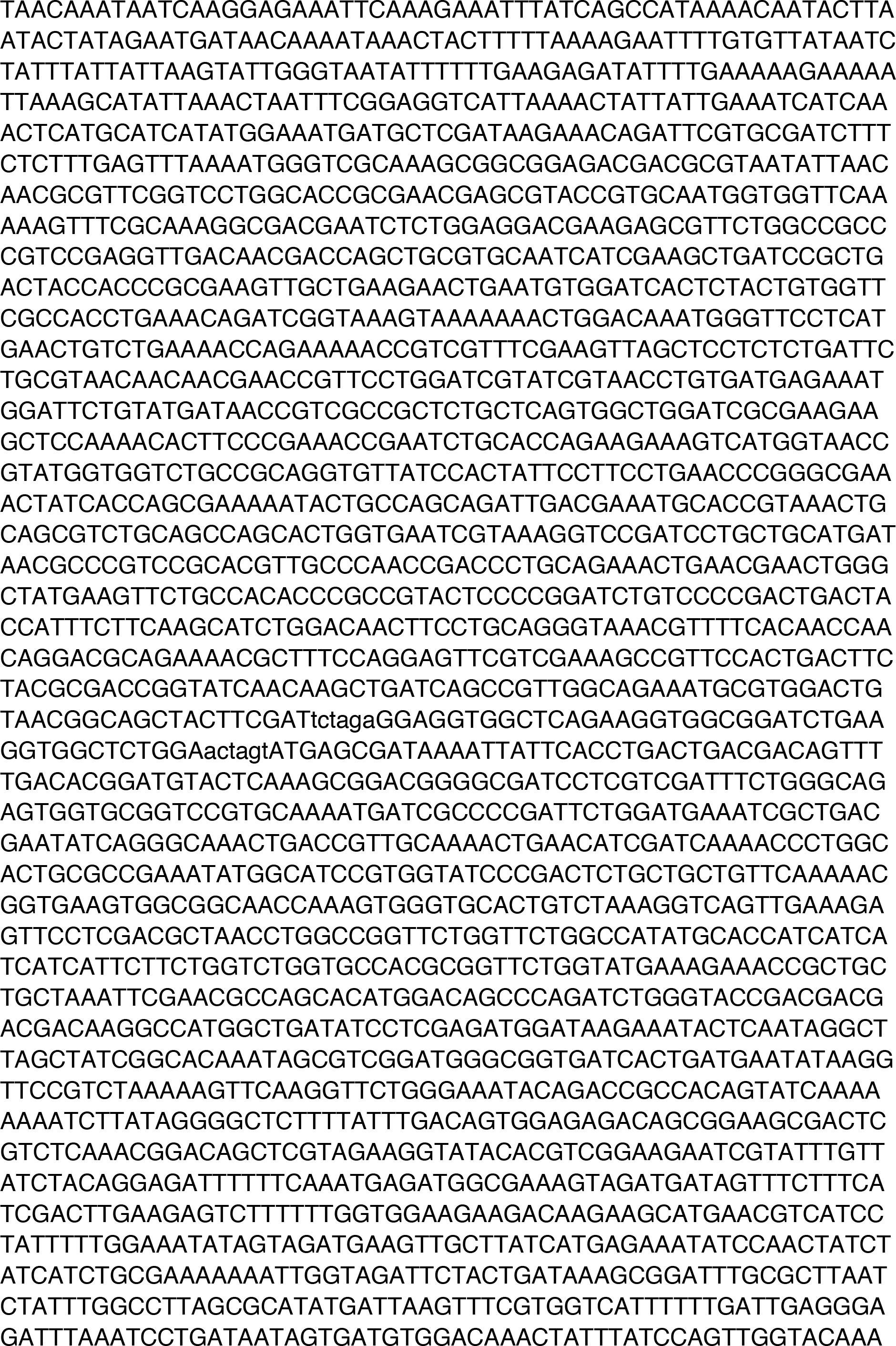
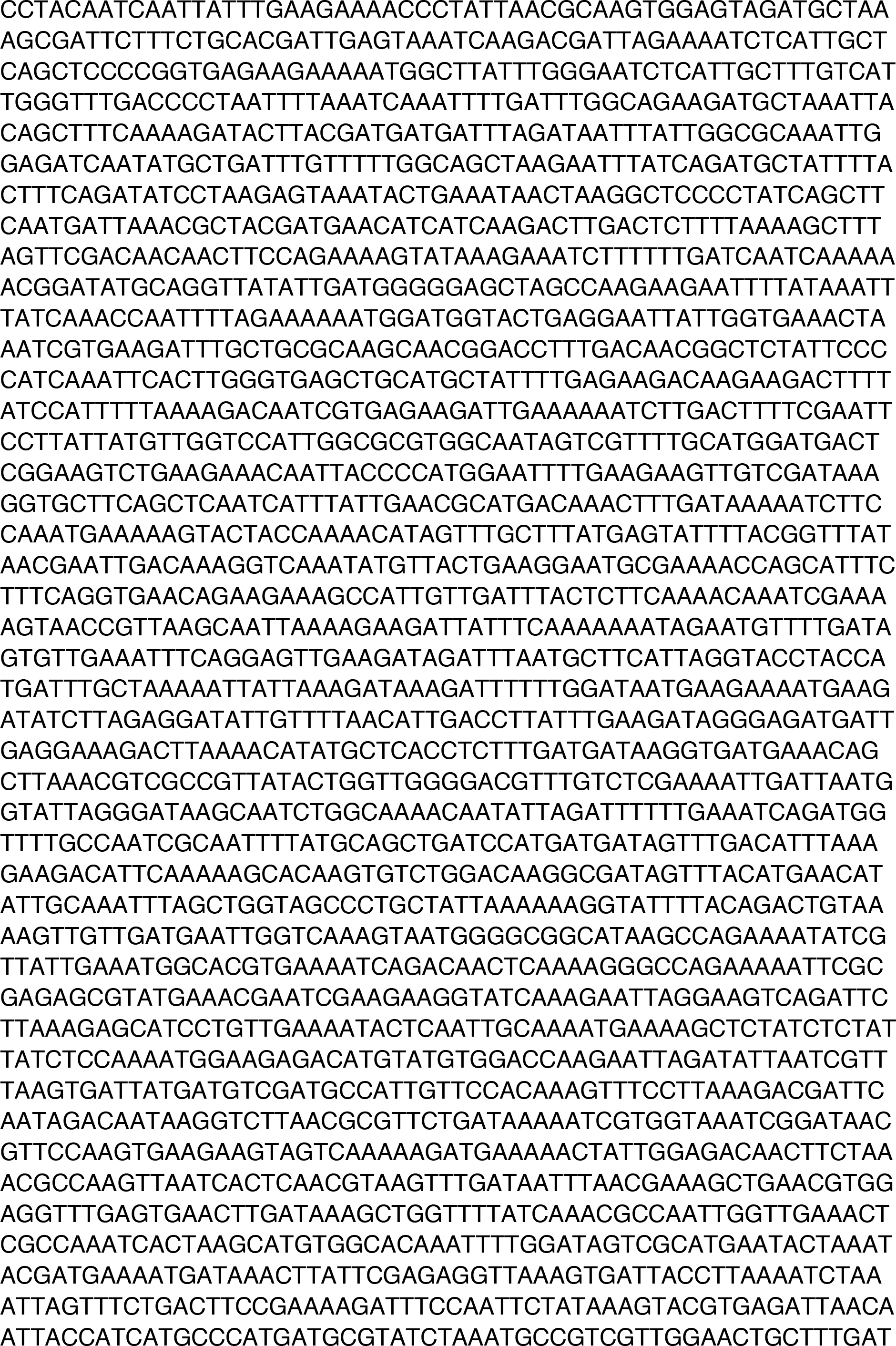
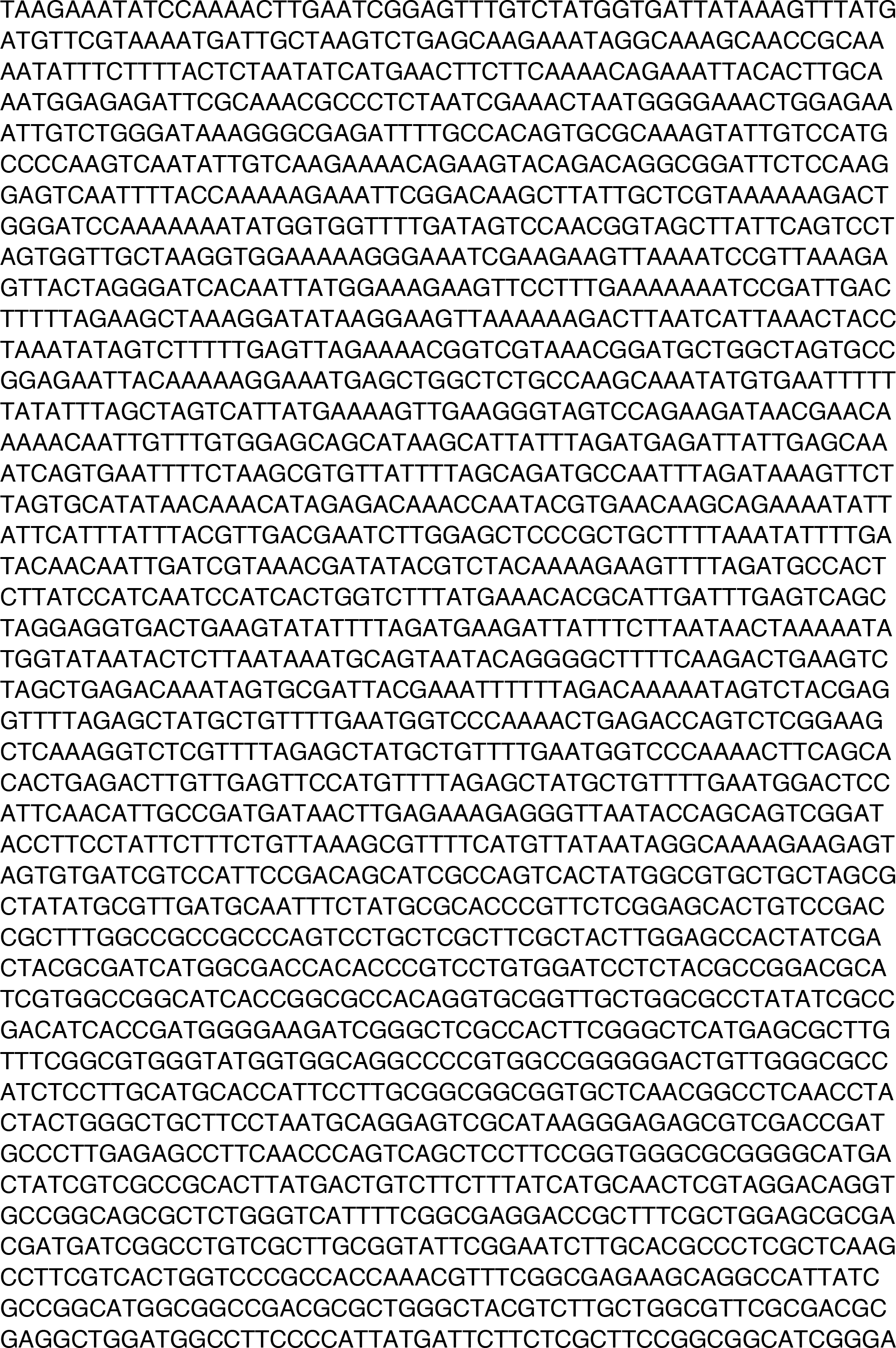
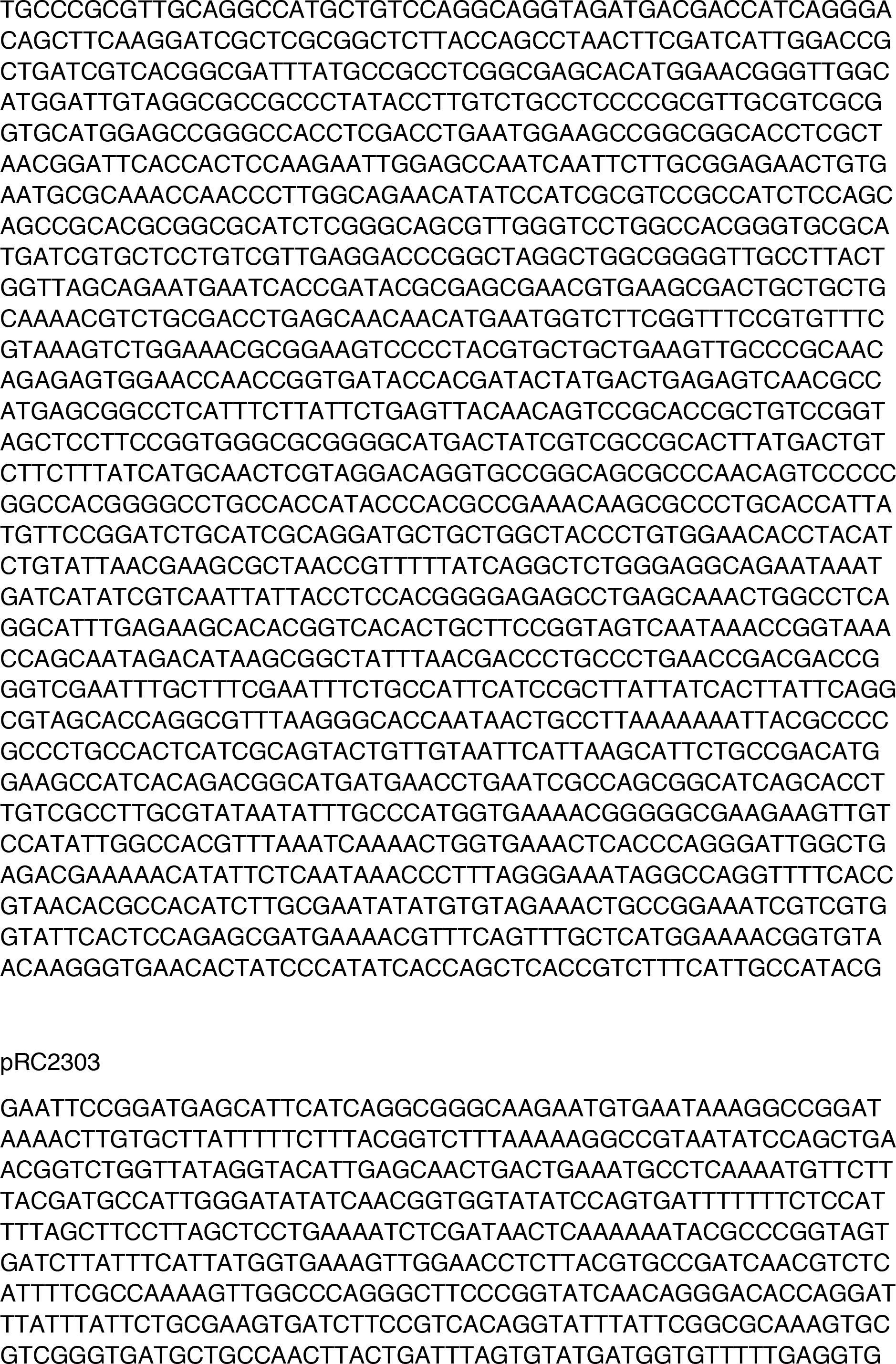
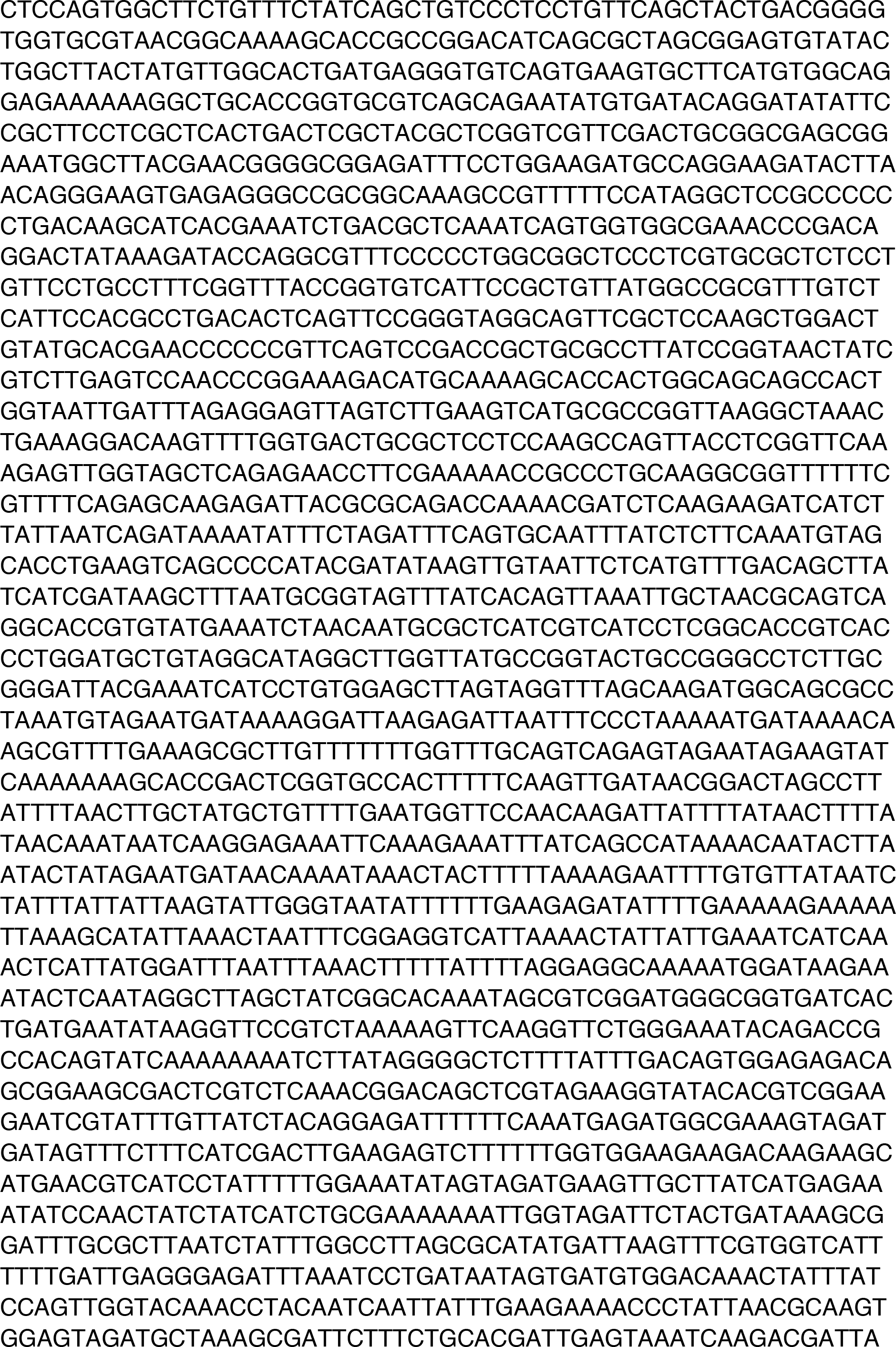
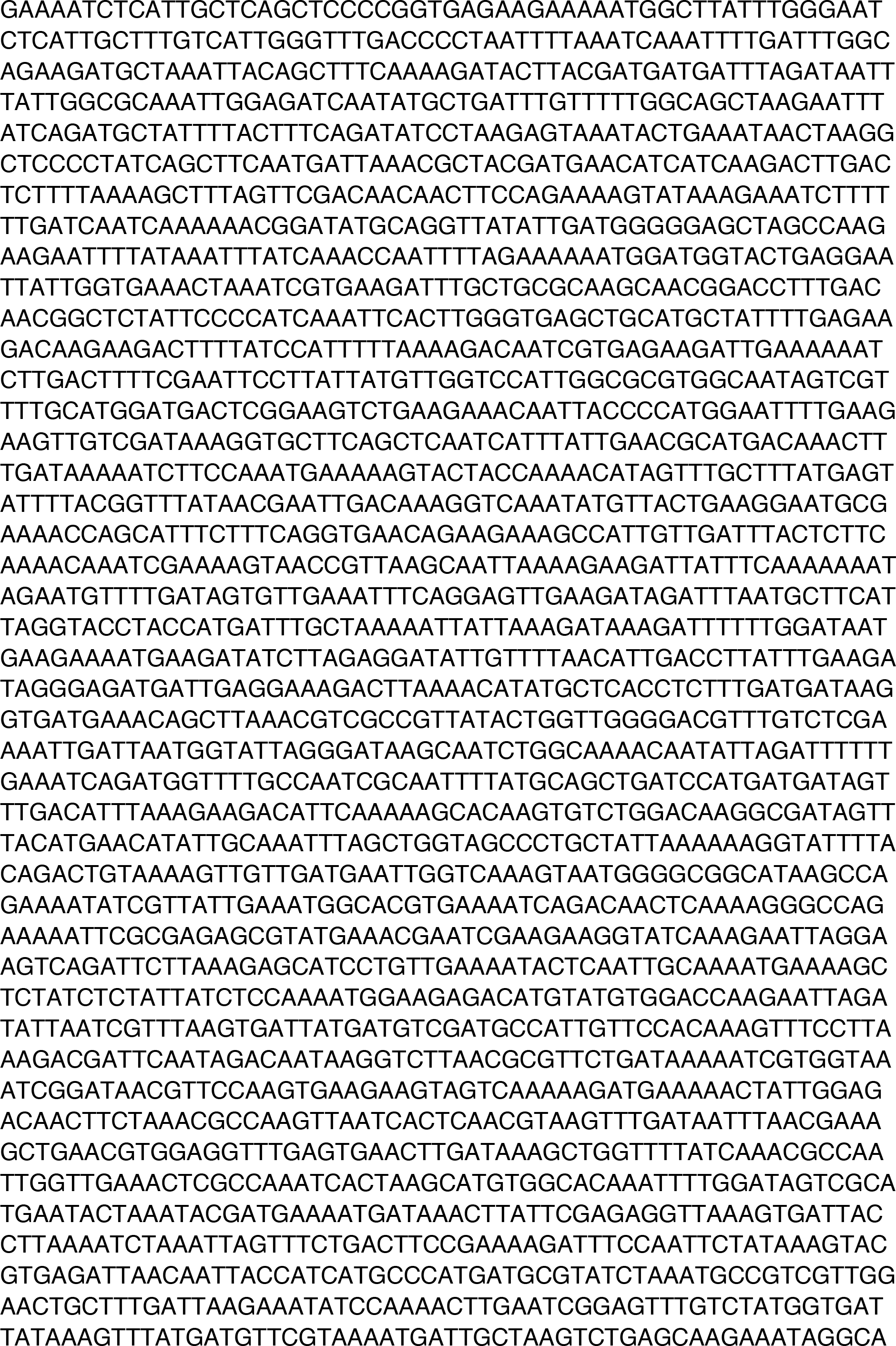
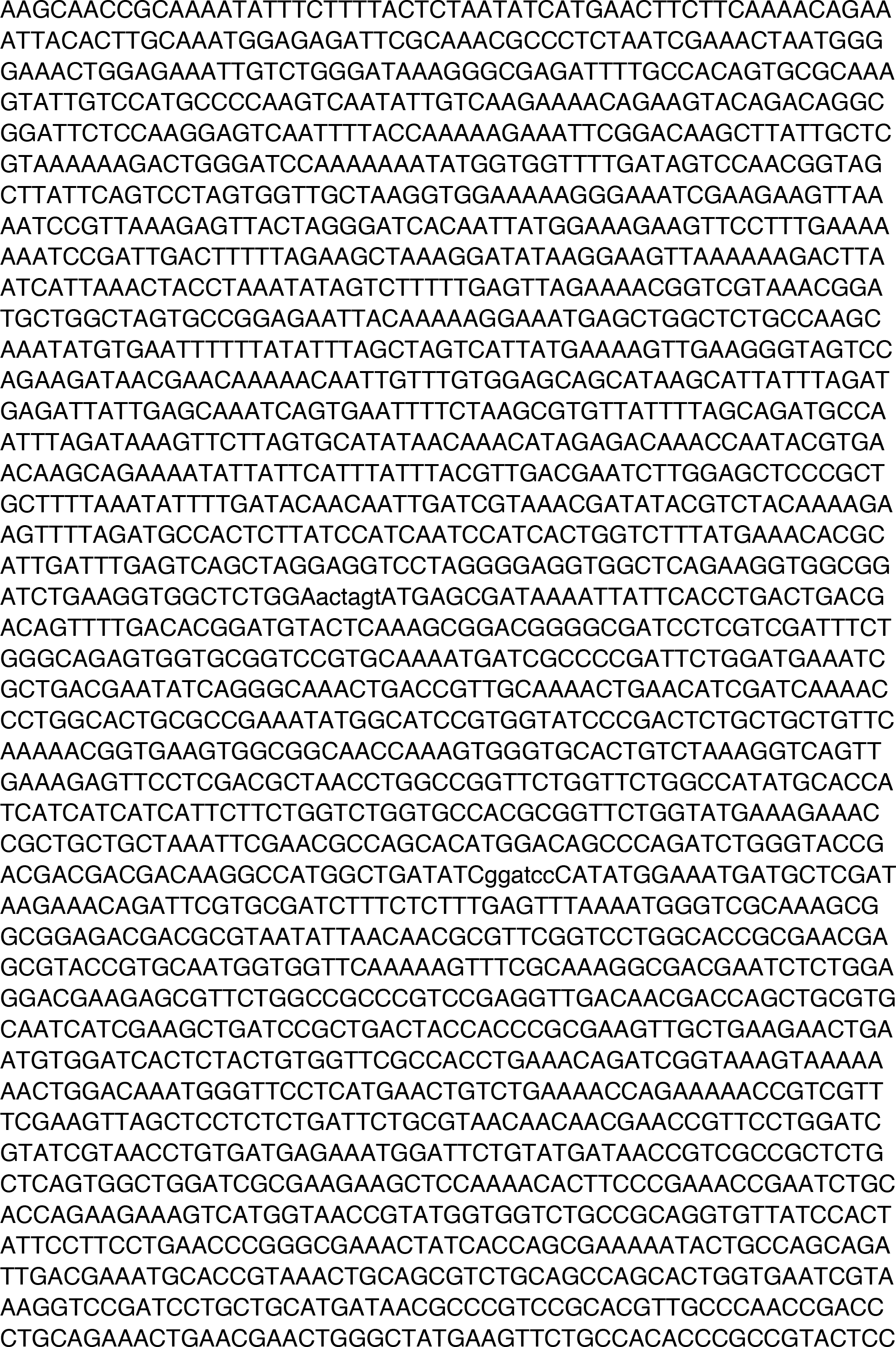
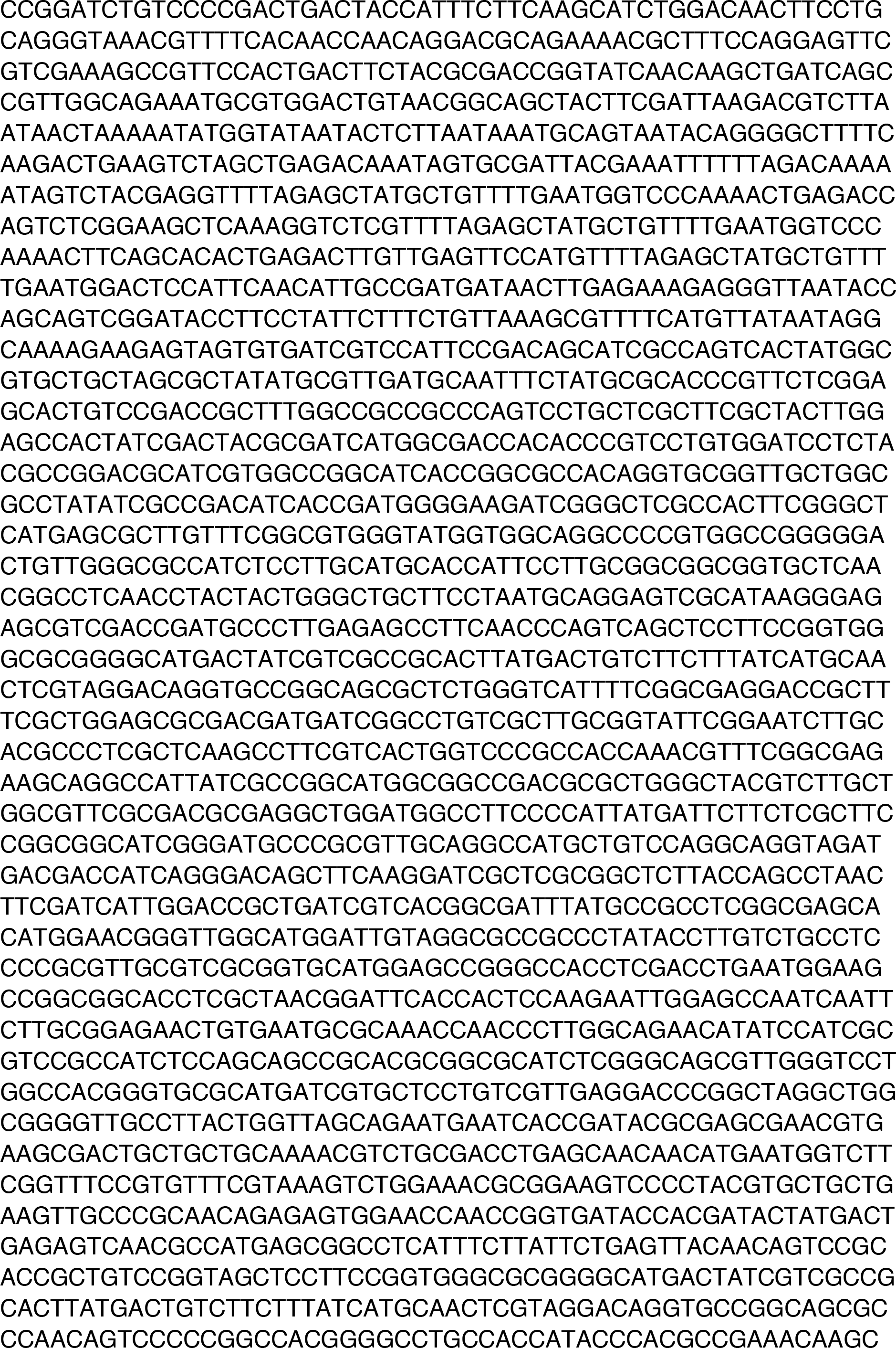
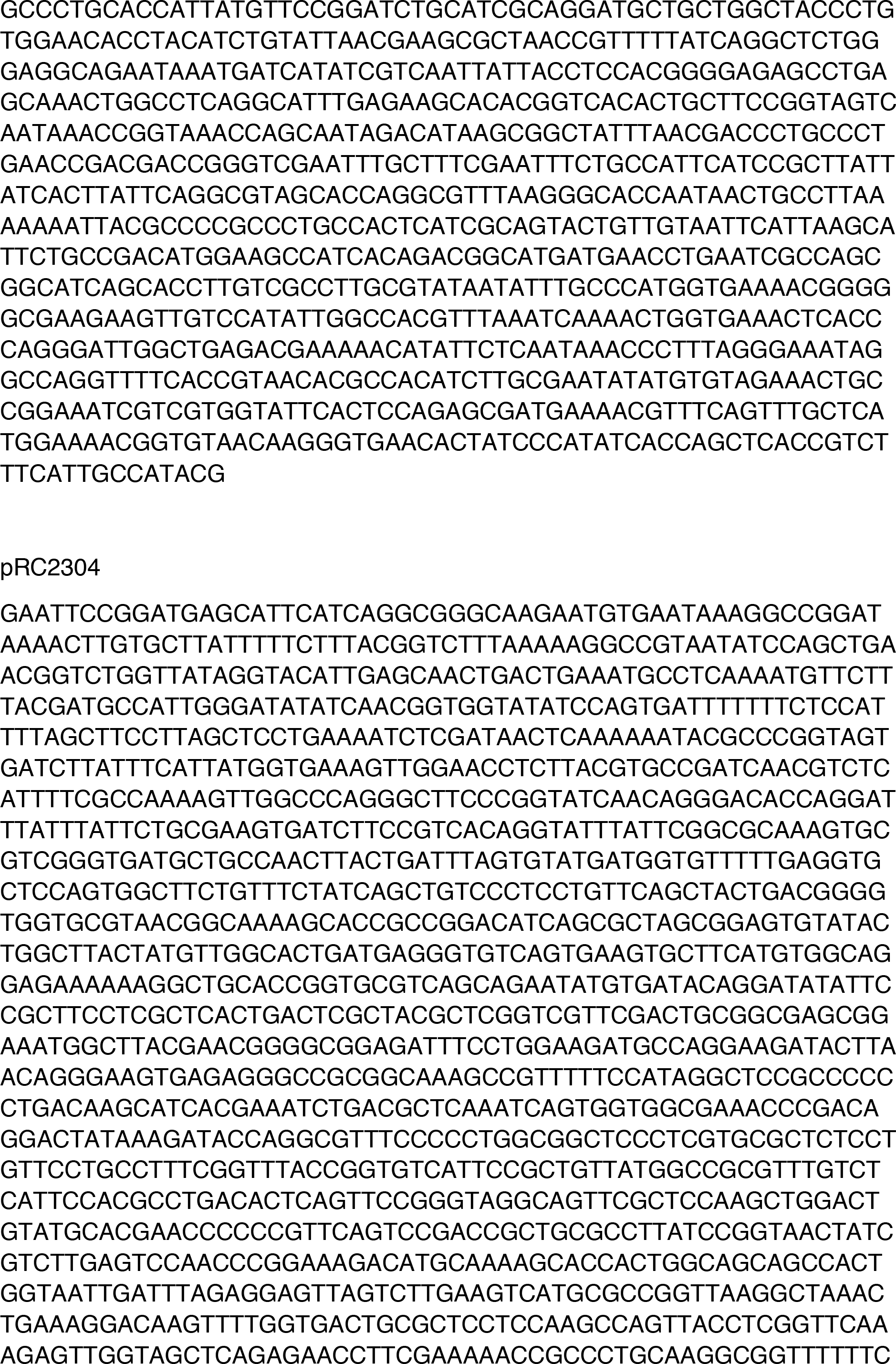
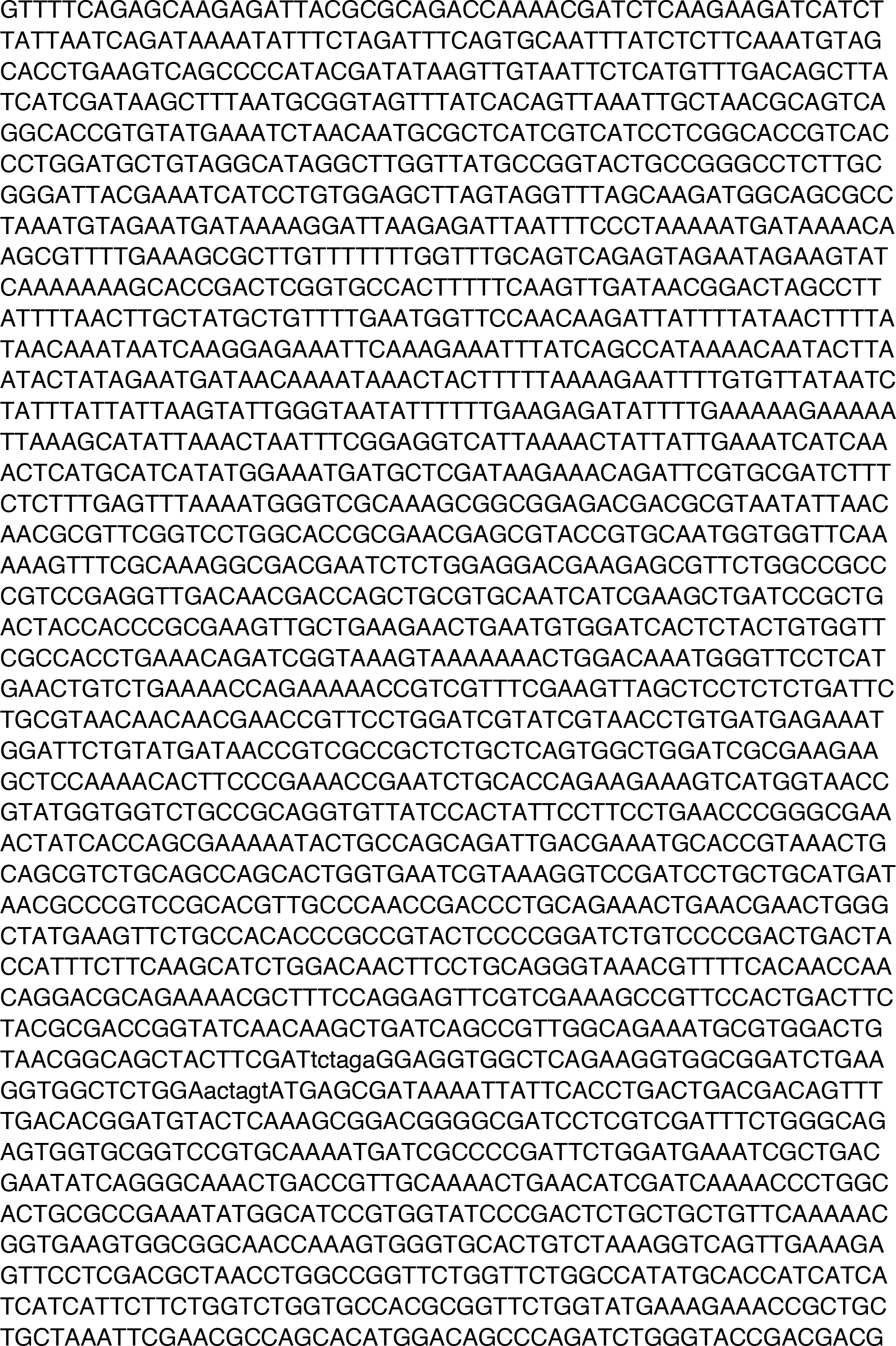
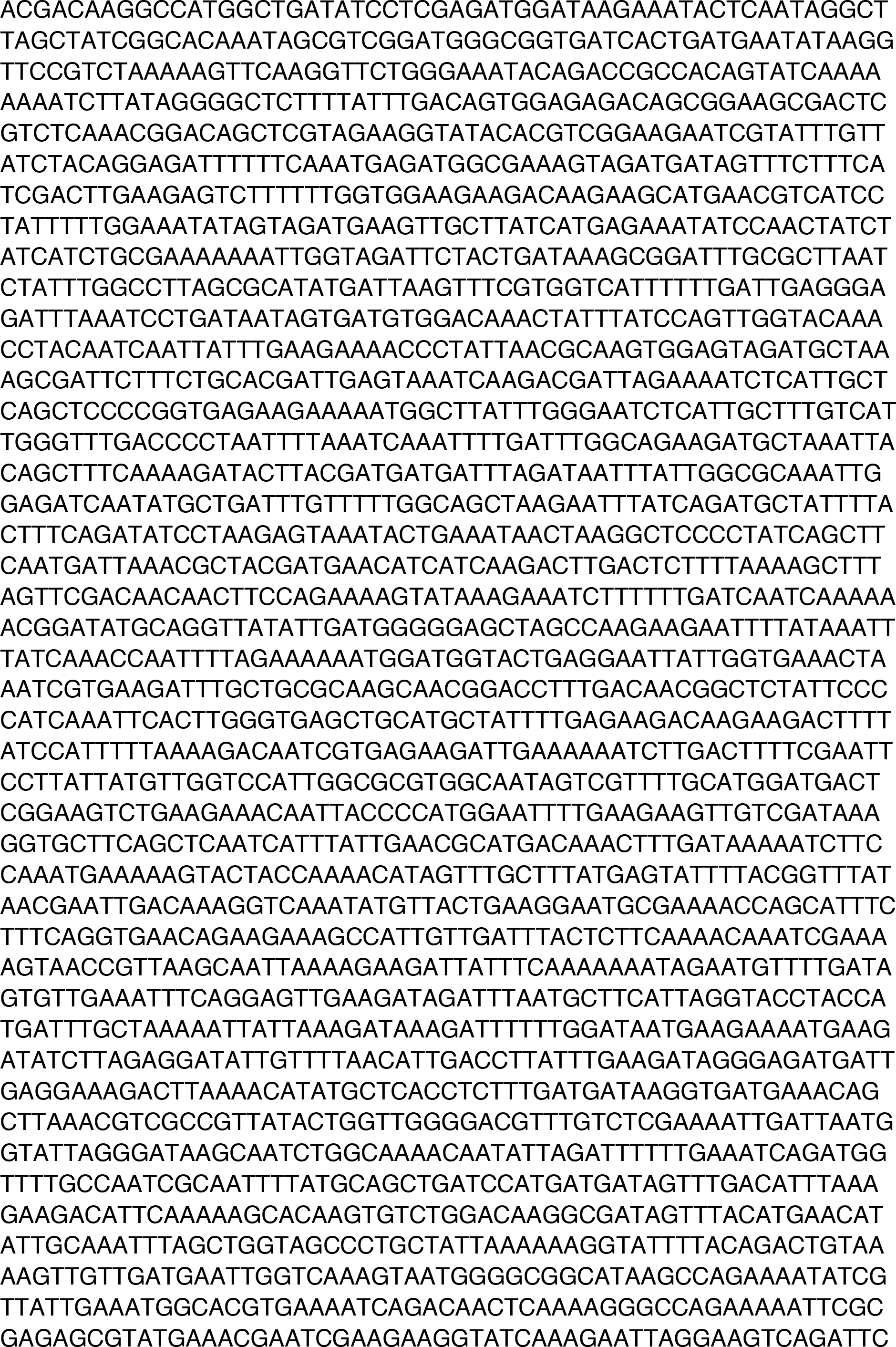
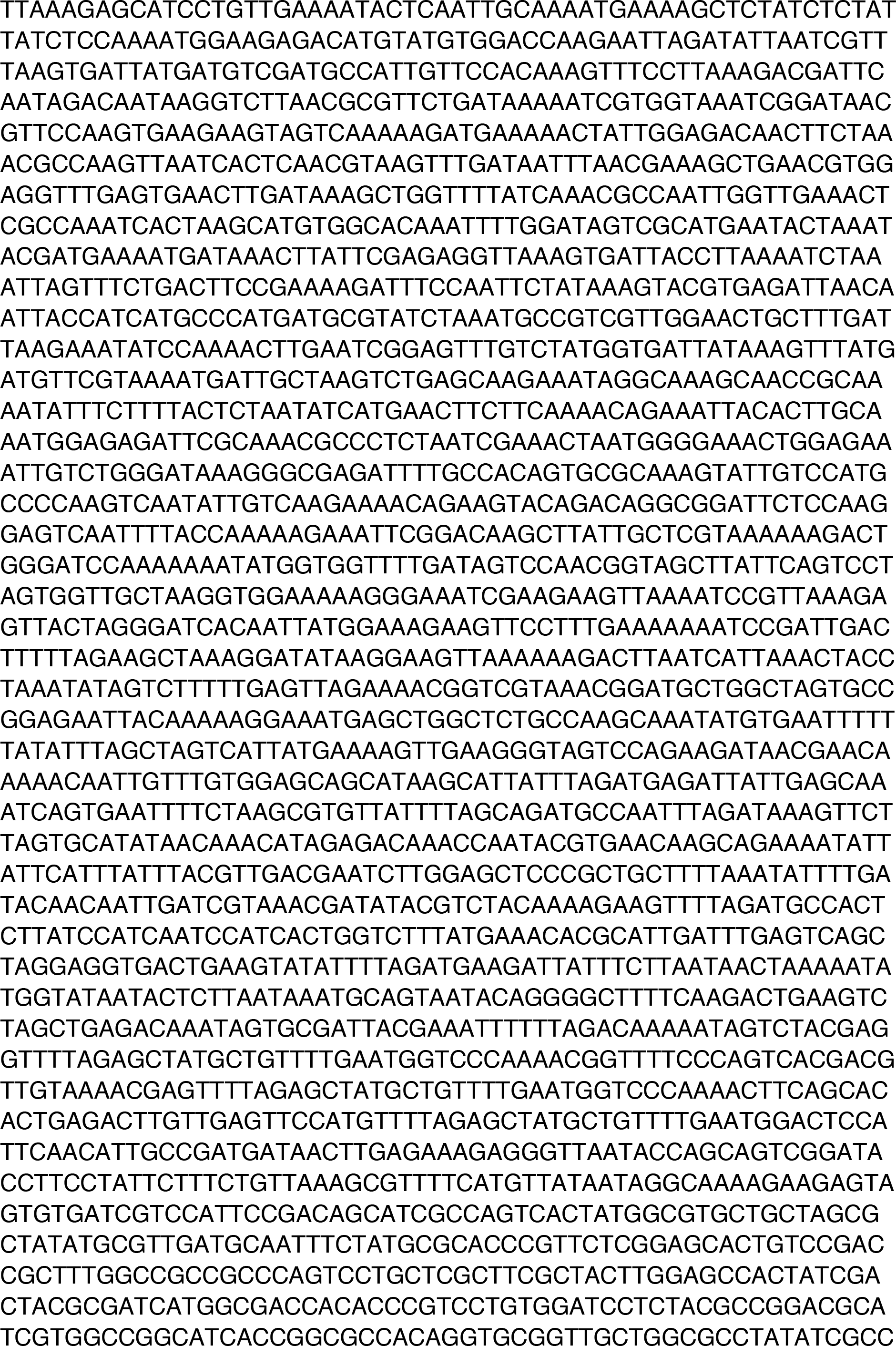
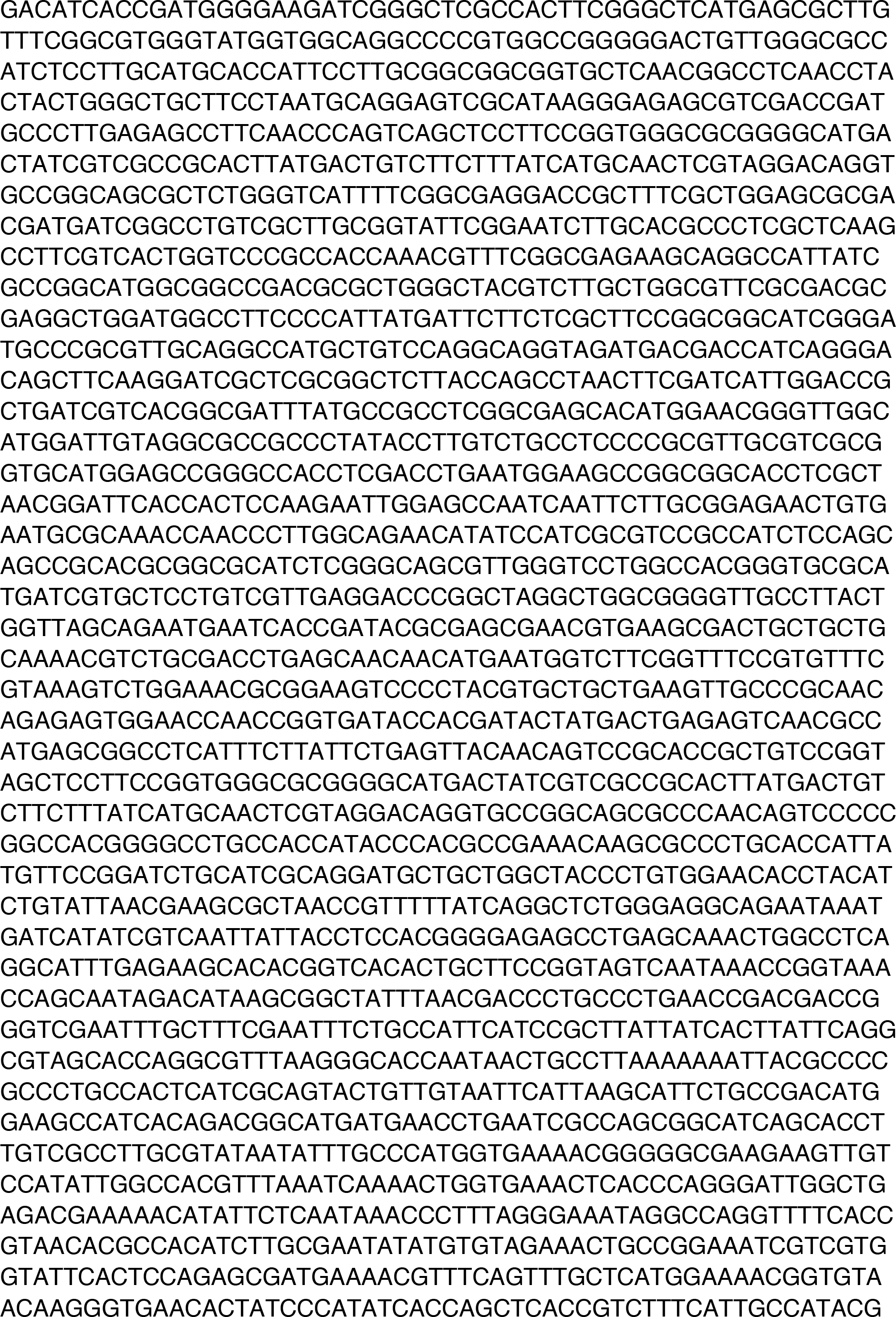
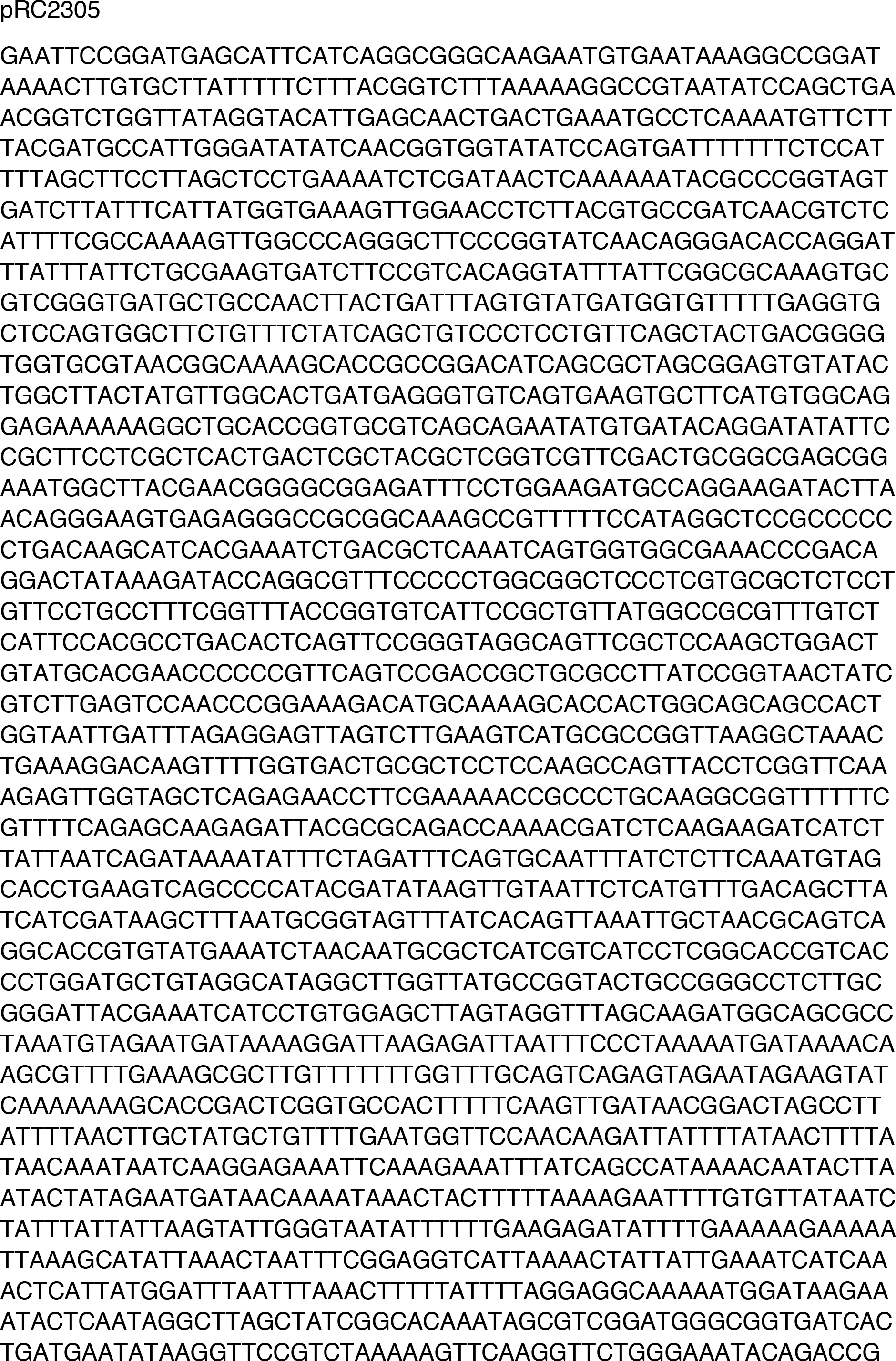
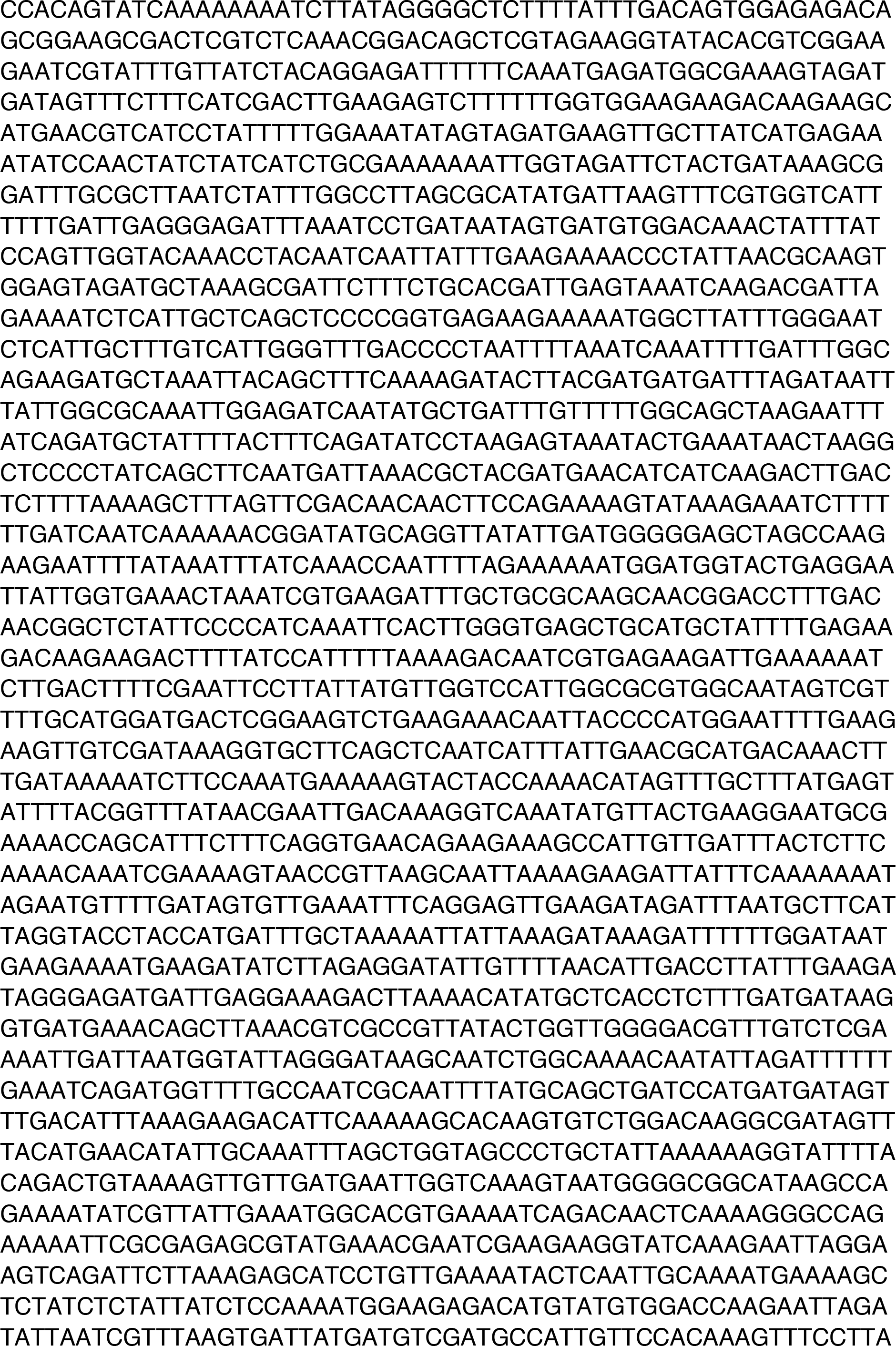
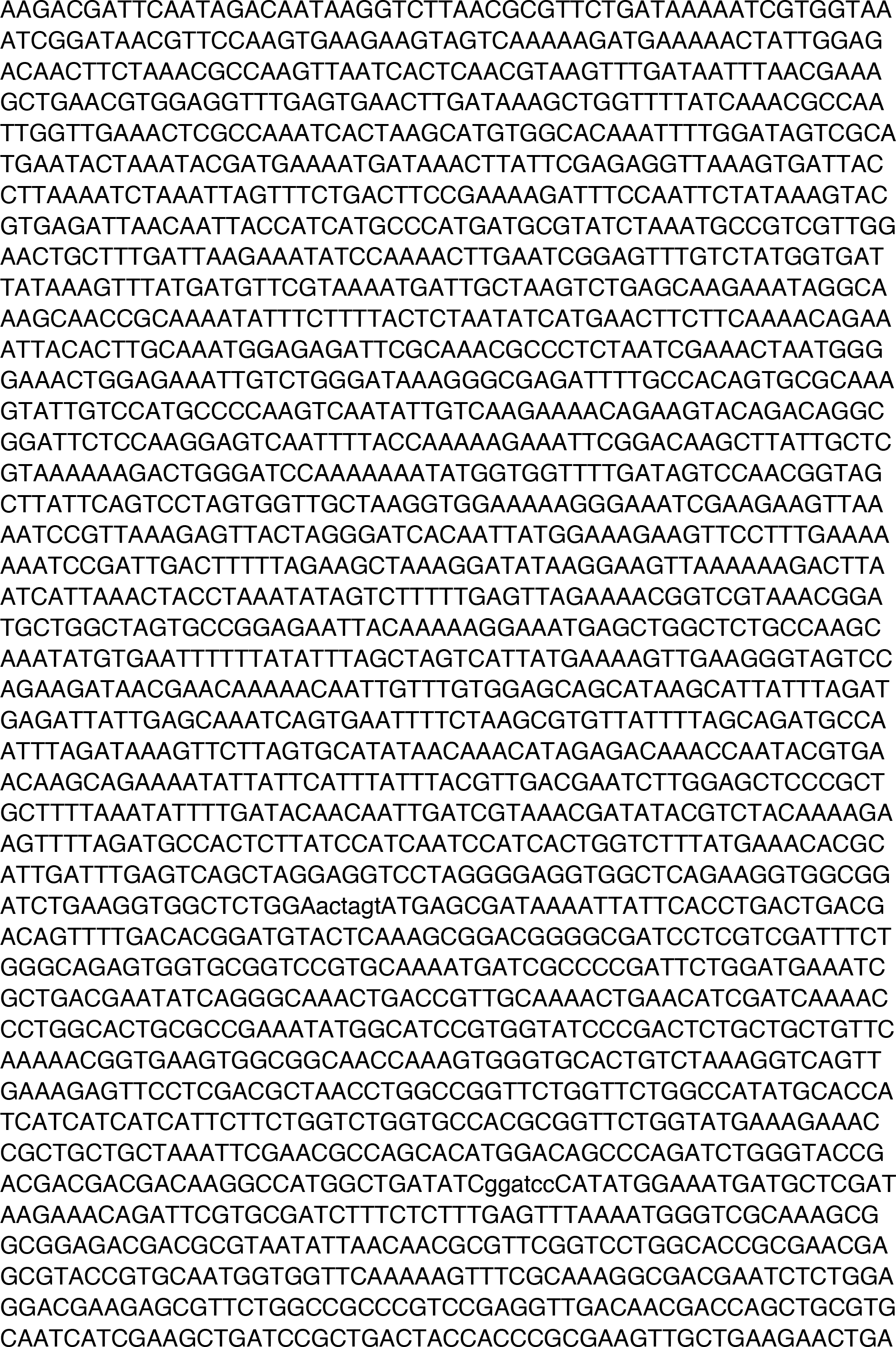
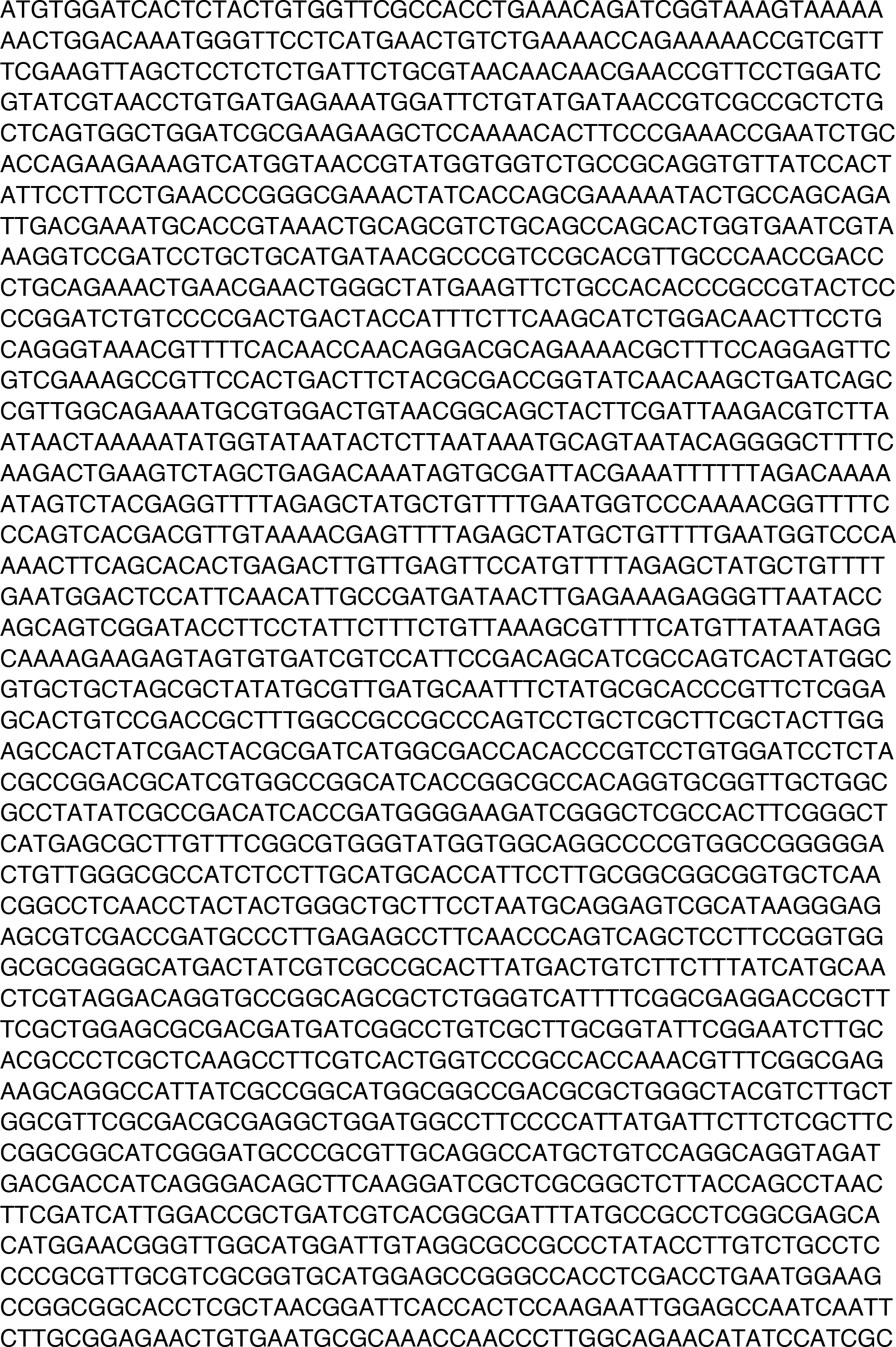
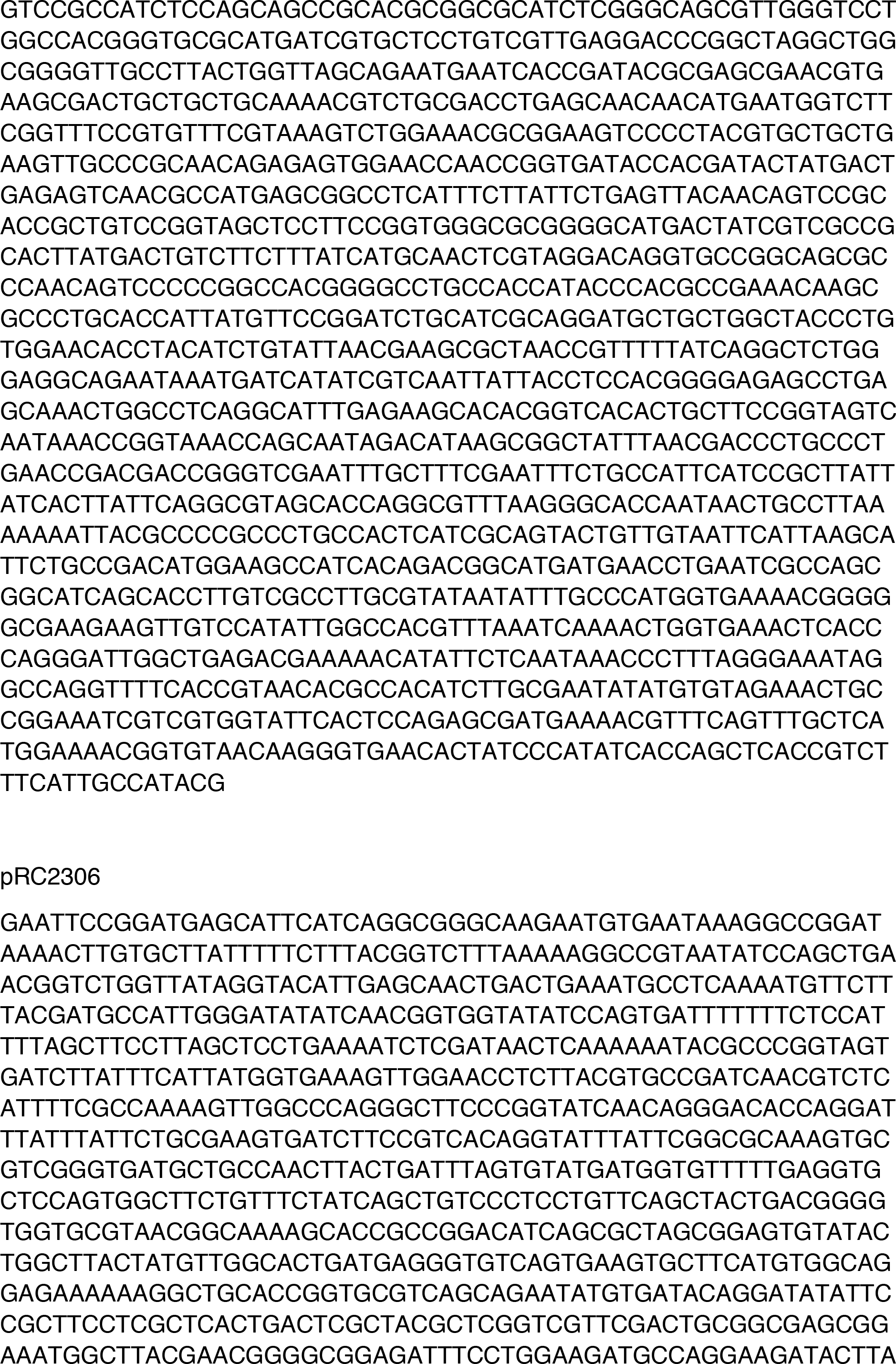
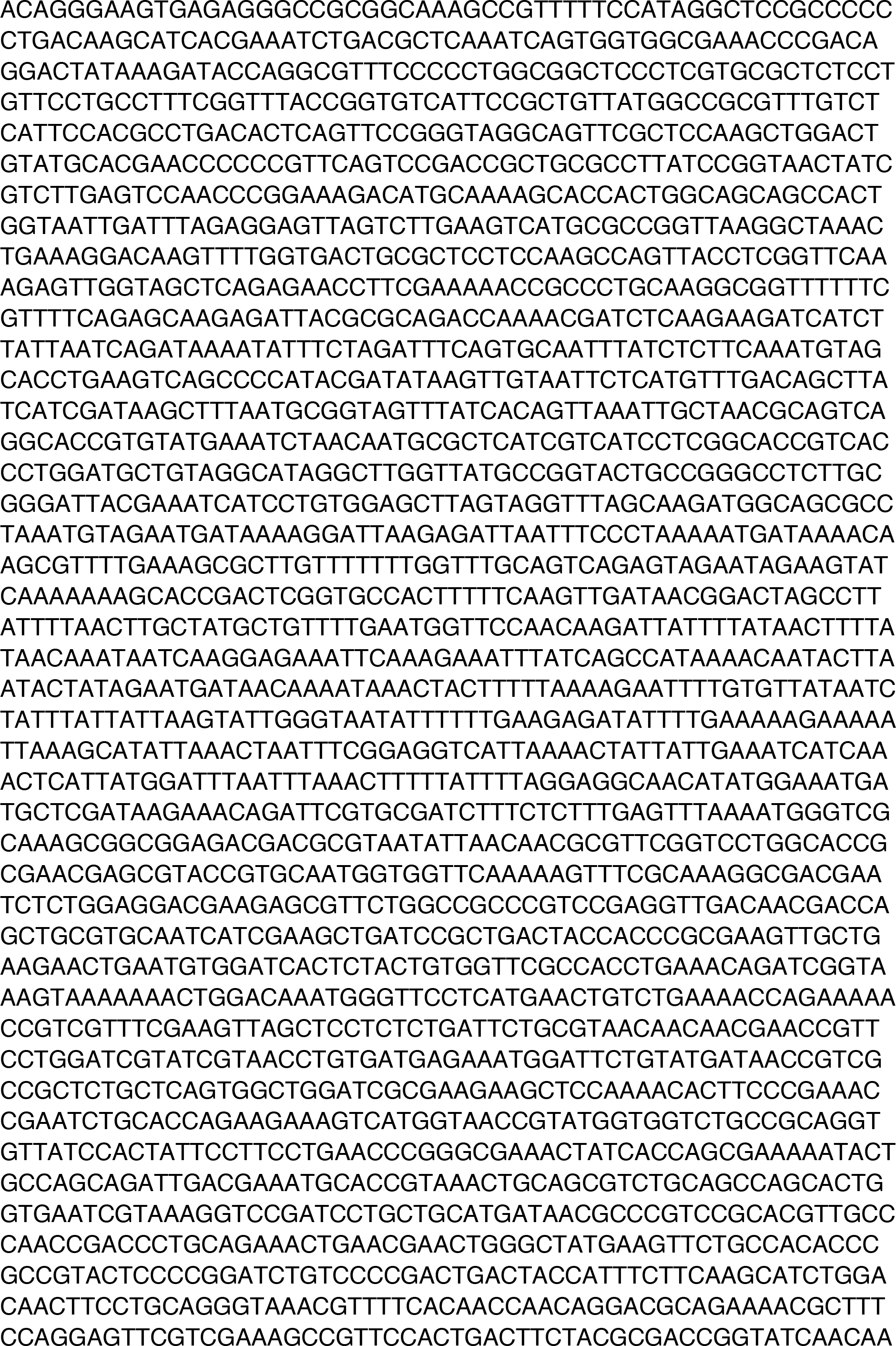
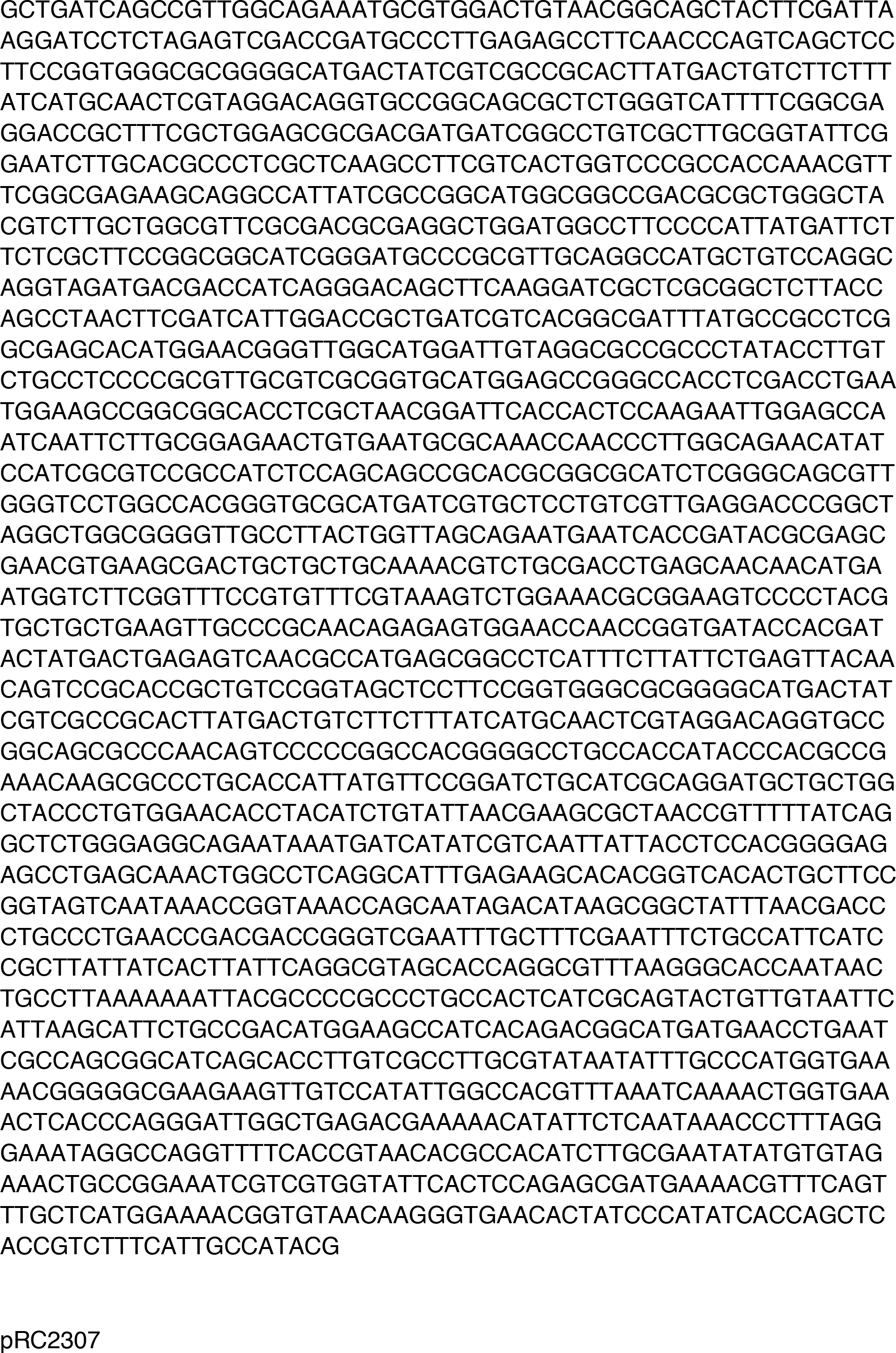
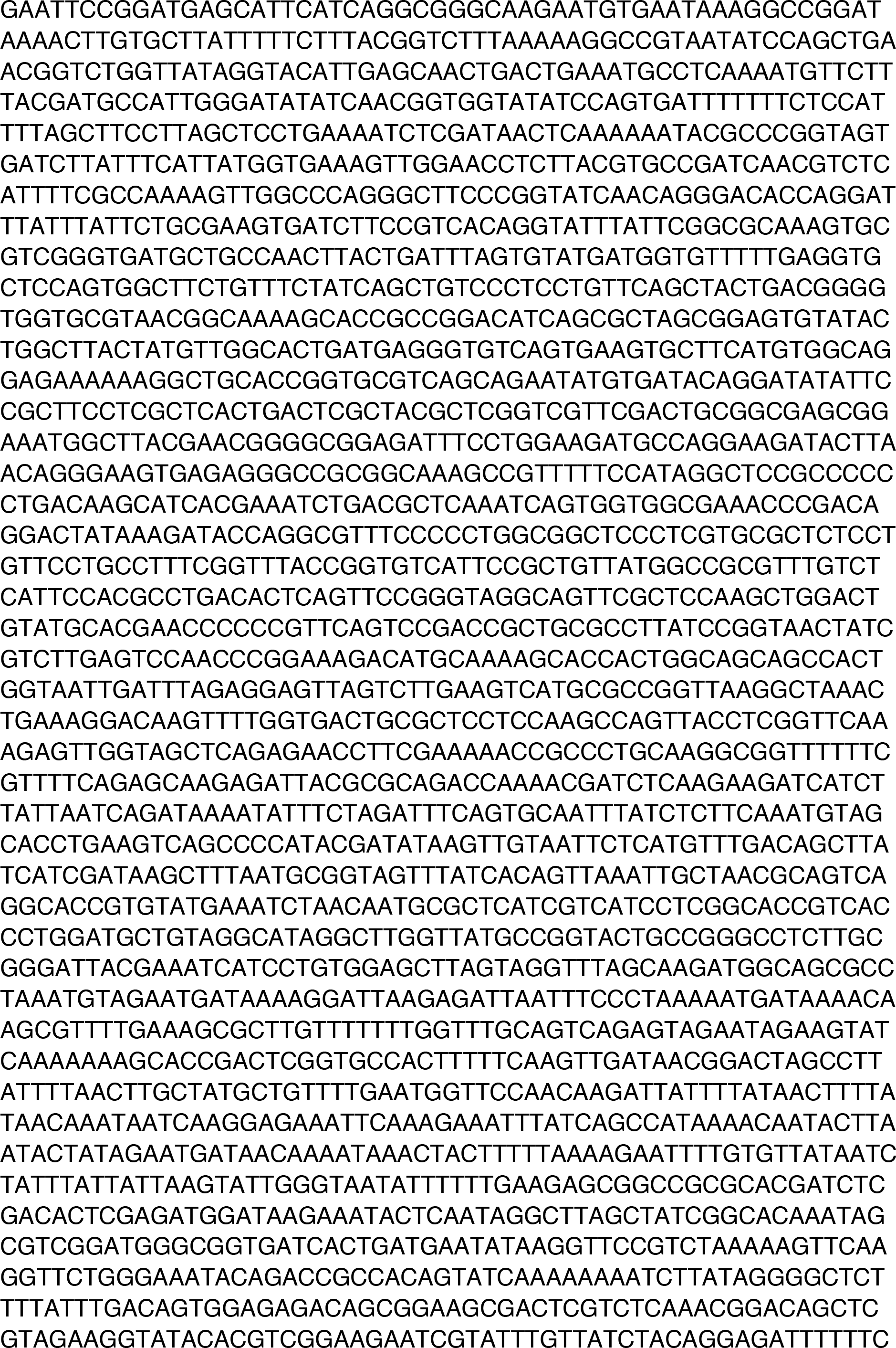
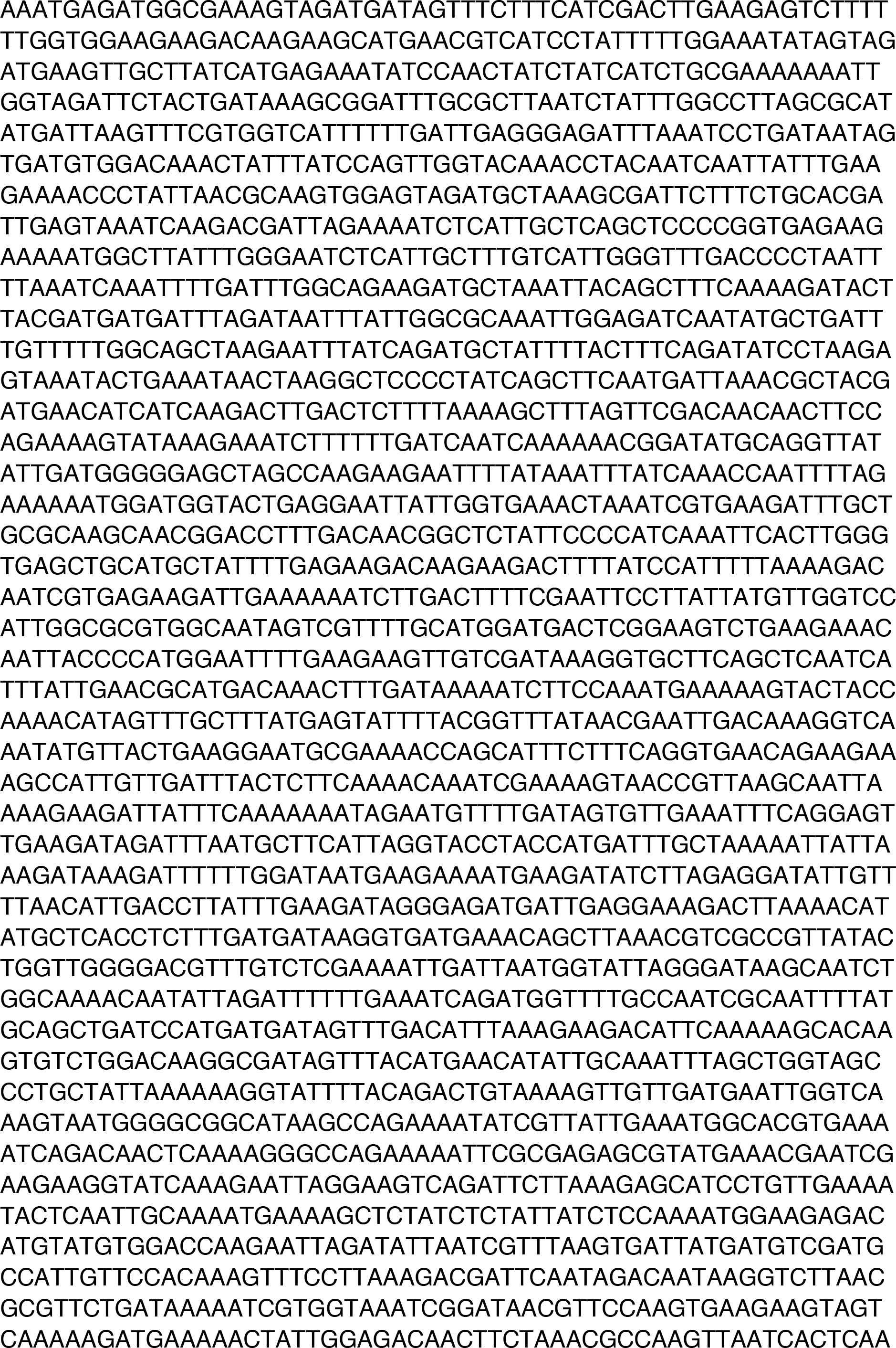
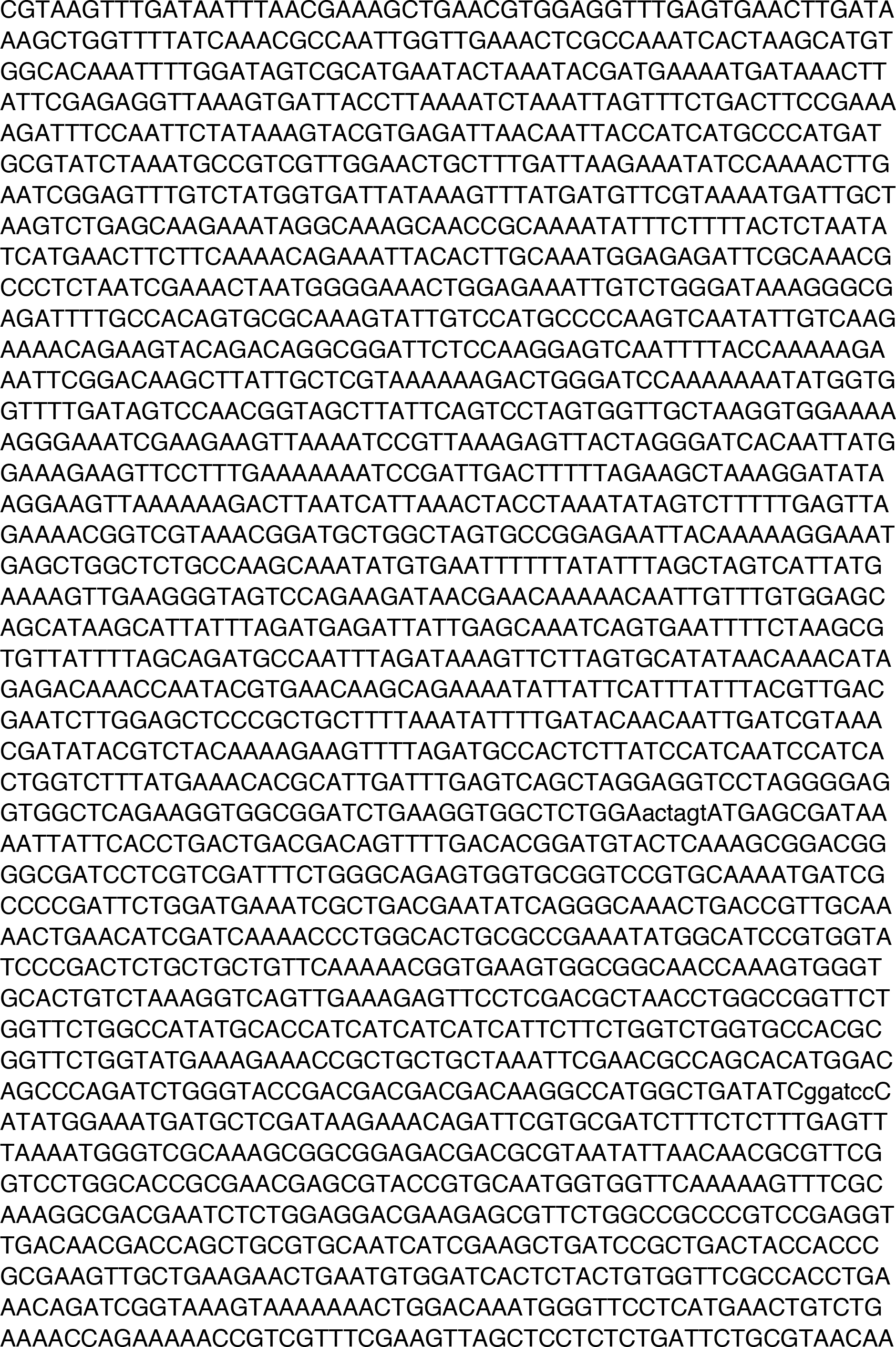
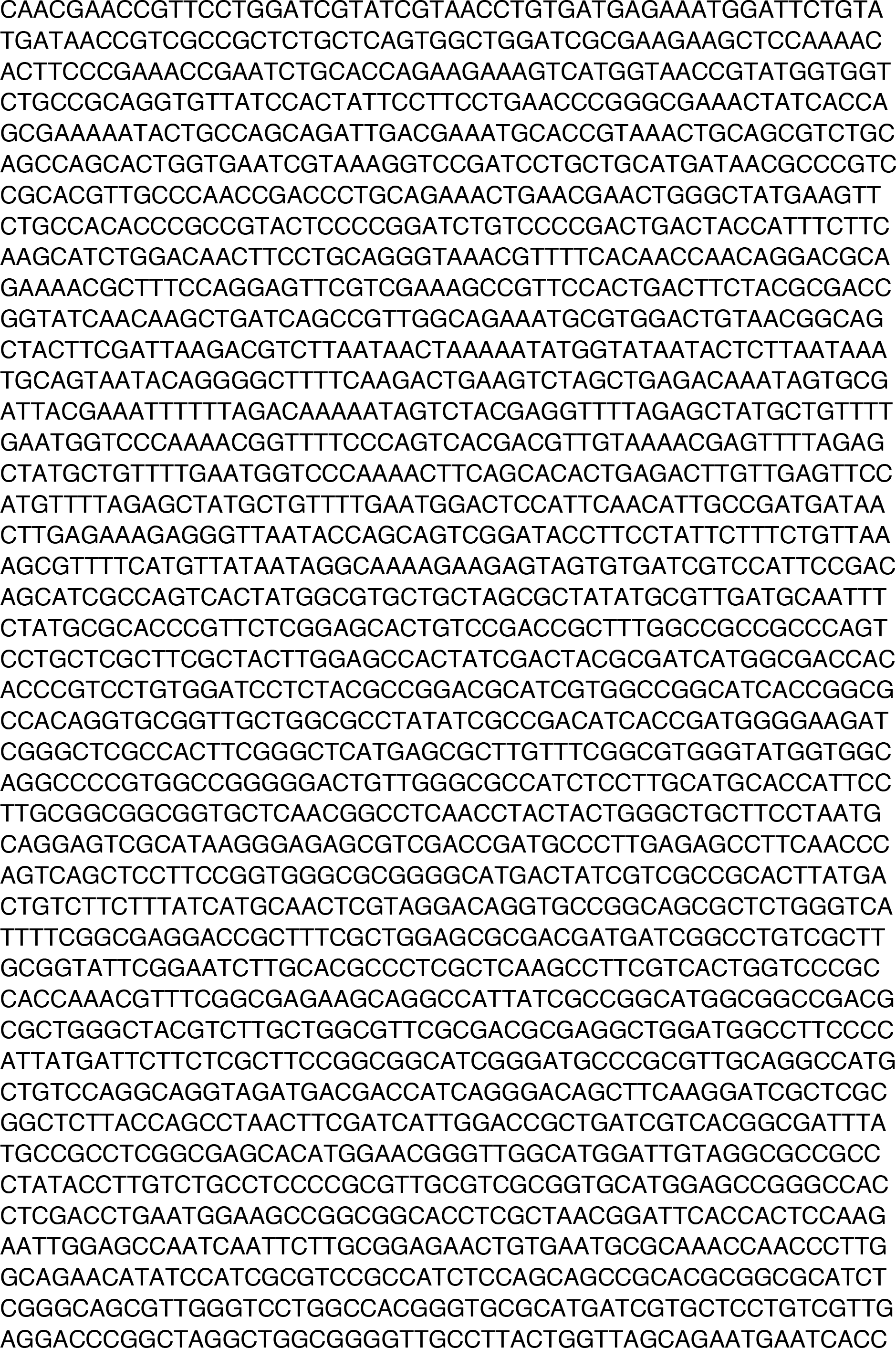
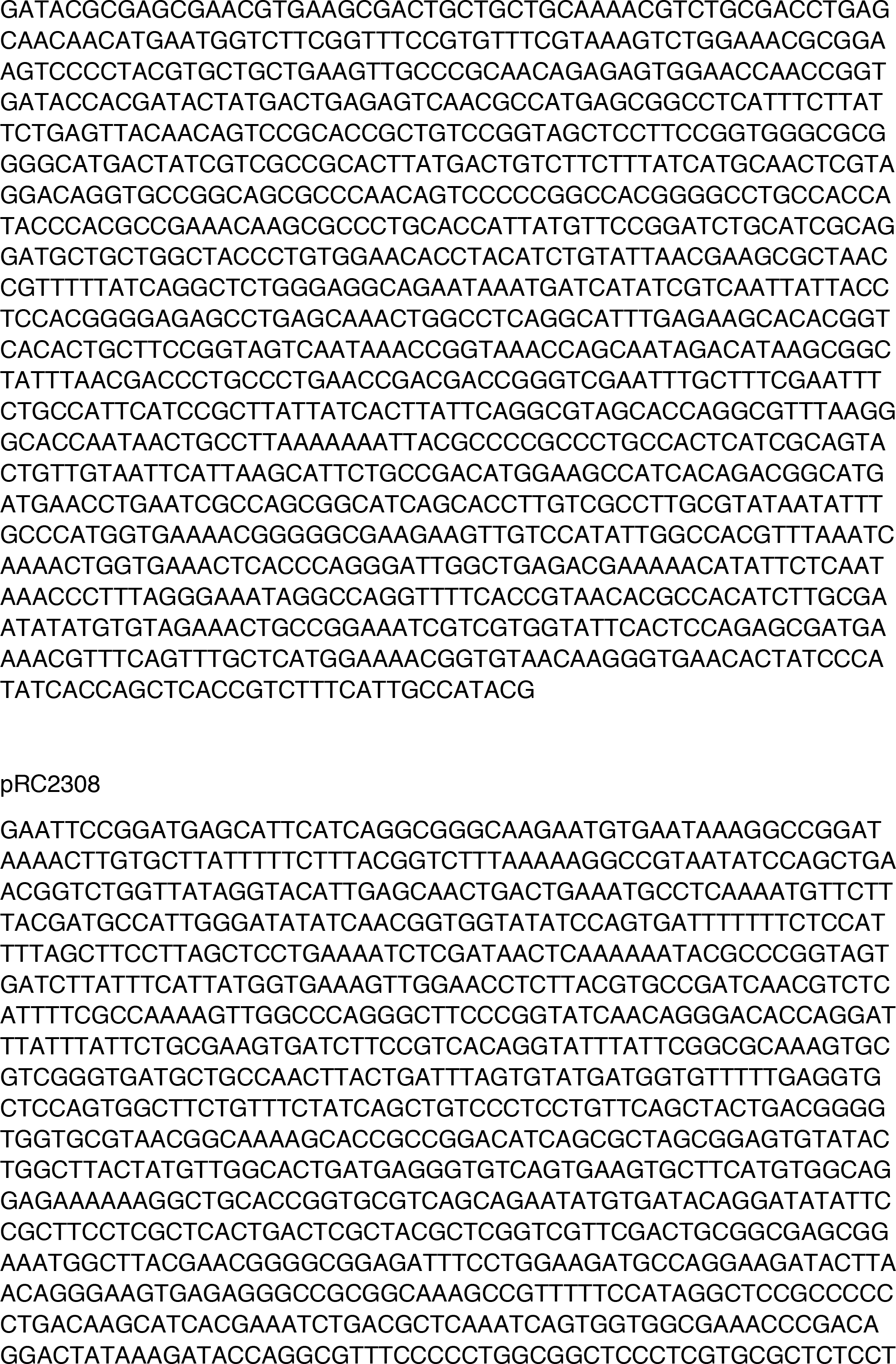
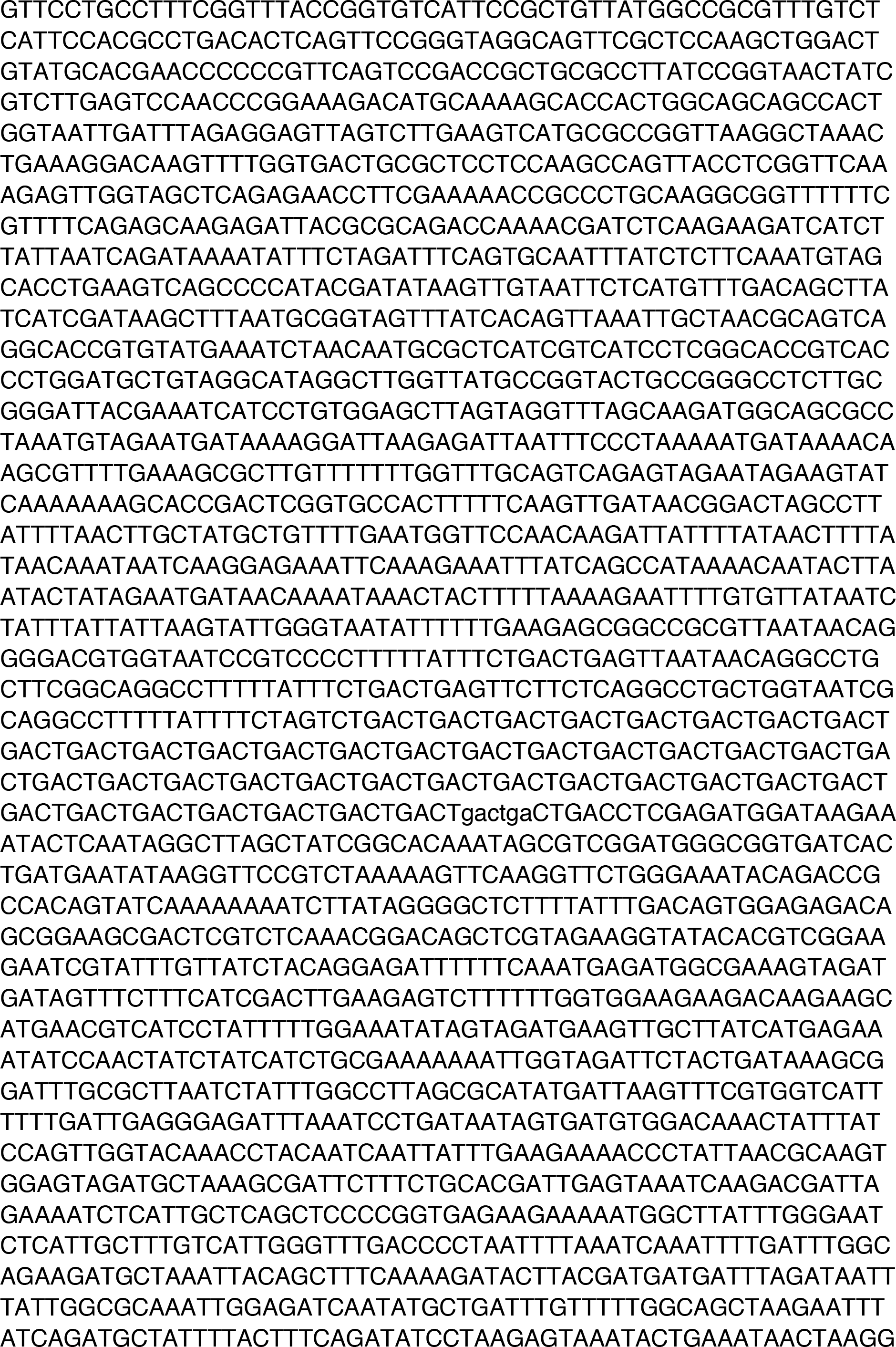
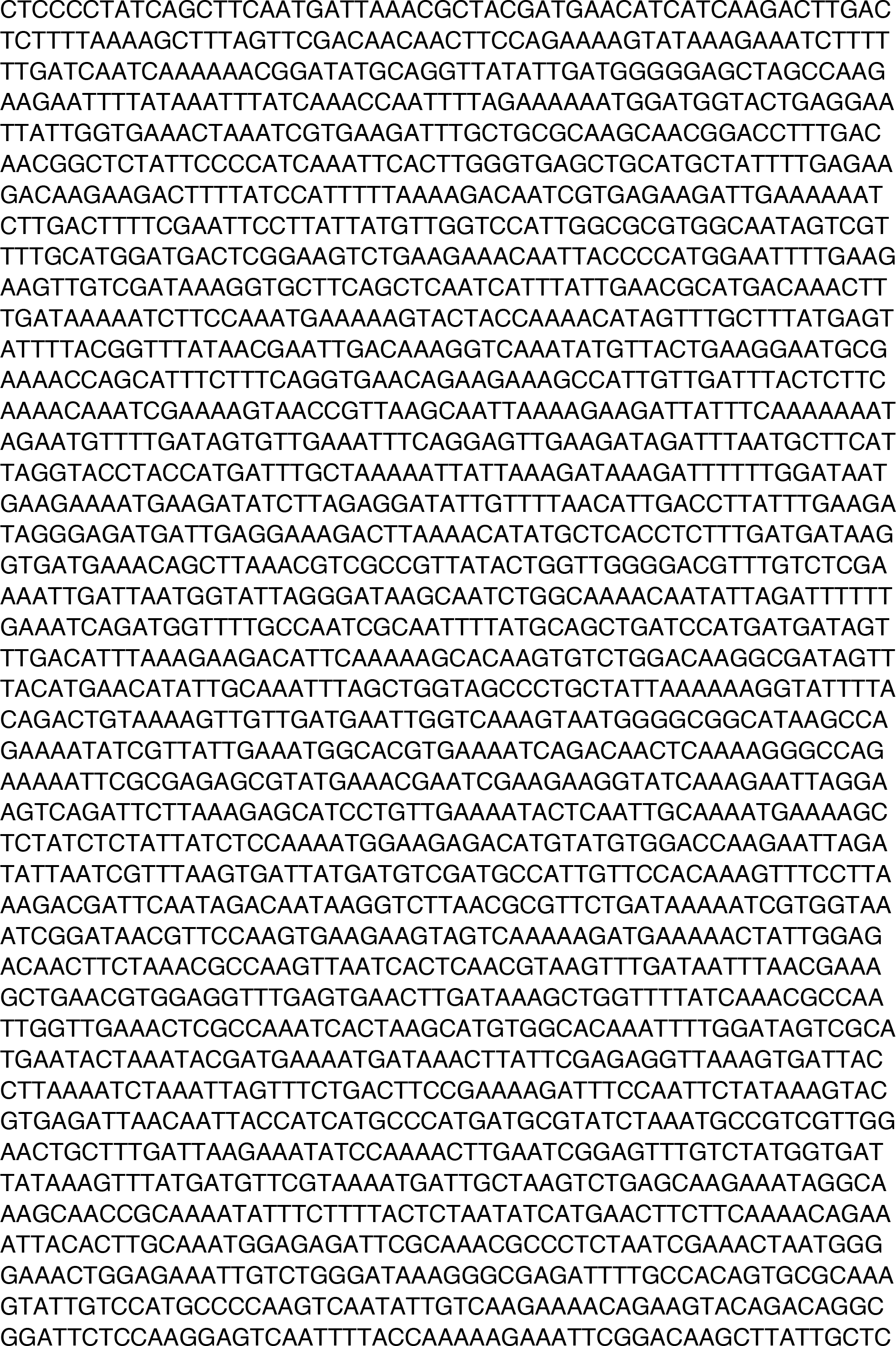
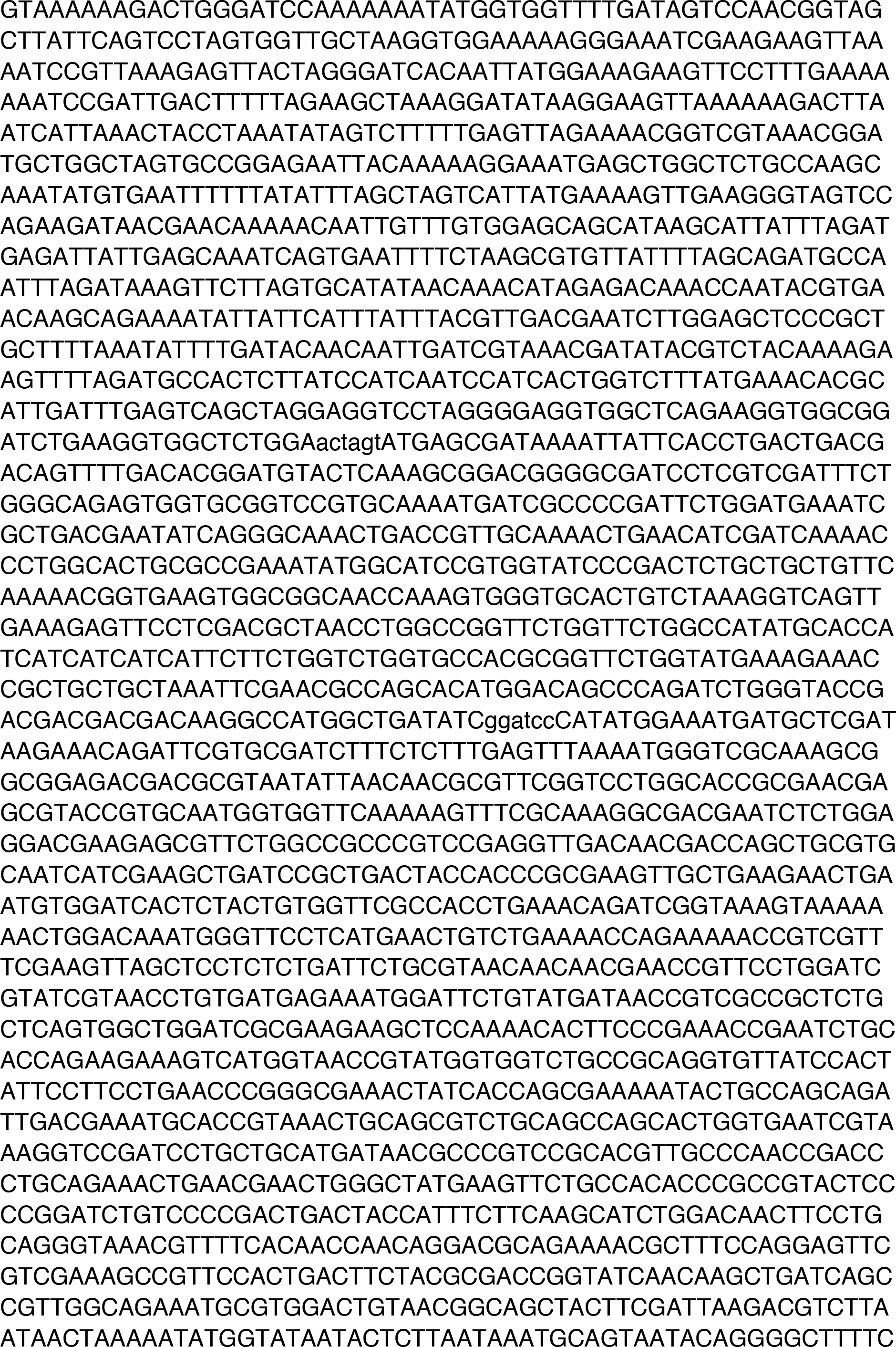
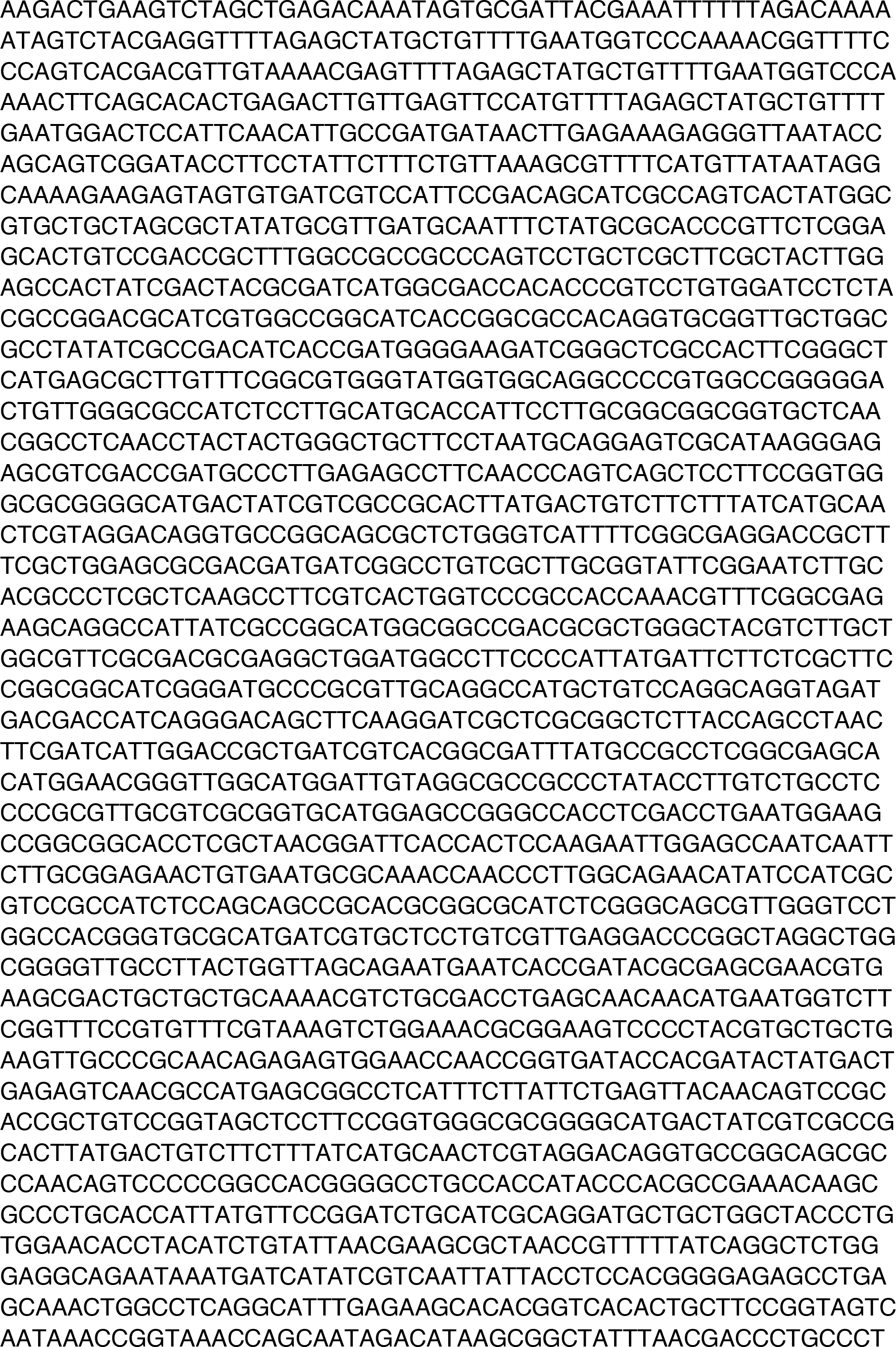
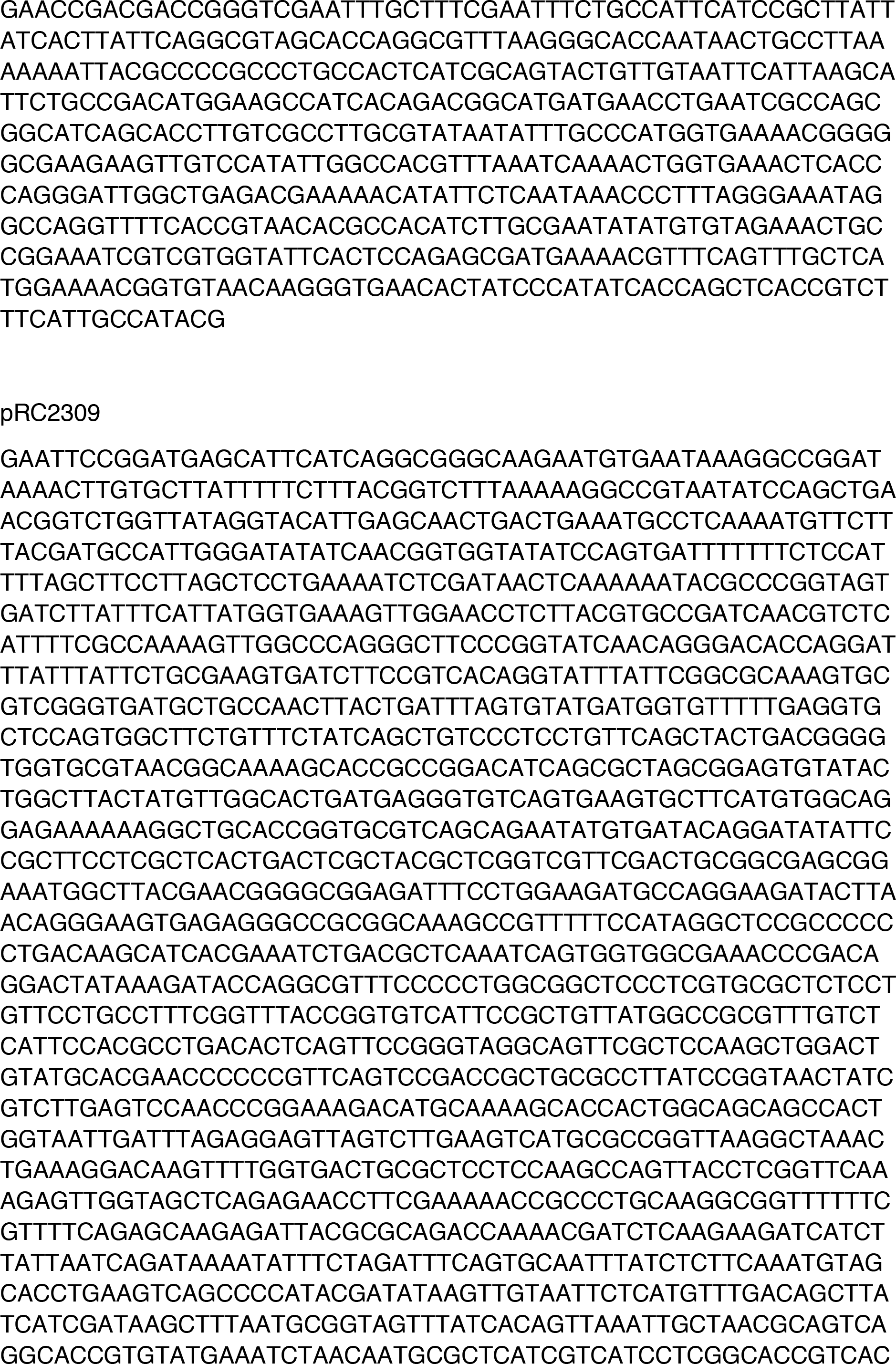
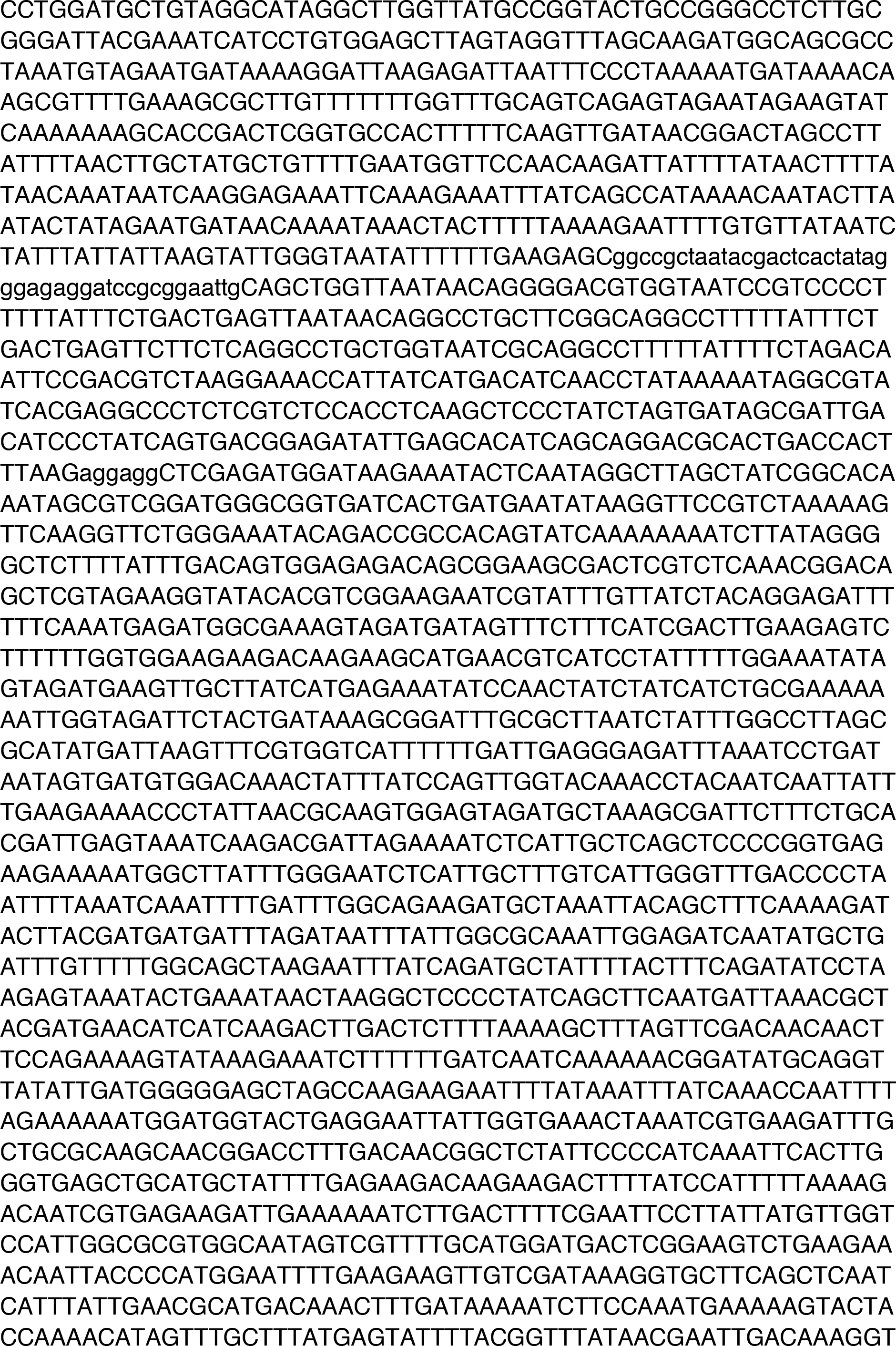
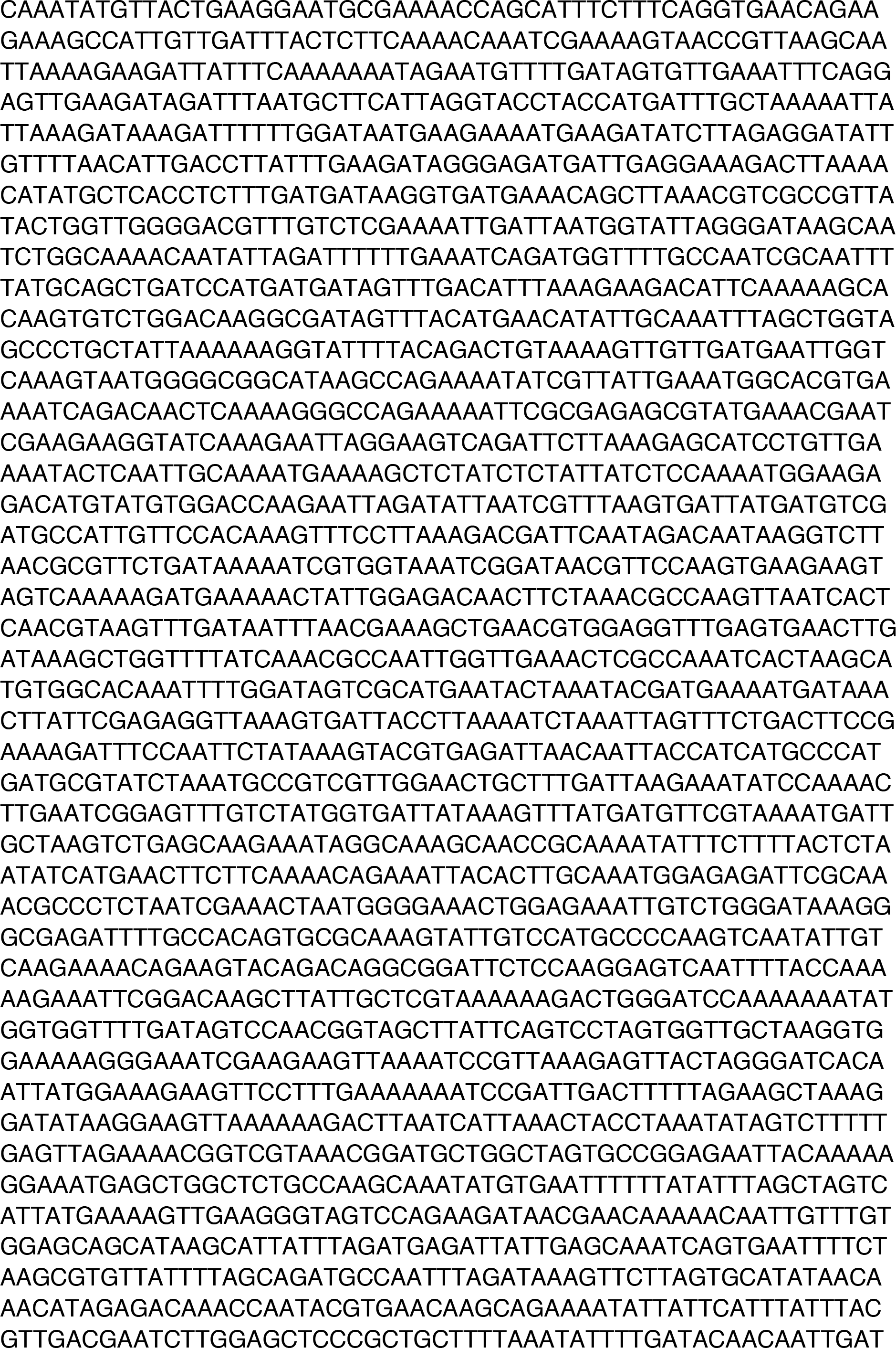
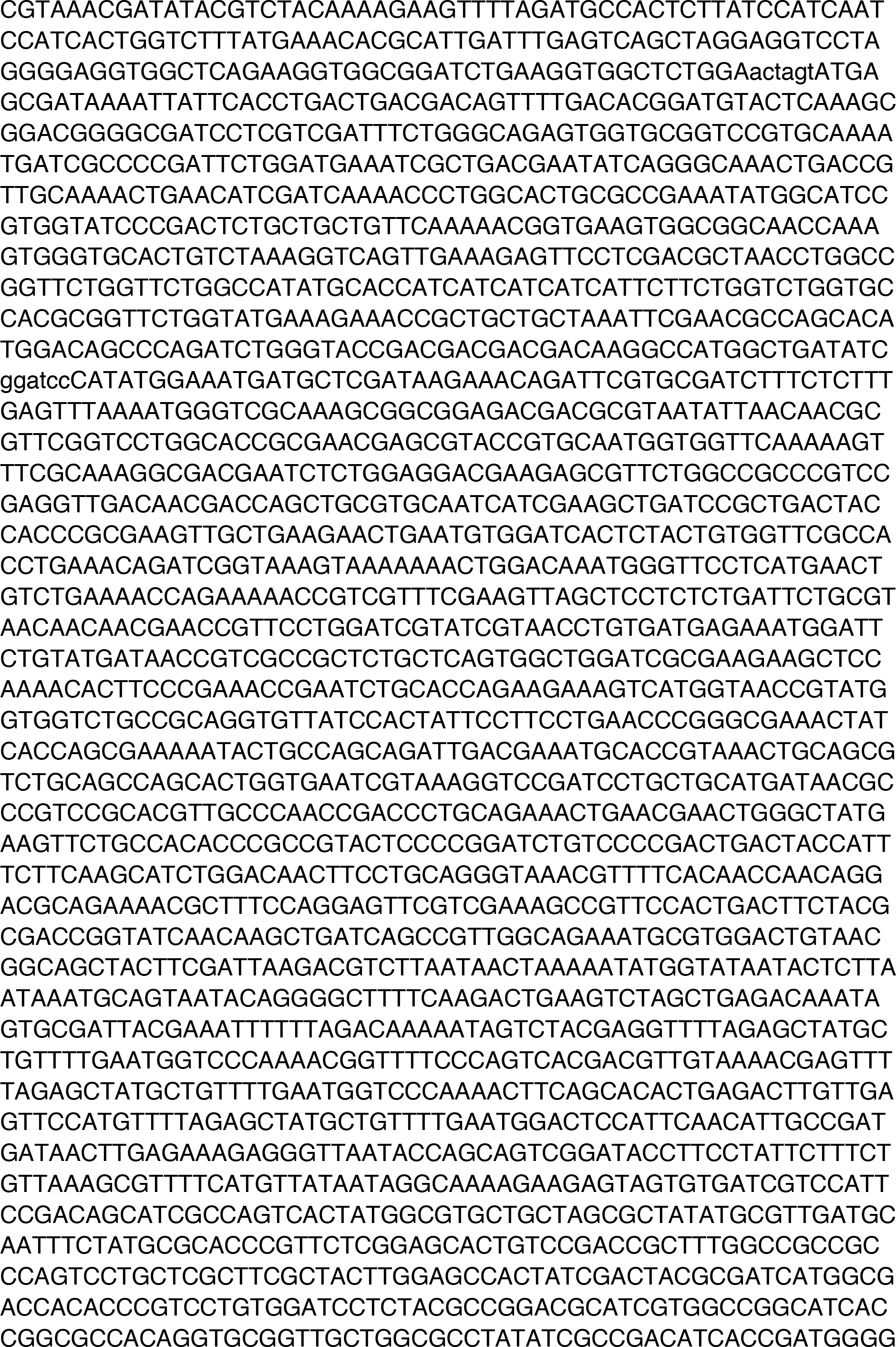
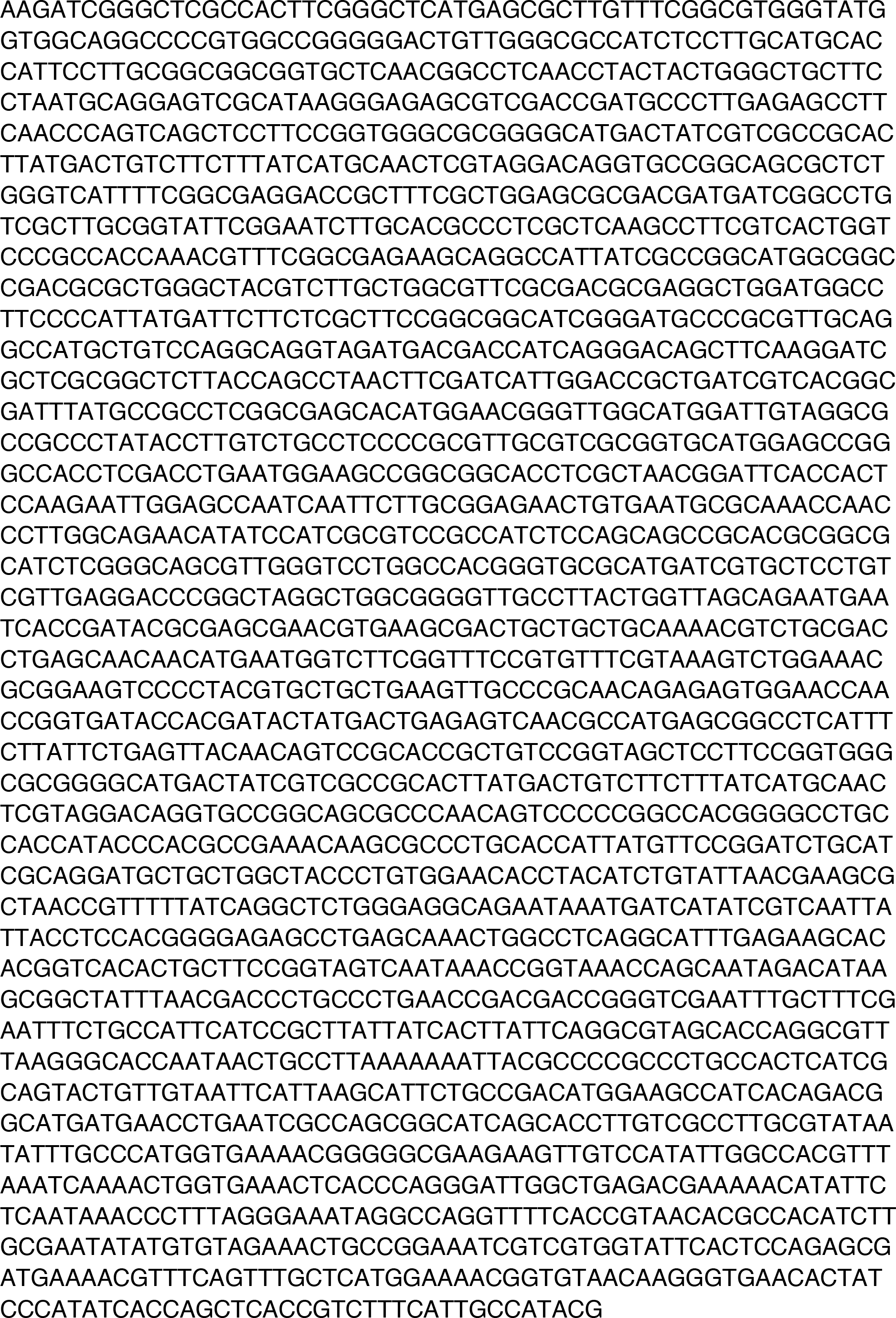
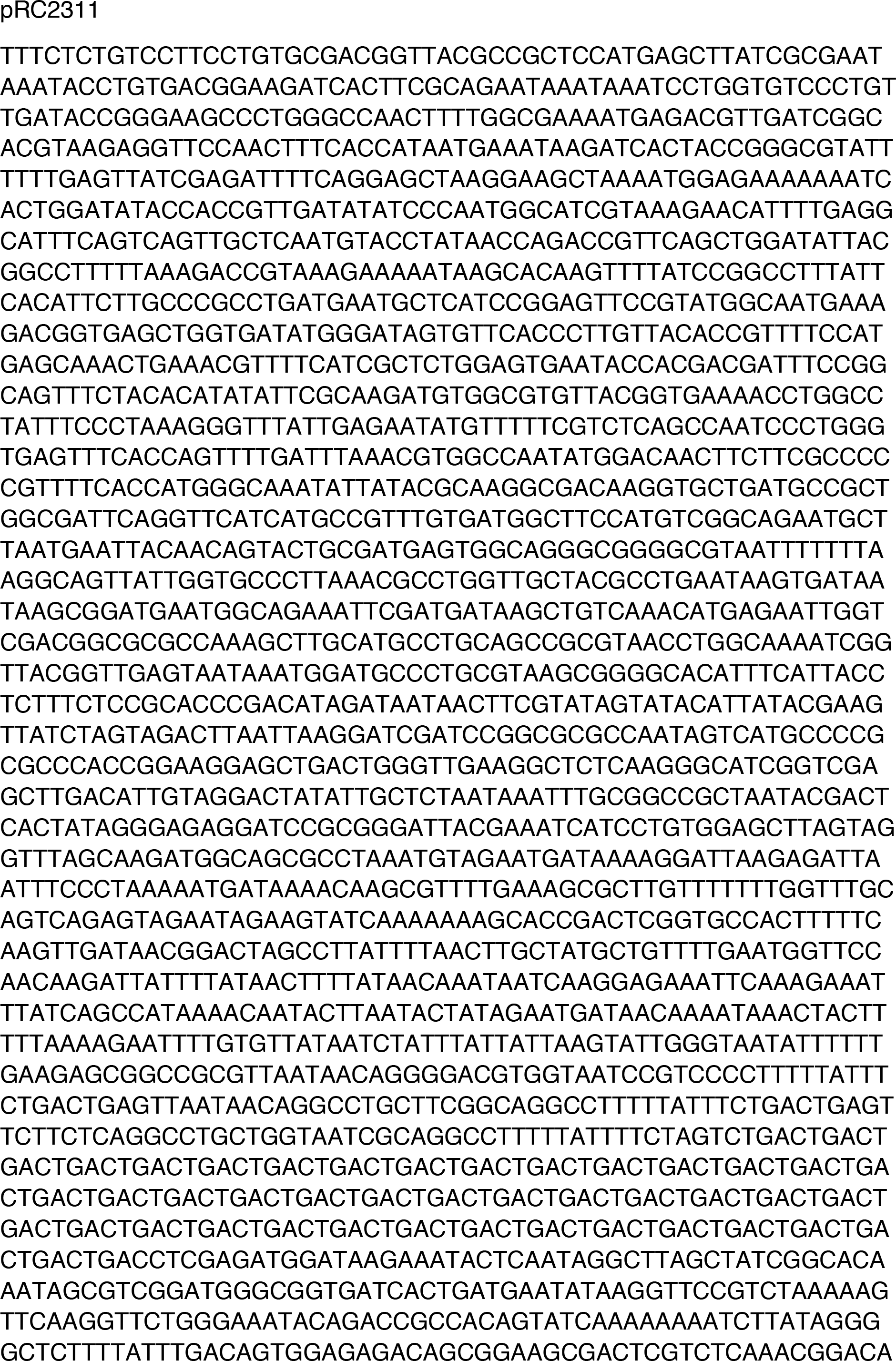
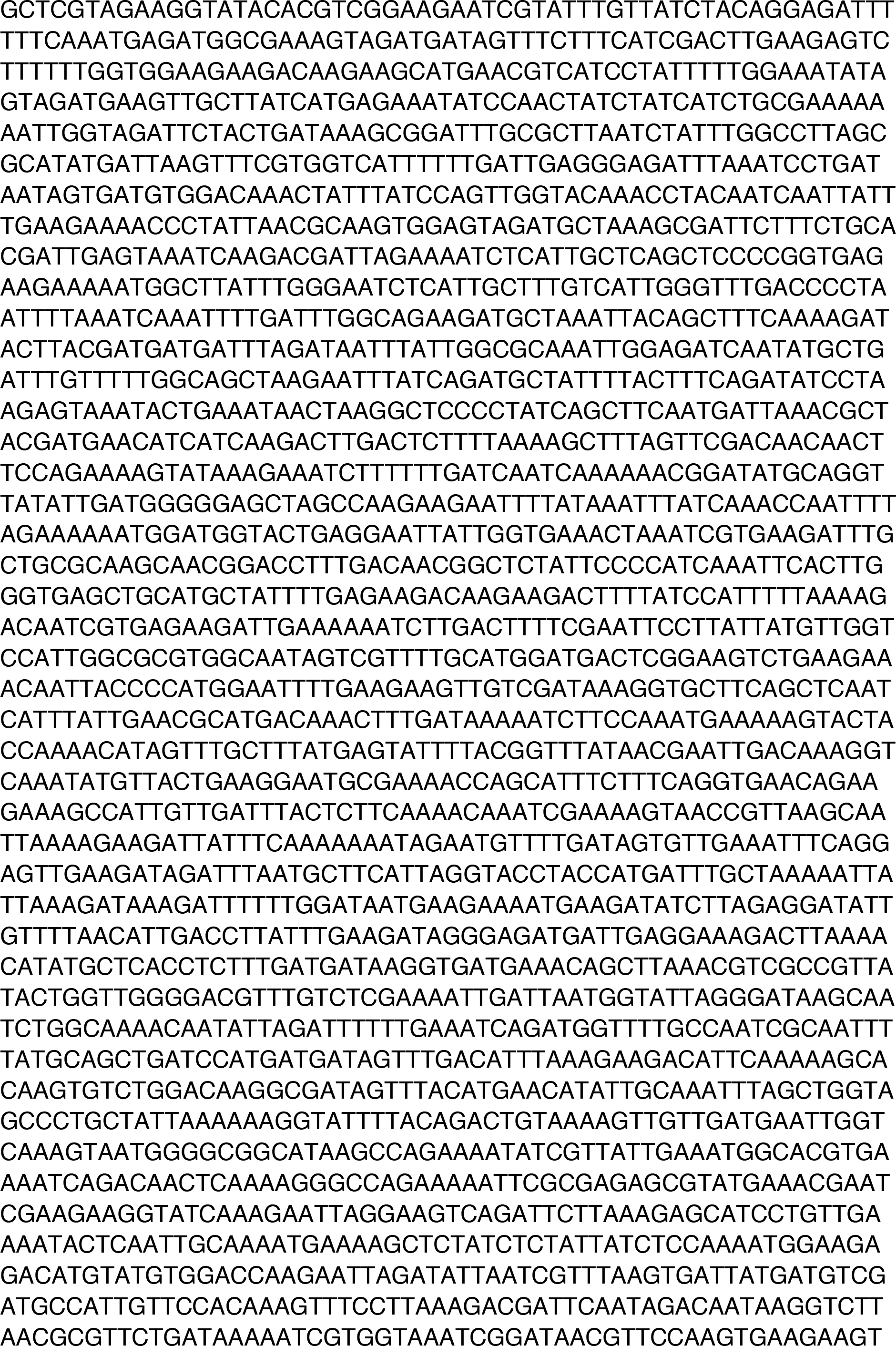
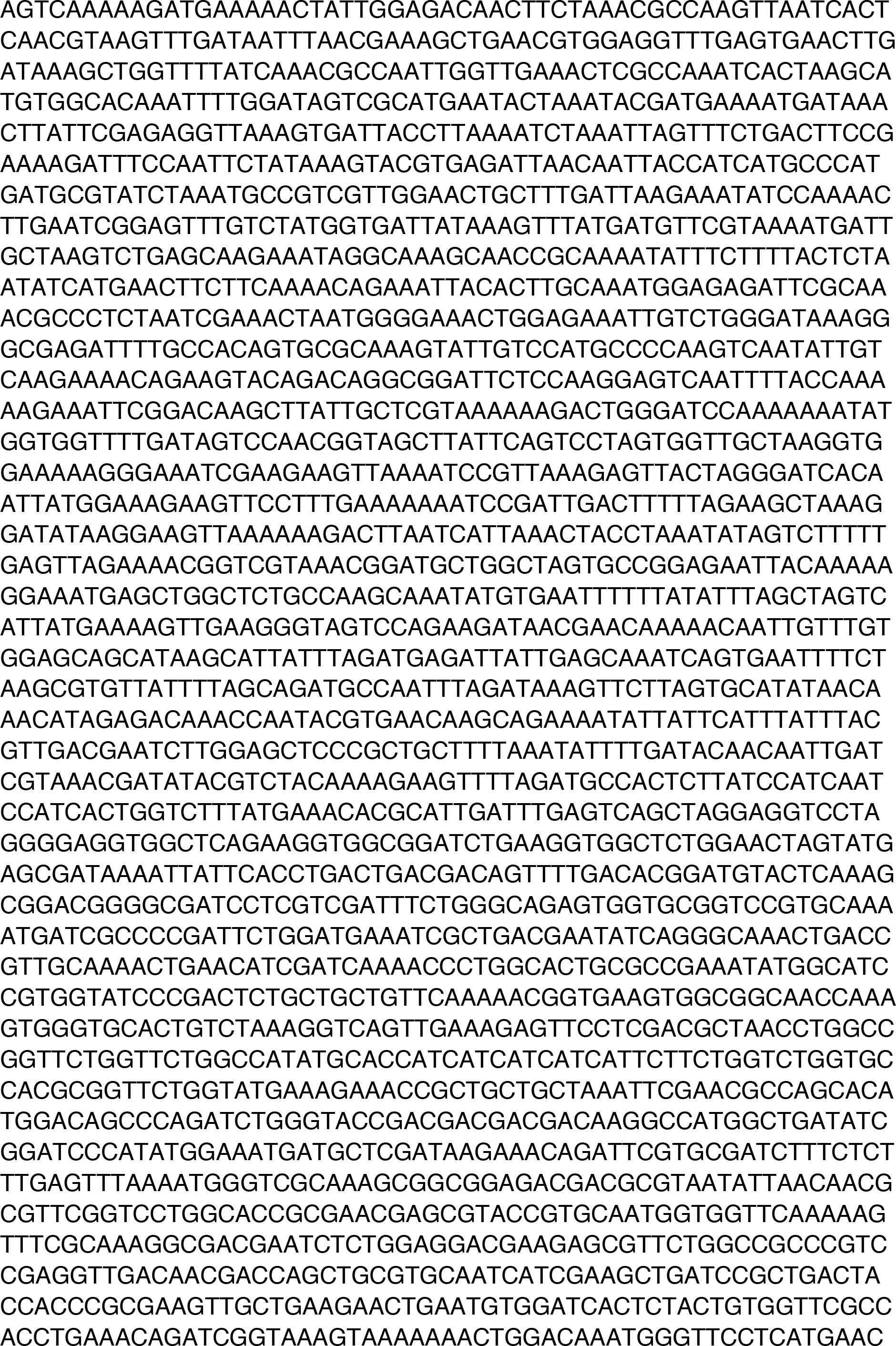
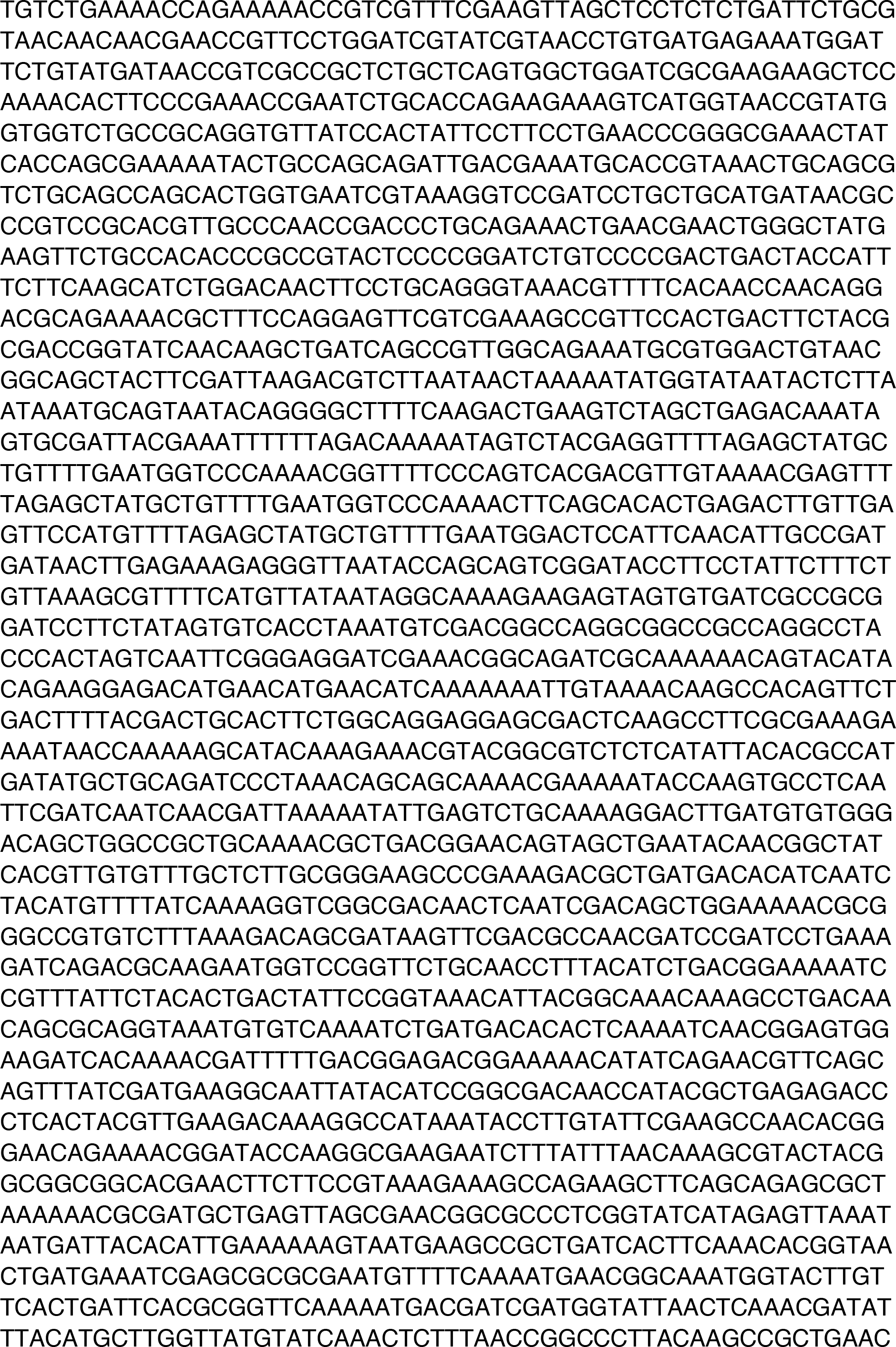
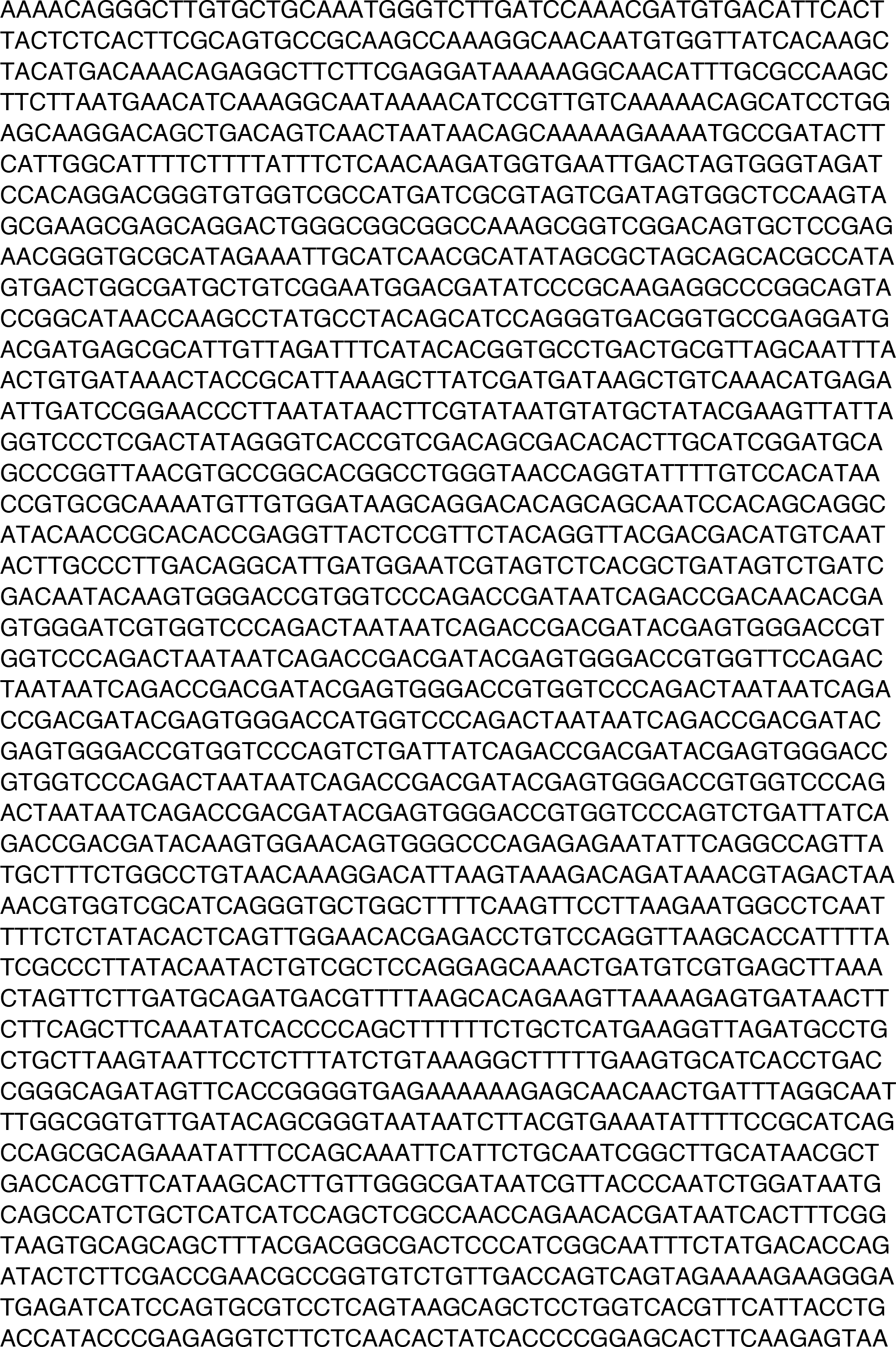
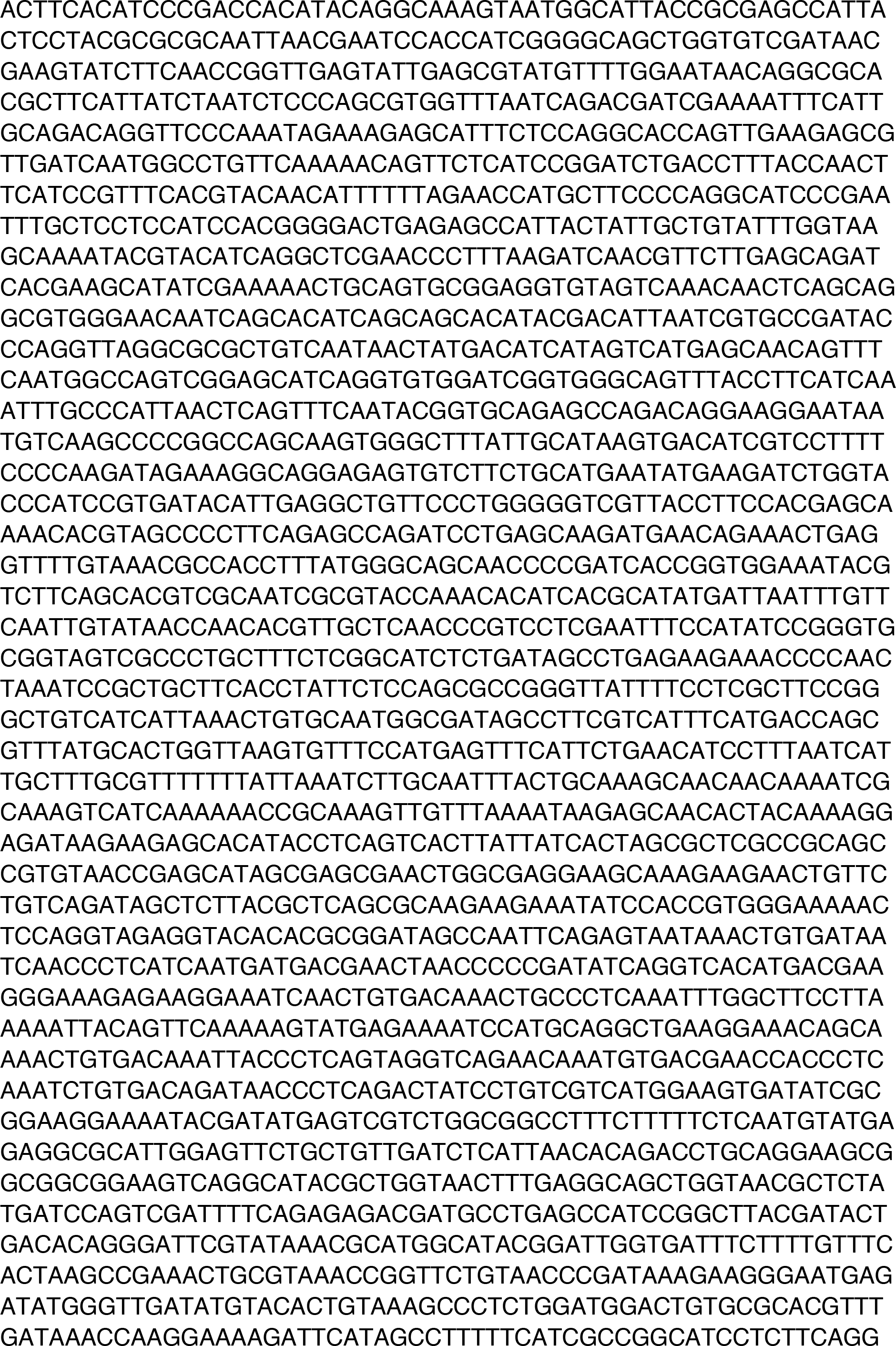
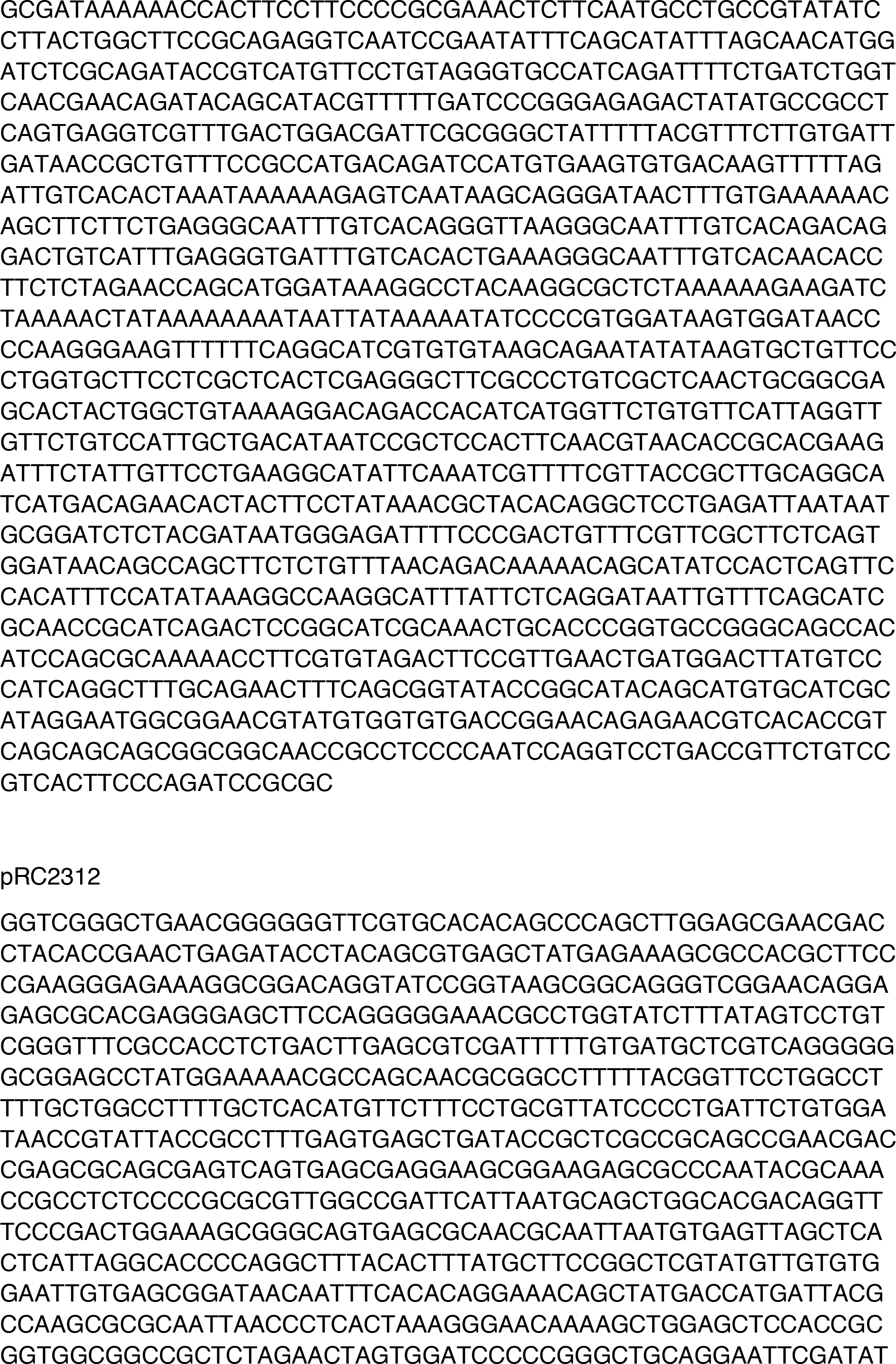
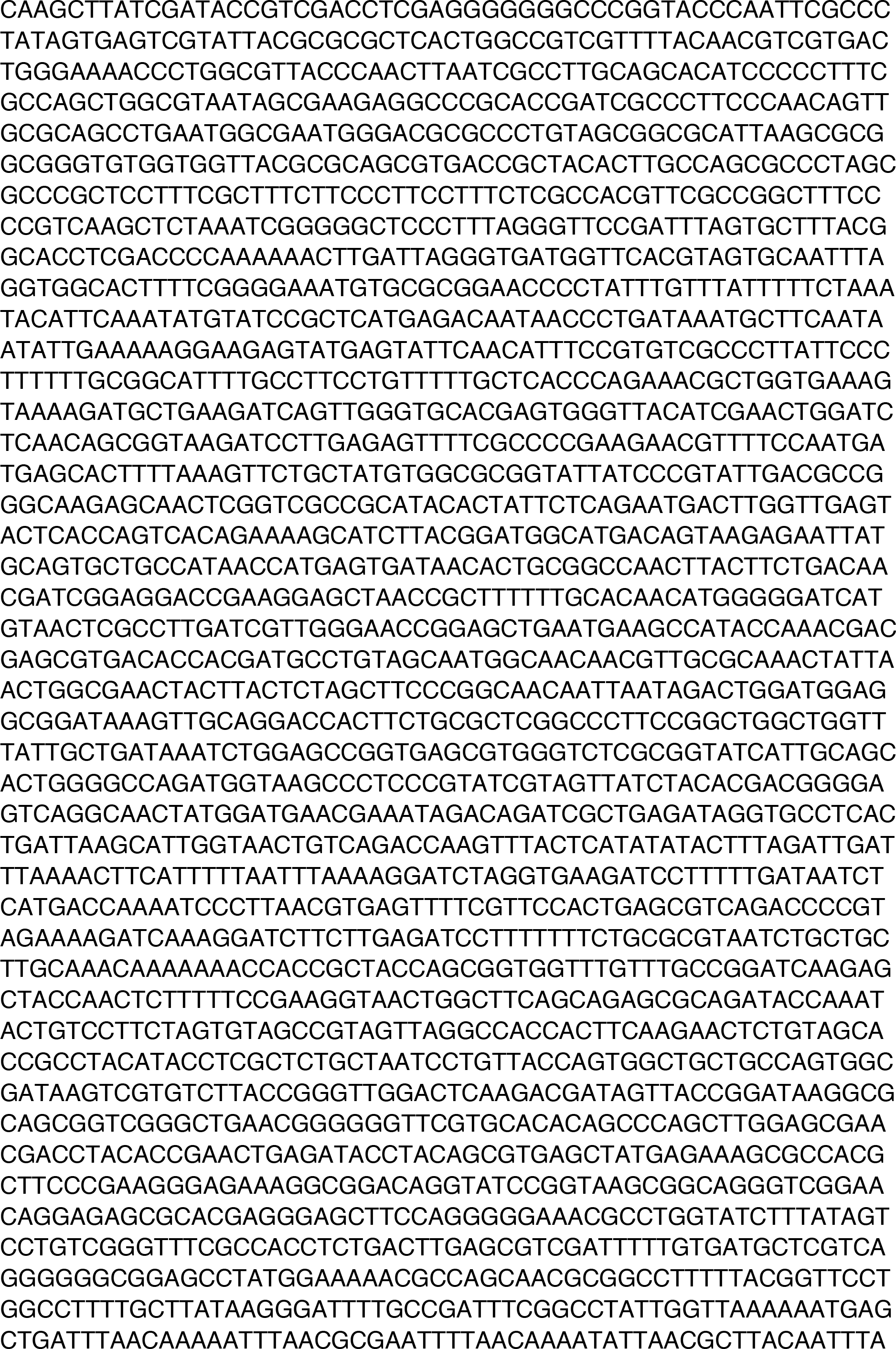
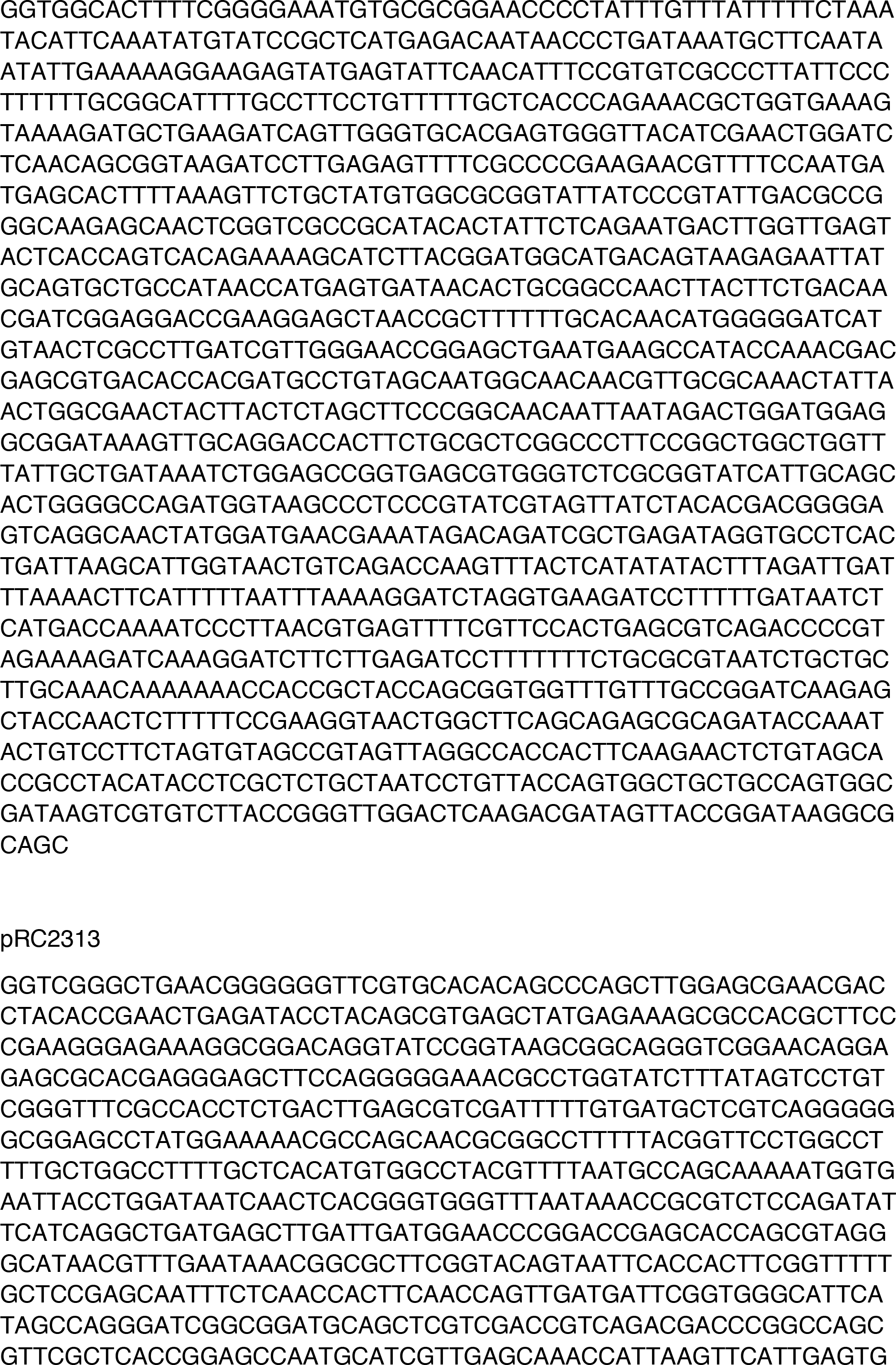
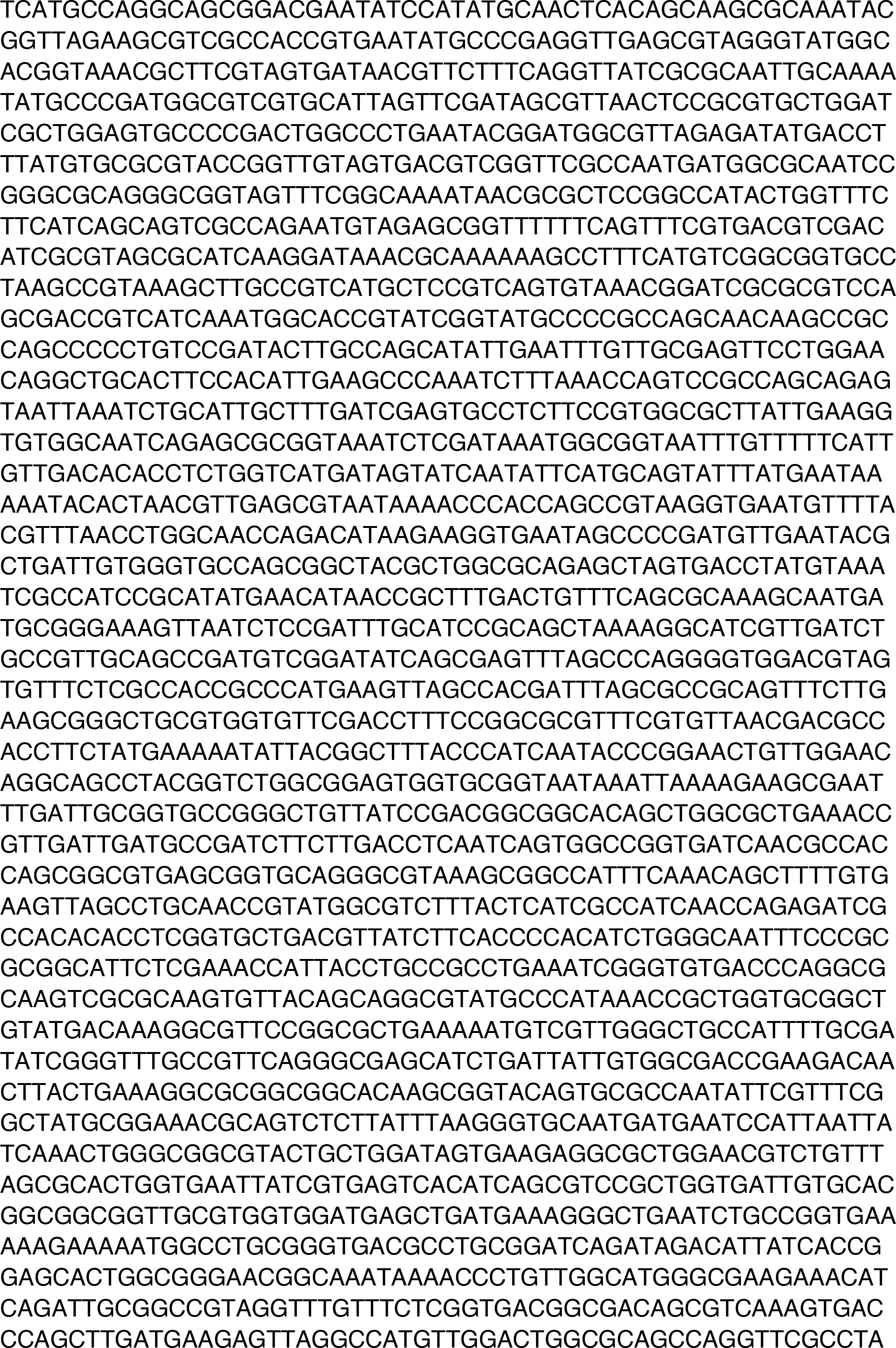
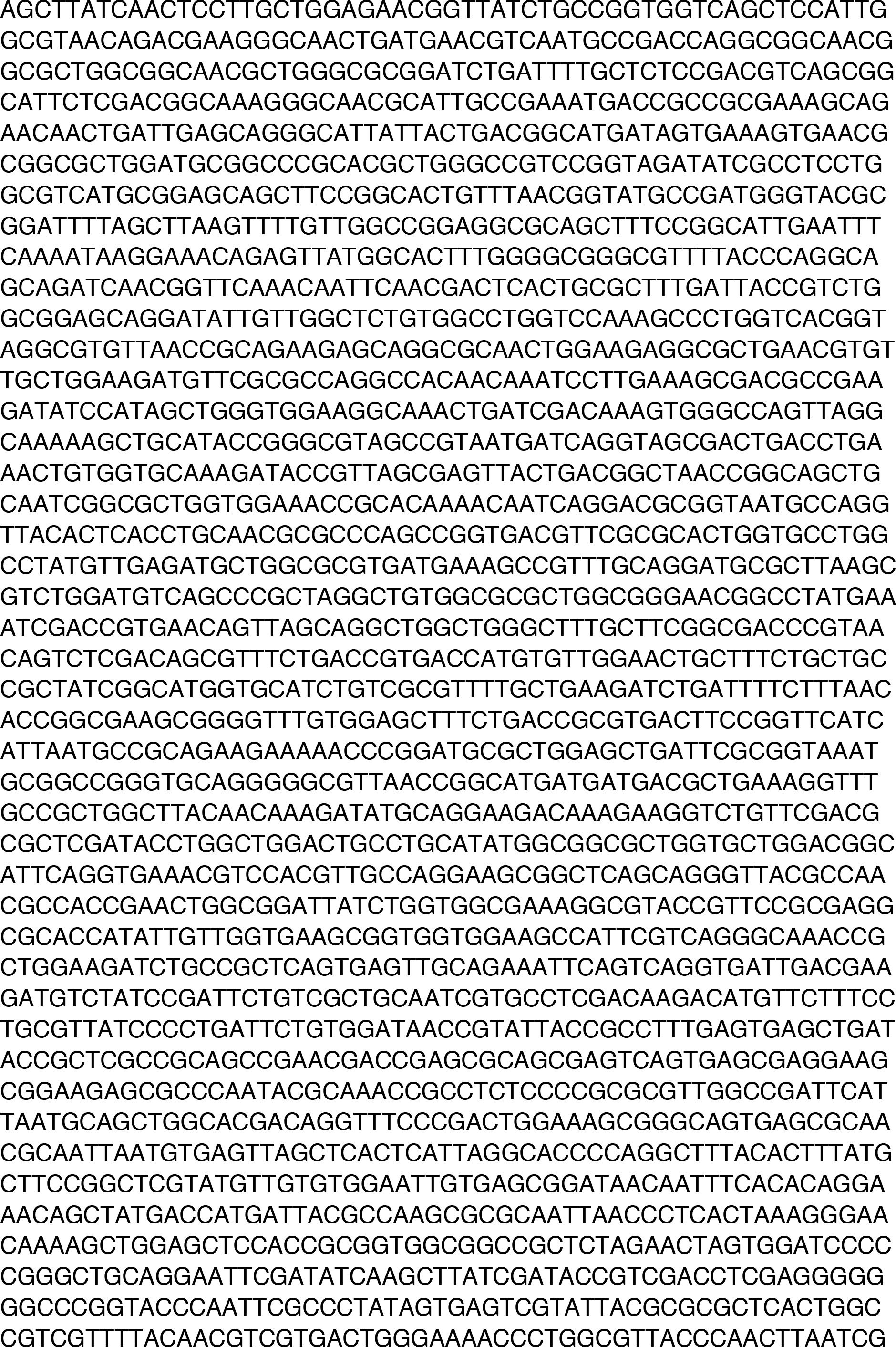
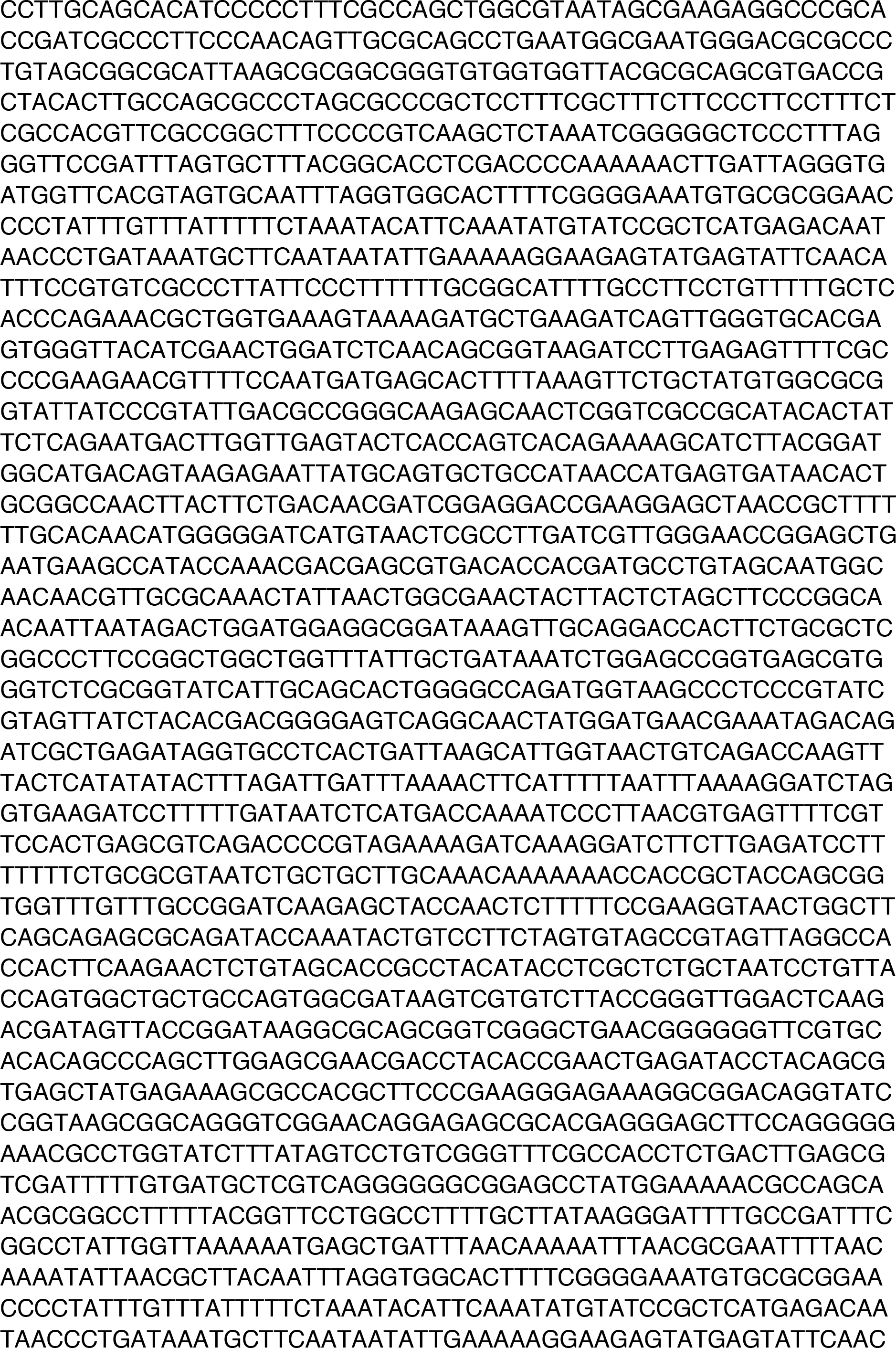
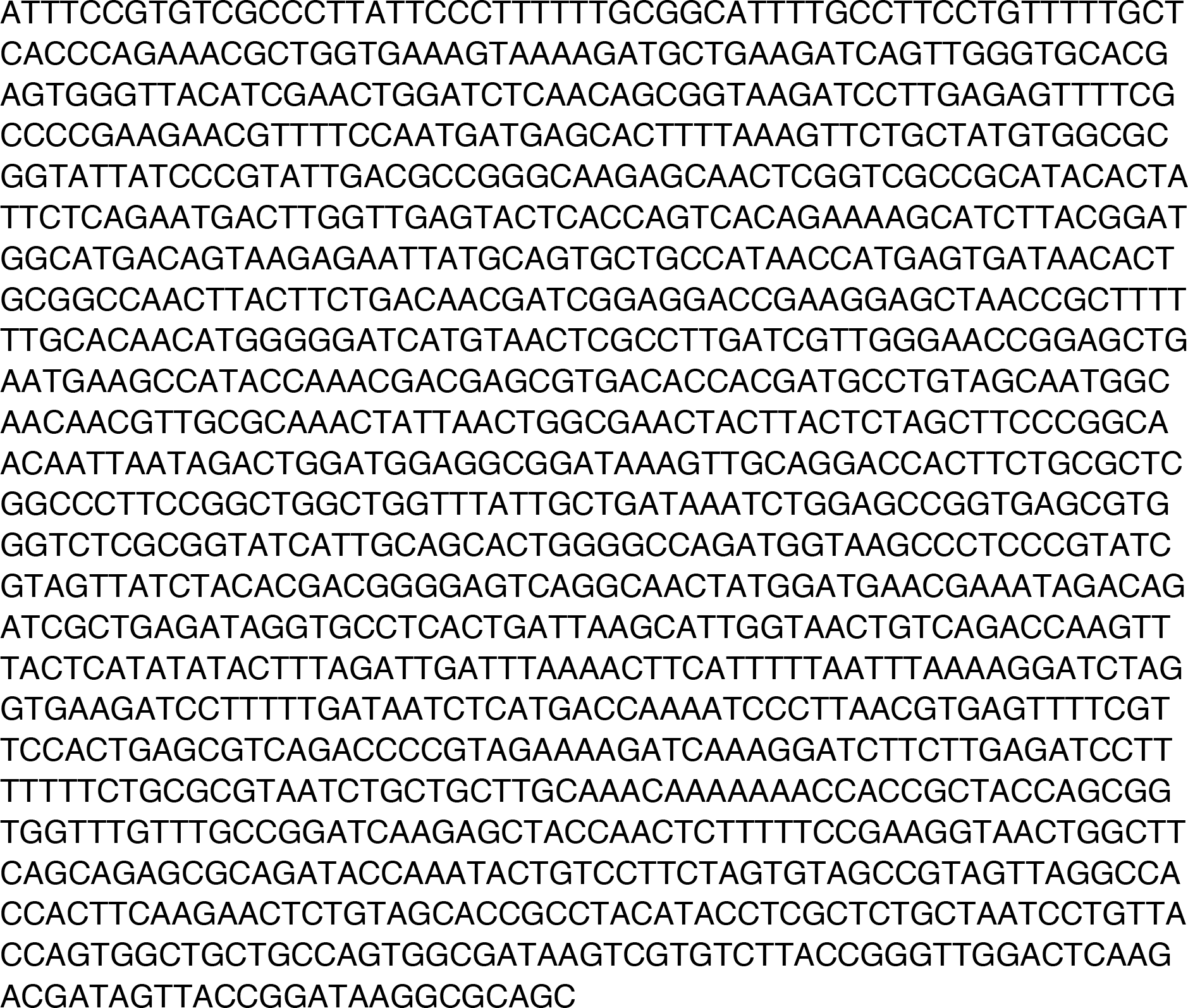
Plasmid Sequences

**Supplemental Table 2.**
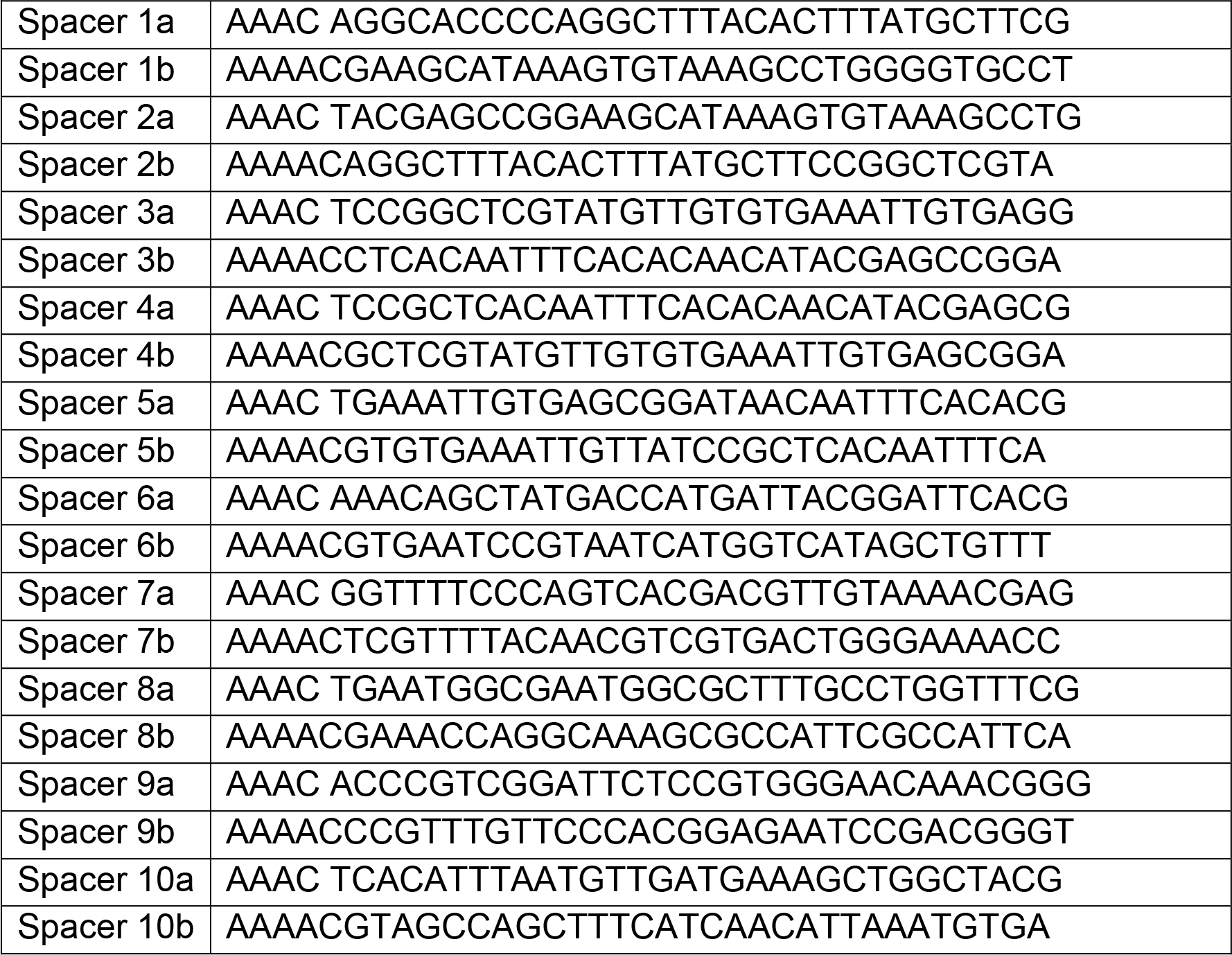
CRISPR spacer sequences Sequences of the oligonucleotides that were annealed together to make the oligoduplexes encoding the spacer sequences encoding the ten candidate gRNAs used in Figure 1.

**Supplemental Table 3.**
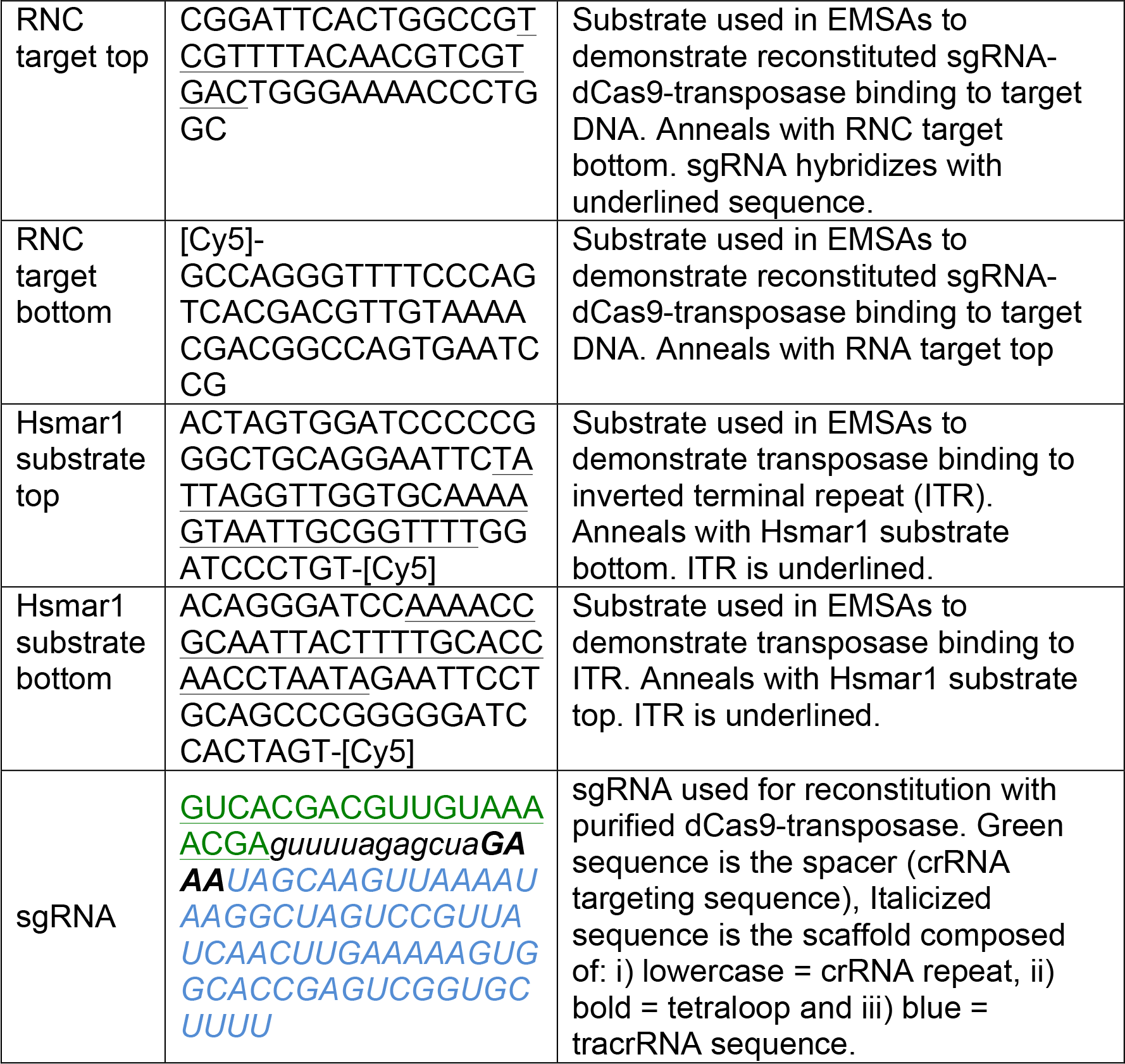
Oligonucleotide sequences

## REFERENCES

Ammar, I., Gogol-Doring, A., Miskey, C., Chen, W., Cathomen, T., Izsvak, Z., and Ivics, Z. (2012). Retargeting transposon insertions by the adeno-associated virus Rep protein. Nucleic acids research 40, 6693–6712.

Anders, C., Niewoehner, O., and Jinek, M. (2015). In Vitro Reconstitution and Crystallization of Cas9 Endonuclease Bound to a Guide RNA and a DNA Target. Methods in enzymology 558, 515–537.

Balciunas, D., Wangensteen, K.J., Wilber, A., Bell, J., Geurts, A., Sivasubbu, S., Wang, X., Hackett, P.B., Largaespada, D.A., McIvor, R.S., et al. (2006). Harnessing a high cargo-capacity transposon for genetic applications in vertebrates. PLoS genetics 2, e169.

Bikard, D., Jiang, W., Samai, P., Hochschild, A., Zhang, F., and Marraffini, L.A. (2013). Programmable repression and activation of bacterial gene expression using an engineered CRISPR-Cas system. Nucleic acids research 41, 7429–7437.

Blundell-Hunter, G., Tellier, M., and Chalmers, R. (2018). Transposase subunit architecture and its relationship to genome size and the rate of transposition in prokaryotes and eukaryotes. Nucleic Acids Research, gky794–gky794.

Branda, C.S., and Dymecki, S.M. (2004). Talking about a revolution: The impact of site-specific recombinases on genetic analyses in mice. Developmental cell 6, 7–28.

Chandrasegaran, S., and Smith, J. (1999). Chimeric restriction enzymes: what is next? Biological chemistry 380, 841–848.

Claeys Bouuaert, C., and Chalmers, R. (2010). Transposition of the human Hsmar1 transposon: rate-limiting steps and the importance of the flanking TA dinucleotide in second strand cleavage. Nucleic acids research 38, 190–202.

Claeys Bouuaert, C., and Chalmers, R. (2013). Hsmar1 Transposition Is Sensitive to the Topology of the Transposon Donor and the Target. PLoS ONE 8, e53690.

Claeys Bouuaert, C., and Chalmers, R. (2017). A single active site in the mariner transposase cleaves DNA strands of opposite polarity. Nucleic acids research 45, 11467–11478.

Claeys Bouuaert, C., Lipkow, K., Andrews, S.S., Liu, D., and Chalmers, R. (2013). The autoregulation of a eukaryotic DNA transposon. eLife 2, e00668.

Claeys Bouuaert, C., Liu, D., and Chalmers, R. (2011). A simple topological filter in a eukaryotic transposon as a mechanism to suppress genome instability. Molecular and cellular biology 31, 317–327.

Claeys Bouuaert, C., Walker, N., Liu, D., and Chalmers, R. (2014). Crosstalk between transposase subunits during cleavage of the mariner transposon. Nucleic acids research 42, 5799–5808.

Conte, E., Mende, L., Grainge, I., and Colloms, S.D. (2019). A Mini-ISY100 Transposon Delivery System Effective in γ Proteobacteria. Frontiers in microbiology 10, 280–280.

Crenes, G., Ivo, D., Herisson, J., Dion, S., Renault, S., Bigot, Y., and Petit, A. (2009). The bacterial Tn9 chloramphenicol resistance gene: an attractive DNA segment for Mos1 mariner insertions. Molecular genetics and genomics: MGG 281, 315–328.

Dawson, A., and Finnegan, D.J. (2003). Excision of the Drosophila mariner transposon Mos1. Comparison with bacterial transposition and V(D)J recombination. Molecular cell 11, 225–235.

Ding, S., Wu, X., Li, G., Han, M., Zhuang, Y., and Xu, T. (2005). Efficient transposition of the piggyBac (PB) transposon in mammalian cells and mice. Cell 122, 473–483.

Dornan, J., Grey, H., and Richardson, J.M. (2015). Structural role of the flanking DNA in mariner transposon excision. Nucleic acids research 43, 2424–2432.

Feng, X., Bednarz, A.L., and Colloms, S.D. (2010). Precise targeted integration by a chimaeric transposase zinc-finger fusion protein. Nucleic acids research 38, 1204–1216.

Gilbert, L.A., Horlbeck, M.A., Adamson, B., Villalta, J.E., Chen, Y., Whitehead, E.H., Guimaraes, C., Panning, B., Ploegh, H.L., Bassik, M.C., et al. (2014). Genome-Scale CRISPR-Mediated Control of Gene Repression and Activation. Cell 159, 647–661.

Grabundzija, I., Irgang, M., Mates, L., Belay, E., Matrai, J., Gogol-Doring, A., Kawakami, K., Chen, W., Ruiz, P., Chuah, M.K., et al. (2010). Comparative analysis of transposable element vector systems in human cells. Molecular therapy: the journal of the American Society of Gene Therapy 18, 1200–1209.

Ivics, Z., Katzer, A., Stuwe, E.E., Fiedler, D., Knespel, S., and Izsvak, Z. (2007). Targeted Sleeping Beauty transposition in human cells. Molecular therapy: the journal of the American Society of Gene Therapy 15, 1137–1144.

Izsvak, Z., Ivics, Z., and Plasterk, R.H. (2000). Sleeping Beauty, a wide host-range transposon vector for genetic transformation in vertebrates. Journal of molecular biology 302, 93–102.

Lamberg, A., Nieminen, S., Qiao, M., and Savilahti, H. (2002). Efficient insertion mutagenesis strategy for bacterial genomes involving electroporation of in vitro-assembled DNA transposition complexes of bacteriophage mu. Applied and environmental microbiology 68, 705–712.

Lampe, D.J., Churchill, M.E., and Robertson, H.M. (1996). A purified mariner transposase is sufficient to mediate transposition in vitro. The EMBO journal 15, 5470–5479.

Li, K., Wang, G., Andersen, T., Zhou, P., and Pu, W.T. (2014). Optimization of genome engineering approaches with the CRISPR/Cas9 system. PLoS One 9, e105779.

Liang, Q., Kong, J., Stalker, J., and Bradley, A. (2009). Chromosomal mobilization and reintegration of Sleeping Beauty and PiggyBac transposons. Genesis (New York, NY: 2000) 47, 404–408.

Lipkow, K., Buisine, N., and Chalmers, R. (2004). Promiscuous target interactions in the mariner transposon Himar1. The Journal of biological chemistry 279, 48569–48575.

Liu, D., and Chalmers, R. (2014). Hyperactive mariner transposons are created by mutations that disrupt allosterism and increase the rate of transposon end synapsis. Nucleic acids research 42, 2637–2645.

Liu, X., Homma, A., Sayadi, J., Yang, S., Ohashi, J., and Takumi, T. (2016). Sequence features associated with the cleavage efficiency of CRISPR/Cas9 system. Scientific reports 6, 19675.

Lohe, A.R., and Hartl, D.L. (1996). Reduced germline mobility of a mariner vector containing exogenous DNA: effect of size or site? Genetics 143, 1299–1306.

Luo, W., Galvan, D.L., Woodard, L.E., Dorset, D., Levy, S., and Wilson, M.H. (2017). Comparative analysis of chimeric ZFP-, TALE- and Cas9-piggyBac transposases for integration into a single locus in human cells. Nucleic acids research 45, 8411–8422.

Ma, H., Tu, L.C., and Naseri, A. (2016). CRISPR-Cas9 nuclear dynamics and target recognition in living cells. Journal of Cell Biology 214, 529–537.

Maragathavally, K.J., Kaminski, J.M., and Coates, C.J. (2006). Chimeric Mos1 and piggyBac transposases result in site-directed integration. FASEB journal: official publication of the Federation of American Societies for Experimental Biology 20, 1880–1882.

Mates, L., Chuah, M.K., Belay, E., Jerchow, B., Manoj, N., Acosta-Sanchez, A., Grzela, D.P., Schmitt, A., Becker, K., Matrai, J., et al. (2009). Molecular evolution of a novel hyperactive Sleeping Beauty transposase enables robust stable gene transfer in vertebrates. Nature genetics 41, 753–761.

Miller, J. (1972). Experiments in Molecular Genetics. Cold Spring Harbor Laboratory.

Miskey, C., Papp, B., Mates, L., Sinzelle, L., Keller, H., Izsvak, Z., and Ivics, Z. (2007). The ancient mariner sails again: transposition of the human Hsmar1 element by a reconstructed transposase and activities of the SETMAR protein on transposon ends. Molecular and cellular biology 27, 4589–4600.

Mitra, R., Fain-Thornton, J., and Craig, N.L. (2008). piggyBac can bypass DNA synthesis during cut and paste transposition. The EMBO journal 27, 1097–1109.

Munoz-Lopez, M., Siddique, A., Bischerour, J., Lorite, P., Chalmers, R., and Palomeque, T. (2008). Transposition of Mboumar-9: identification of a new naturally active mariner-family transposon. Journal of molecular biology 382, 567–572.

Nishiyama, K., Ichihashi, N., Kazuta, Y., and Yomo, T. (2015). Development of a reporter peptide that catalytically produces a fluorescent signal through alpha-complementation. Protein science: a publication of the Protein Society 24, 599–603.

Owens, J.B., Mauro, D., Stoytchev, I., Bhakta, M.S., Kim, M.S., Segal, D.J., and Moisyadi, S. (2013). Transcription activator like effector (TALE)-directed piggyBac transposition in human cells. Nucleic acids research 41, 9197–9207.

Robertson, H.M., and Zumpano, K.L. (1997). Molecular evolution of an ancient mariner transposon, Hsmar1, in the human genome. Gene 205, 203–217.

Song, A.J., and Palmiter, R.D. (2018). Detecting and Avoiding Problems When Using the Cre-lox System. Trends in genetics: TIG 34, 333–340.

Tellier, M., and Chalmers, R. (2018). Development of a papillation assay using constitutive promoters to find hyperactive transposases. bioRxiv, 423012.

Trubitsyna, M., Grey, H., Houston, D.R., Finnegan, D.J., and Richardson, J.M. (2015). Structural Basis for the Inverted Repeat Preferences of mariner Transposases. The Journal of biological chemistry 290, 13531–13540.

Trubitsyna, M., Michlewski, G., Finnegan, D.J., Elfick, A., Rosser, S.J., Richardson, J.M., and French, C.E. (2017). Use of mariner transposases for one-step delivery and integration of DNA in prokaryotes and eukaryotes by transfection. Nucleic acids research 45, e89.

Urnov, F.D., Miller, J.C., Lee, Y.L., Beausejour, C.M., Rock, J.M., Augustus, S., Jamieson, A.C., Porteus, M.H., Gregory, P.D., and Holmes, M.C. (2005). Highly efficient endogenous human gene correction using designed zinc-finger nucleases. Nature 435, 646–651.

Way, J.C., Davis, M.A., Morisato, D., Roberts, D.E., and Kleckner, N. (1984). New Tn10 derivatives for transposon mutagenesis and for construction of lacZ operon fusions by transposition. Gene 32, 369–379.

Yant, S.R., Huang, Y., Akache, B., and Kay, M.A. (2007). Site-directed transposon integration in human cells. Nucleic acids research 35, e50.

Zhang, F., Cong, L., Lodato, S., Kosuri, S., Church, G.M., and Arlotta, P. (2011). Efficient construction of sequence-specific TAL effectors for modulating mammalian transcription. Nature Biotechnology 29, 149.

